# Acquired FGFR and FGF alterations confer resistance to estrogen receptor (ER) targeted therapy in ER+ metastatic breast cancer

**DOI:** 10.1101/605436

**Authors:** Pingping Mao, Ofir Cohen, Kailey J. Kowalski, Justin G. Kusiel, Jorge E. Buendia-Buendia, Michael S. Cuoco, Pedro Exman, Seth A. Wander, Adrienne G. Waks, Utthara Nayar, Jon Chung, Samuel Freeman, Orit Rozenblatt-Rosen, Vincent A. Miller, Federica Piccioni, David E. Root, Aviv Regev, Eric P. Winer, Nancy U. Lin, Nikhil Wagle

**Affiliations:** Center for Cancer Precision Medicine, Dana-Farber Cancer Institute, Boston, MA; Department of Medical Oncology, Dana-Farber Cancer Institute, Boston, MA; Harvard Medical School, Boston, MA; Broad Institute of MIT and Harvard, Cambridge, MA; Department of Medicine, Brigham and Women’s Hospital, Boston, MA; Foundation Medicine, Inc. Cambridge, MA; Klarman Cell Observatory, Broad Institute of MIT and Harvard, Cambridge MA; Howard Hughes Medical Institute and Koch Institute of Integrative Cancer Research, Department of Biology, Massachusetts Institute of Technology, Cambridge, MA

**Keywords:** Drug resistance, endocrine therapy, ER+ breast cancer, FGF, FGFR

## Abstract

Beyond acquired mutations in the estrogen receptor (ER), mechanisms of resistance to ER-directed therapies in ER+ breast cancer have not been clearly defined. We conducted a genome-scale functional screen spanning 10,135 genes to investigate genes whose overexpression confer resistance to selective estrogen receptor degraders. Pathway analysis of candidate resistance genes demonstrated that the FGFR, ERBB, insulin receptor, and MAPK pathways represented key modalities of resistance. In parallel, we performed whole exome sequencing in paired pre-treatment and post-resistance biopsies from 60 patients with ER+ metastatic breast cancer who had developed resistance to ER-targeted therapy. The FGFR pathway was altered via *FGFR1, FGFR2,* or *FGF3* amplifications or *FGFR2* mutations in 24 (40%) of the post-resistance biopsies. In 12 of the 24 post-resistance tumors exhibiting FGFR/FGF alterations, these alterations were not detected in the corresponding pre-treatment tumors, suggesting that they were acquired or enriched under the selective pressure of ER-directed therapy. *In vitro* experiments in ER+ breast cancer cells confirmed that FGFR/FGF alterations led to fulvestrant resistance as well as cross-resistance to the CDK4/6 inhibitor palbociclib. RNA sequencing of resistant cell lines treated with different drug combinations demonstrated that FGFR/FGF induced resistance through ER reprogramming and activation of the MAPK pathway. The resistance phenotypes were reversed by FGFR inhibitors, a MEK inhibitor, and/or a SHP2 inhibitor, suggesting potential treatment strategies. The detection of targetable, clonally acquired genetic alterations in the FGFR pathway in metastatic tumor biopsies highlights the value of serial tumor testing to dissect mechanisms of resistance in human breast cancer and its potential application in directing clinical management.

## Introduction

Approximately 70% of breast cancers express the estrogen receptor (ER), and estrogen signaling drives breast cancer cell growth and progression [1]. Endocrine therapies are commonly used to treat ER+ breast cancer and work by reducing estrogen levels or targeting the estrogen receptor through functional inhibition or degradation. Although these endocrine therapies, including tamoxifen, aromatase inhibitors (AI), and the selective estrogen receptor degrader (SERD) fulvestrant have improved survival for ER+ breast cancer patients, within the metastatic setting resistance to endocrine therapies is nearly universal and remains a key challenge in reducing breast cancer morbidity and mortality [2].

Although various resistance mechanisms have been proposed for tamoxifen and aromatase inhibitor resistance, including loss or modification in ER expression (*ESR1* activating mutations and *ESR1* fusions) [3–7], and regulation of alternative signal transduction pathways (PI3K/AKT/mTOR and EGFR/ERBB2/MAPK) [8–10], mechanisms of resistance to SERDs remain understudied. Mechanisms of endocrine resistance identified in patients include acquired mutations in the estrogen receptor itself [4–7], acquired activating mutations in *ERBB2* (HER2) [11, 12], loss of function of *NF1* [13], and other alterations in MAPK pathway genes [14]. Additional mechanisms remain to be identified.

Gain-of-function screens have played a pivotal role in identification of resistance mechanisms to targeted therapies in various cancer types [15–17]. In breast cancer, several functional screen studies identified *IGF1R, KRAS* and *ESR1* as mechanisms of resistance to tamoxifen and/or estrogen deprivation [18–20]. However, genome-scale functional screens for SERD resistance have not been reported.

We conducted a genome-scale gain-of-function screen in ER+ breast cancer cells spanning 17,255 overexpressed lentiviral open reading frames (ORFs) to investigate genes whose overexpression was sufficient to confer resistance to the SERDs fulvestrant and GDC-0810 [21]. In parallel, we sought to identify endocrine resistance mechanisms of clinical significance through genomic profiling of paired pre-treatment and post-treatment tumor samples from 60 patients with ER+ metastatic breast cancer who developed resistance to endocrine therapy.

The intersection of top candidate resistance mechanisms from both approaches converged on the fibroblast growth factor receptor (FGFR) pathway. Here, we demonstrate that acquired FGFR/FGF alterations identified in patients with resistant metastatic breast cancer cause resistance to a variety of ER-directed therapies as well as to CDK4/6 inhibitors via MAPK pathway activation, and that this can be overcome by combination therapy targeting ER, the FGFR pathway, and/or the MAPK pathway.

## Results

### A Genome-Scale Gain-of-Function Screen for Resistance to Selective Estrogen Receptor Degraders

To identify the spectrum of genes whose overexpression confers resistance to SERDs *in vitro*, we expressed 17,255 human open reading frames (ORFs), corresponding to 10,135 distinct human genes, in ER+ T47D breast cancer cells in the presence of fulvestrant or GDC-0810. T47D cells were infected with the pooled lentiviral ORF library hORFeome [22]. Fulvestrant, GDC-0810, or vehicle control (DMSO) was added following infection and selection. ORF representation was assessed by sequencing after 21 days of drug exposure. Genes that confer drug resistance will be enriched under drug selection, indicated by a positive log fold change (LFC) for ORF representation before and after DMSO/drug selection.

Using a Z score >3 as a criterion to identify resistance candidates, we identified 64 genes (93 ORFs) that conferred resistance to fulvestrant and 57 genes (83 ORFs) that conferred resistance to GDC-0810 (Fig.1A and Supplemental Table.1). 37 genes (55 ORFs) conferred resistance to both drugs, a degree of overlap which was anticipated given the mechanistic similarities between fulvestrant and GDC-0810. The LFC and corresponding Z score for each ORF in fulvestrant and GDC-0810 treatment arms were highly correlated, with a correlation coefficient of 0.77 (Fig.1A).

**Figure 1.**
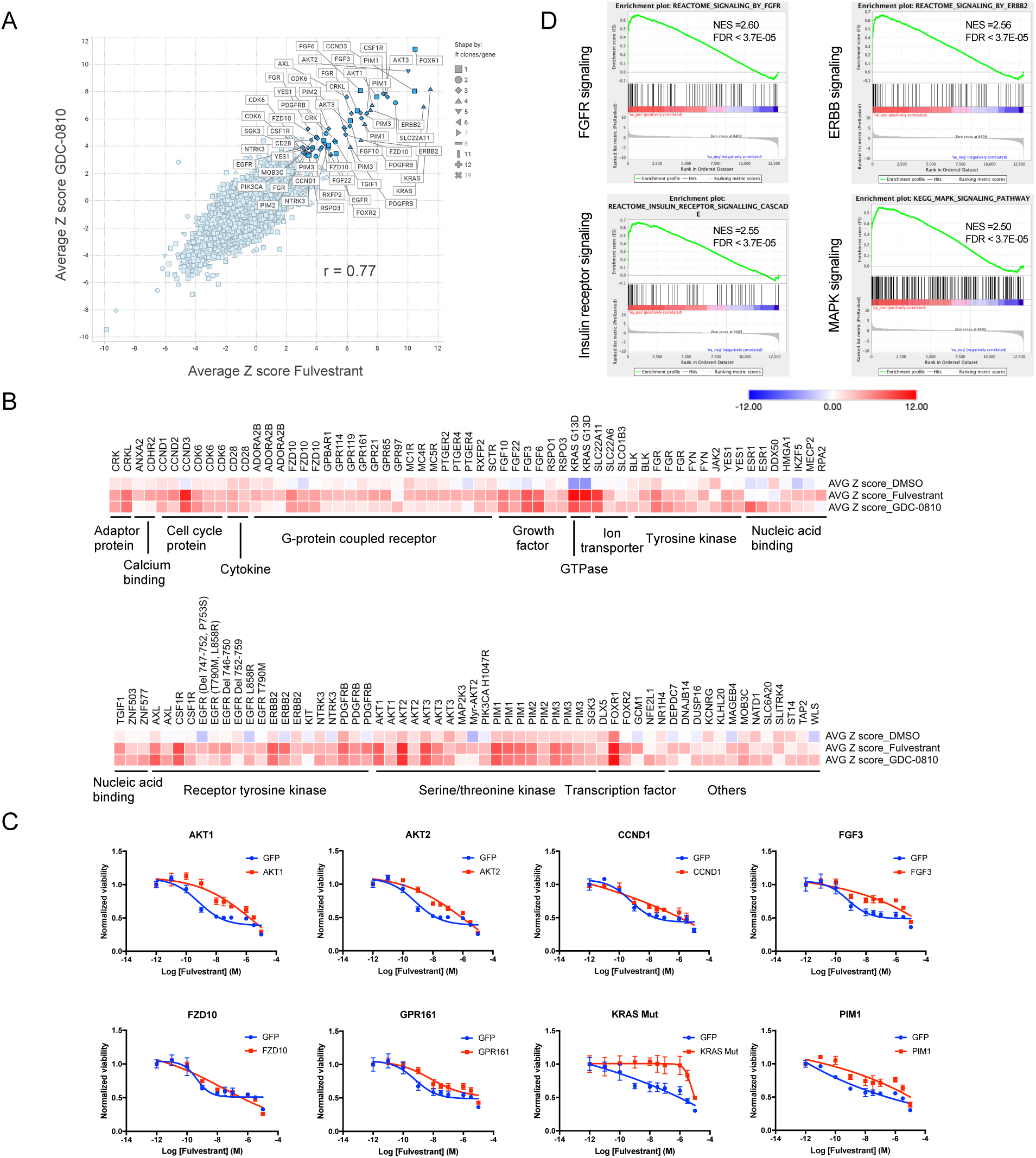
A genome-scale gain-of-function screen identified resistance genes to fulvestrant and GDC-0810. 17,255 human open reading frames (ORFs), corresponding to 10,135 distinct genes, were expressed in ER+ T47D breast cancer cells in the presence of fulvestrant or GDC-0810. Fulvestrant, GDC-0810, or vehicle control (DMSO) was added following infection and selection. ORF representation was assessed by sequencing after 21 days of drug exposure. Genes that confer drug resistance will be enriched under drug selection, indicated by a positive log fold change (LFC) for ORF representation before and after DMSO/drug selection. A, The average Z score for LFC of each ORF was plotted for both the fulvestrant (X-axis) and GDC-0810 (Y-axis) arms. The average Z score was calculated from three replicates in each condition. The ORFs with a Z score > 3 in both drug arms are highlighted and labeled with gene ID. The shape of each data point represents the total number of ORFs for that gene in the library. B, Heatmap of top ORF hits with a Z score > 3 in fulvestrant or GDC-0810 arm. The Z score in the DMSO arm is also presented. ORF hits are grouped by their molecular function according to Uniprot annotation. Information on the complete list of ORFs can be found in Supplemental Table.1. C, Individual ORFs were overexpressed in T47D cells and validated to confer resistance to fulvestrant by drug response curves. *KRAS* G12D ORF was used for overexpression in T47D cells while other selected ORFs are wildtype. Cell viability was measured by CellTiter-Glo and all the data points were normalized to growth under DMSO condition. Results shown are ± SD and representative of three independent experiments. D, Gene set enrichment analysis was performed for the gene list ranked by LFC in the fulvestrant arm. For genes with multiple ORFs, the ORF with highest LFC was selected. 1000 permutations were performed for the analysis. NES, normalized enrichment score. The full list of nominated pathways is shown in Supplemental Table.2.

To confirm these results, we conducted a secondary screen using a smaller pooled library consisting of 570 ORFs to validate candidates nominated by the primary screen. The secondary screen was performed in both T47D and MCF7 cell lines with a similar screen process as the primary screen. Top resistance genes found in the primary screen were again enriched in the secondary screen, including *FGF* genes*, FOXR1, AKT* genes*, PIM* genes and several *GPCR* genes (Supplemental Data Fig.S1). Many top ranked resistance genes (*CSF1R, FGF3, FGF6, FOXR1* and *PIM2*) were shared between T47D and MCF7 cells (Supplemental Data Fig.S2). However, distinct resistance genes were also observed in each cell line, suggesting some resistance mechanisms may be cell context-dependent.

Functional categories of candidate resistance genes include serine/threonine kinases (*PIK3CA, AKT1/2/3, PIM1/2/3*), receptor tyrosine kinases (*EGFR, ERBB2, PDGFRB*), growth factors (*FGF3/6/10/22*), cell cycle regulatory proteins (*CCND1, CCND2, CCND3, CDK6*) and G-protein coupled receptors (*GPCR*) (Fig.1B). As further validation, we overexpressed 13 ORFs belonging to these categories individually in T47D cells and they conferred resistance to fulvestrant to various degrees (Fig.1C and Supplemental Data Fig.S3A). Most of the 13 ORFs also conferred resistance to GDC-0810 (Supplemental Data Fig.S3B).

Gene Set Enrichment Analysis (GSEA) of the candidate resistance genes demonstrated enrichment in 4 functional pathways: FGFR signaling, ERBB signaling, insulin receptor signaling, and the MAPK pathway (Fig.1D, Supplemental Data Fig.S4 and Supplemental Table.2). Consistent with this, we recently demonstrated that acquired *ERBB2* activating mutations activate the MAPK pathway and cause endocrine resistance in patients with ER+ metastatic breast cancer [11]. Several recent studies have also shown that alterations in MAPK pathway genes are enriched in endocrine-resistant tumors [13, 14]. We sought to further examine the role of FGFR and FGF genes in resistance to SERDs in MBC.

### Identification of acquired FGFR and FGF alterations in metastatic biopsies from patients with resistant ER+ MBC

To examine the potential role of FGFR and FGF alterations in the development of endocrine resistance clinically, we analyzed whole exome sequencing (WES) data from paired pre-treatment and post-treatment metastatic tumor biopsies or cell free DNA from 60 patients with ER+ metastatic breast cancer who had received at least one endocrine therapy (tamoxifen, AI, SERDs) for more than 120 days between the two biopsies [23].

Amongst the 60 post-treatment samples, we found *FGFR1* amplifications in 15% (9/60), *FGFR2* amplifications in 5% (3/60), *FGFR2* activating mutations in 3.3% (2/60), and *FGF3* amplifications in 28.3% (17/60) – for a total of 40% (24/60) of the cohort with at least one alteration in one of these four genes (Fig.2A). Overall, the prevalence of *FGFR1, FGFR2*, and *FGF3* alterations in the resistant metastatic setting seen here is increased compared to what was observed in previously published cohorts of primary ER+ breast cancer, such as The Cancer Genome Atlas (TCGA) [24] (Supplemental Table.3). The incidence of *FGFR2* alterations (6.7%), in particular, is markedly increased compared to primary treatment-naive breast cancer, in which the incidence is less than 2% in TCGA (Supplemental Table.3).

**Figure 2.**
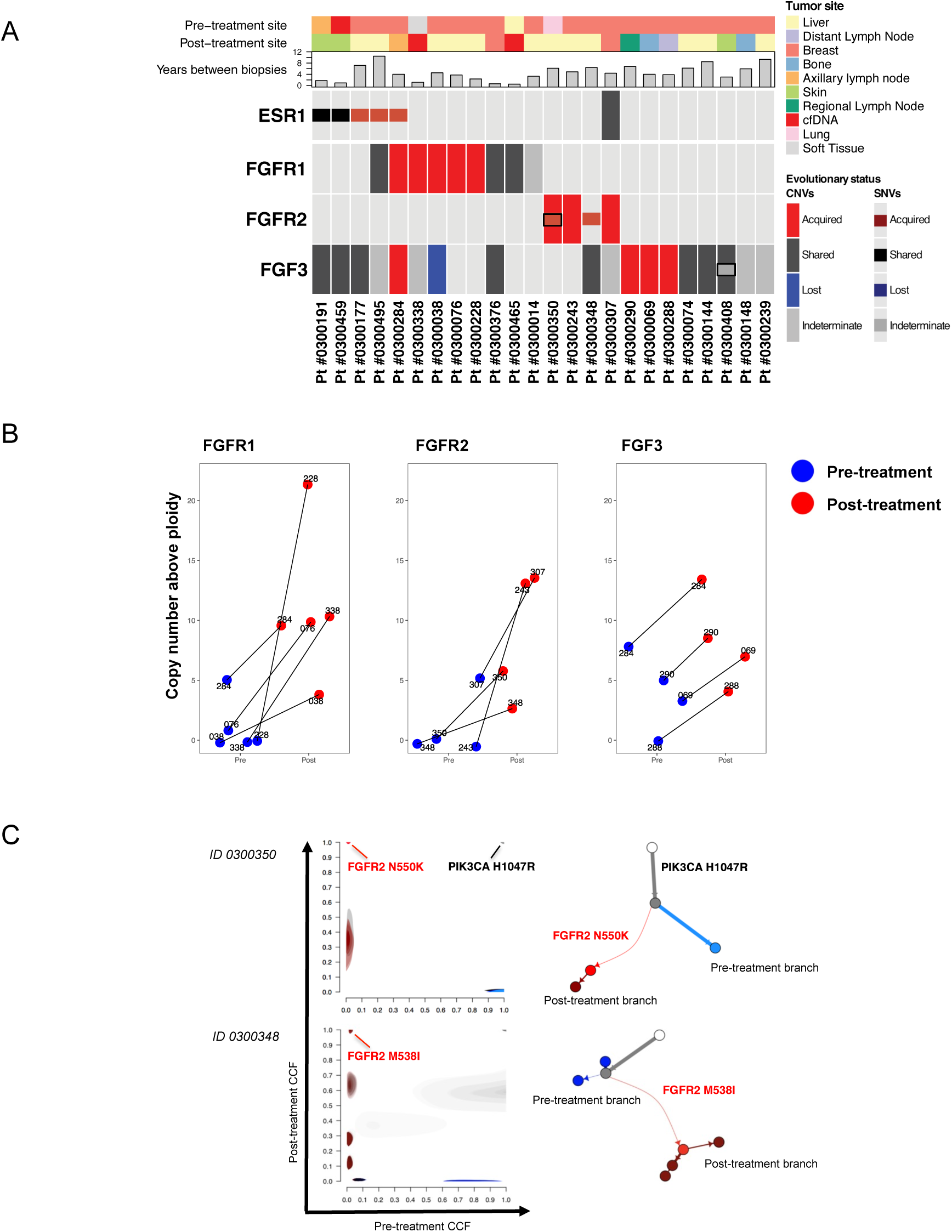
Identification of acquired FGFR and FGF alterations in metastatic biopsies from patients with resistant ER+ MBC. A, Evolutionary status of *ESR1, FGFR1, FGFR2*, and *FGF3* alterations is presented by comparing the pre-treatment and post-treatment mutational status for each patient (red = acquired, blue = lost, black = shared, grey = indeterminate). Clinical and pathology tracks depict the site of biopsy for both matched samples, and the duration between the pre-treatment and post-treatment biopsies. B, Copy number alterations for *FGFR1, FGFR2* and *FGF3* in pre- and post-treatment tumor samples are shown with copy number above ploidy (CNAP) depicted to illustrate the magnitude of the acquired amplification in each case. To better measure segment-specific copy number, we subtracted the genome ploidy for each sample to compute CNAP. The purity and ploidy for tumor samples are shown in Supplemental Table.4. C, Clonal evolution analysis showing the overall clonal structure and acquisition for FGFR2 mutations observed in two patients. In the pre-treatment biopsies, *FGFR2* M538I (*ID 0300348*) and *FGFR2* N550K (*ID 0300350*) were with cancer cell fraction (CCF) of 2% (single read) and 0% (unobserved), respectively, while being observed as clonal mutations in the post-treatment sample with a CCF of 1. The phylogenetic relationships among clones are reconstructed for each patient starting from the normal cell (white circle) connected to the ancestral cancer cells (grey trunk). The phylogenetic divergence to the pre-treatment clones (and subclones) is depicted with blue edges, and phylogenetic divergence to the metastatic clones (and subclones) is in red. Selected mutations in cancer genes are marked on the corresponding branches of the cancer phylogeny.

To determine if this enrichment of FGFR/FGF alterations in the metastatic setting was due to acquisition/selection under the selective pressure of endocrine therapy, we compared the WES from the paired pre-treatment and post-treatment samples for the 24 patients that exhibited FGFR/FGF alterations in their post-treatment samples. These 24 pairs of samples included 23 tumor biopsies and one cell-free DNA sample at the pre-treatment timepoint, and 22 tumor biopsies and two cell-free DNA samples at the post-treatment timepoint. We performed an evolutionary analysis to evaluate clonal structure and dynamics, including changes in mutations and copy number. The evolutionary inference and clonal dynamics of mutations was based on changes in the estimated fraction of tumor cells harboring each genomic alteration (the cancer cell fraction, CCF) as previously shown for acquired HER2 mutations [11]. The evolutionary inference of copy number changes was based on measuring differences in copy number amplitudes between pre-treatment and post-treatment samples, while accounting for differences in cancer cell fraction (“purity”) in the sample and correcting for differences in ploidy. The resultant purity-corrected values provide an estimate of “copy number above ploidy” (CNAP) (see Methods).

For this analysis, we define “acquired” alterations as alterations with higher representation in the post-treatment sample as compared to the pre-treatment sample. For single nucleotide variants (SNVs), this means that the mutation had a substantially higher CCF in the post-treatment sample compared to the pre-treatment sample (including lack of detection in the pre-treatment sample despite having sufficient power to detect the mutation). For copy number changes, this means that there was a substantial increase in the overall copy-number in the post-treatment sample. Although we use the term “acquired”, we recognize that when the mutation is not detected in the pre-treatment sample, we cannot distinguish between pre-existing alteration that was selected for and clonally enriched versus *de novo* alterations that developed during the treatment.

In 12 of the 24 patients with FGFR or FGF alterations (50%), the alterations were acquired in the post-treatment sample as compared to the pre-treatment sample (Fig. 2A, marked in red). Five out of nine *FGFR1* amplifications were acquired (55.6%), while all four *FGFR2* alterations were acquired (100%), including one patient (Pt 0300350) with acquisition of both an *FGFR2* mutation and amplification. *FGF3* amplifications were acquired in 4 of 17 tumors (23.5%), including one case in which an *FGFR1* amplification was co-acquired. The concurrent acquisition may suggest that the evolutionary selection of both the ligand and receptor provided additional fitness in this tumor. Among the other 12 patients, the alterations in eight patients were shared in both pre-treatment and post-treatment samples (Fig 2A, marked in black), and evolutionary status of alterations in the remaining four patients was inconclusive (Fig 2A, marked in grey). The increase in copy number (corrected for tumor purity and ploidy, Supplemental Table.4) from pre-treatment to post-treatment for *FGFR1, FGFR2*, and the *FGF3* amplicon in all 12 patients is depicted in Fig.2B.

Two of the acquired alterations found in these 12 patients were SNVs in the *FGFR2* gene: M538I (chr10:123258070C>T, GRCh37, also denoted as M537I, depending on the isoform) and N550K (chr10:123258034A>T, GRCh37, also denoted as N549K, depending on the isoform). N550K is the most common FGFR2 mutation in breast cancer while M538I was previously identified in lung cancer but has not yet been characterized in breast cancer [25]. Figure 2C illustrates the change in the estimated fraction of tumor cells harboring each genomic alteration (CCF) from the pre-treatment biopsy to the resistant biopsy. In both patients, the *FGFR2* mutations were either not detected in the primary tumor, despite sufficient power to detect mutations at this locus (N550K in Pt 0300350) or detected by a single read, inferred in a small fraction (CCF of 2%) of the pre-treatment tumor (M538I in Pt 0300348). In both patients the activating *FGFR2* mutations in the post-treatment biopsies were clonally acquired (CCF of 100%). Pt 0300350 was also found to have an acquired *FGFR2* amplification while Pt 0300348 was found to have gained low-level amplification in *FGFR2* post treatment (Fig.2B).

Notably, the acquired alterations in *FGFR1, FGFR2*, and *FGF3* were largely mutually exclusive with acquired *ESR1* mutations. *ESR1* mutations are the most common mechanism described for acquired endocrine resistance [26]. Although the overall rate of acquired *ESR1* mutation in this cohort is 22% (13/60), among the 12 cases of acquired FGFR and FGF alterations, only one patient also has an acquired ESR1 mutation (Figure 2A). Similarly, only one of these 12 patients had acquired a HER2 mutation (which was an alteration of unknown significance), suggesting that these are also mutually exclusive mechanisms of resistance.

Although we highlight *FGF3* as the key gene in the amplicon (given the results of the gain-of-function screen), *FGF3* resides in genomic proximity to *FGF4, FGF19* and *CCND1* and these four genes are often co-amplified. Here, in 3 out of the 4 cases with acquired *FGF3* amplification, *FGF3* copies were gained without co-acquisition of *CCND1* amplification, suggesting that this acquisition can occur as an independent genomic event. Similarly, 2 out of the 4 cases with acquired *FGF3* did not have co-acquisition of *FGF19* amplification. In all 4 of the cases with *FGF3* amplification*, FGF4* was also co-amplified. The relationship of acquisitions of *FGF3, FGF4, FGF19*, and *CCND1* are depicted in Supplemental Data Fig.S5 and further details about these amplicons are described in Supplemental Methods. Further exploration of the full genomic contexts and other concurrent genetic alterations for the 12 patients with acquired FGFR/FGF alterations are shown in Supplemental Data Fig.S6 and Supplemental Table.5-6.

Figure 3 depicts clinical vignettes for six of the patients with acquired *FGFR1, FGFR2*, and/or *FGF3* alterations in their post-treatment biopsies. All patients were treated with ER-directed therapy before acquiring FGF or FGFR alterations, including tamoxifen (3 patients), AIs (6 patients), and fulvestrant (3 patients). Vignettes for the other six patients with acquired *FGFR1, FGFR2*, and/or *FGF3* alterations in their post-treatment biopsies are shown in Supplemental Data Fig.S7. Detailed clinicopathological features and therapy details for all 12 patients are found in Supplemental Table.7.

**Figure 3.**
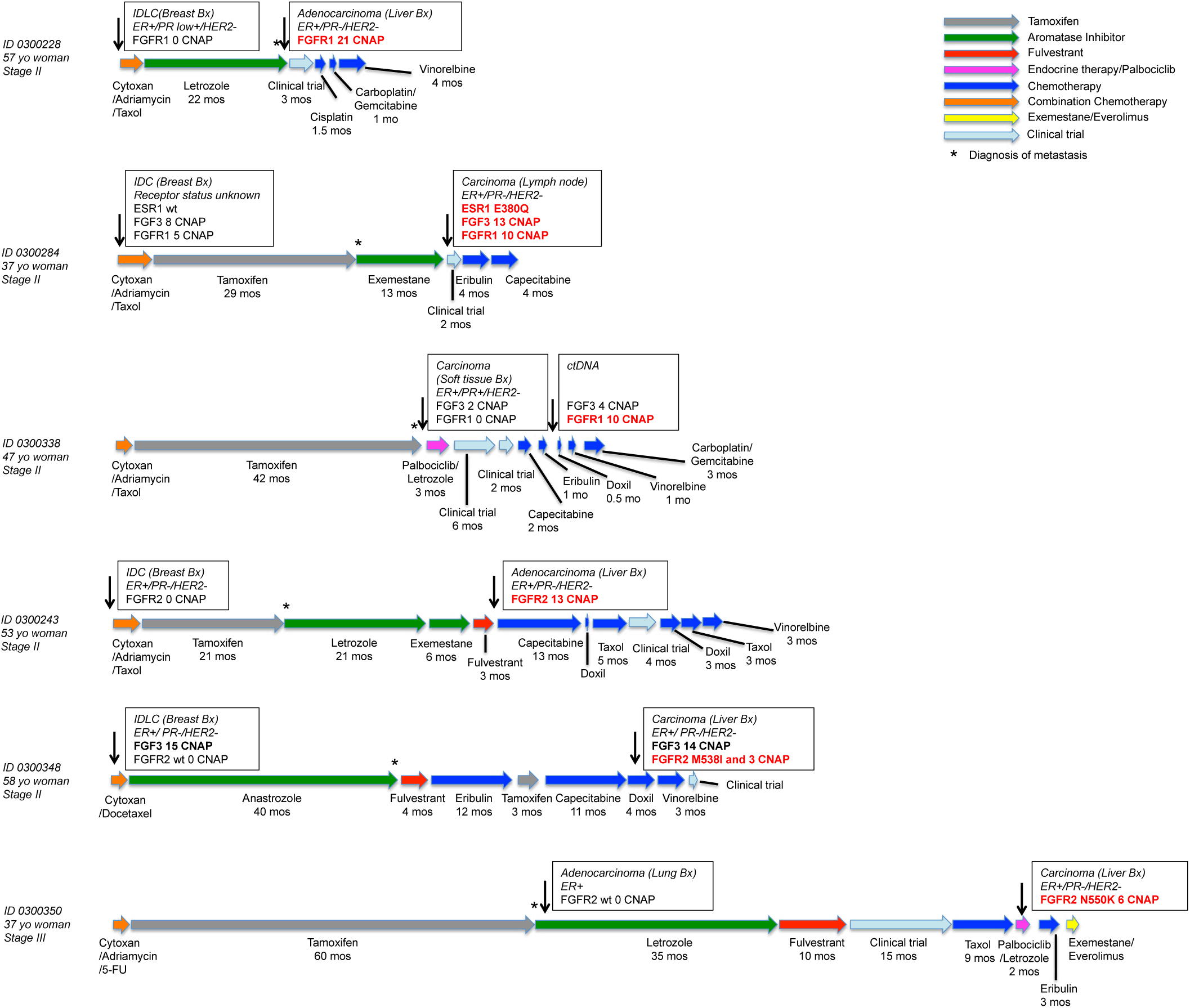
Clinical vignettes of patients who acquired FGFR/FGF alterations following endocrine therapy. The clinical vignettes for selected patients with acquired alterations in *FGFR1, FGFR2*, and/or *FGF3* illustrate detailed information on age and stage of disease at diagnosis, therapies patients received, duration of response to each therapy, and time of biopsies collected during the clinical course. For each biopsy, available information on biopsy type, tissue site, receptor status and selected genomic alterations detected by whole exome sequencing is shown. In each case, the asterisk indicates the time that metastatic disease was diagnosed. The complete clinicopathologic information for each patient is provided in Supplemental Table.7. IDC: invasive ductal carcinoma, IDLC: invasive ductal-lobular carcinoma; CNAP: copy number above ploidy; yo: years old; Bx: biopsy; PR: progesterone receptor; wt: wildtype.

In addition to these 12 patients in our cohort, we identified several additional patients with acquired *FGFR1* and *FGFR2* activating mutations following the development of resistance to endocrine therapy (Supplemental Data Fig.S8 and Supplemental Table.8). FM patient 1 acquired a clonal *FGFR1* N546K mutation (a known activating mutation paralogous to *FGFR2* N550K) following treatment with an AI. FM patient 2 acquired a subclonal *FGFR2* N550K mutation after treatment with tamoxifen, AI and fulvestrant. FM patient 3 acquired a subclonal *FGFR2* K660N mutation, another activating mutation in the kinase domain [27], after treatment with tamoxifen.

In summary, we observed acquired alterations in *FGFR1, FGFR2*, or *FGF3* in 20% (12/60) of patients with endocrine resistant ER+ MBC – comparable to the known frequency of acquired mutations in *ESR1* – highlighting the important role of the FGFR pathway in acquired resistance to ER-directed therapies.

### Active FGFR signaling leads to resistance to SERDs through activation of the MAP kinase pathway

To further investigate how FGFR/FGF genes may confer resistance to ER-directed therapy, we treated T47D cells with FGF3, FGF6, FGF10 or FGF22 ligand. Each of these ligands resulted in resistance to fulvestrant (Fig.4A). This effect was reversed by PD173074, a pan-FGFR inhibitor (Fig.4A). The addition of these FGF ligands enhanced phosphorylation of ERK and AKT, which was reversed by PD173074 (Fig.4B). The effect of FGF ligands on downstream effectors was enhanced when FGFR1 was simultaneously overexpressed in T47D, which has relatively low expression of FGFR1 [28] (Supplemental Data Fig.S9). FGF3, FGF6, FGF10 and FGF22 also reduced fulvestrant sensitivity in MCF7 cells (Supplemental Data Fig. S10 A-B). Similar results have been shown previously for FGF2, which was reported to activate MEK-ERK to drive fulvestrant resistance in ER+ breast cancer cells [29].

**Figure 4.**
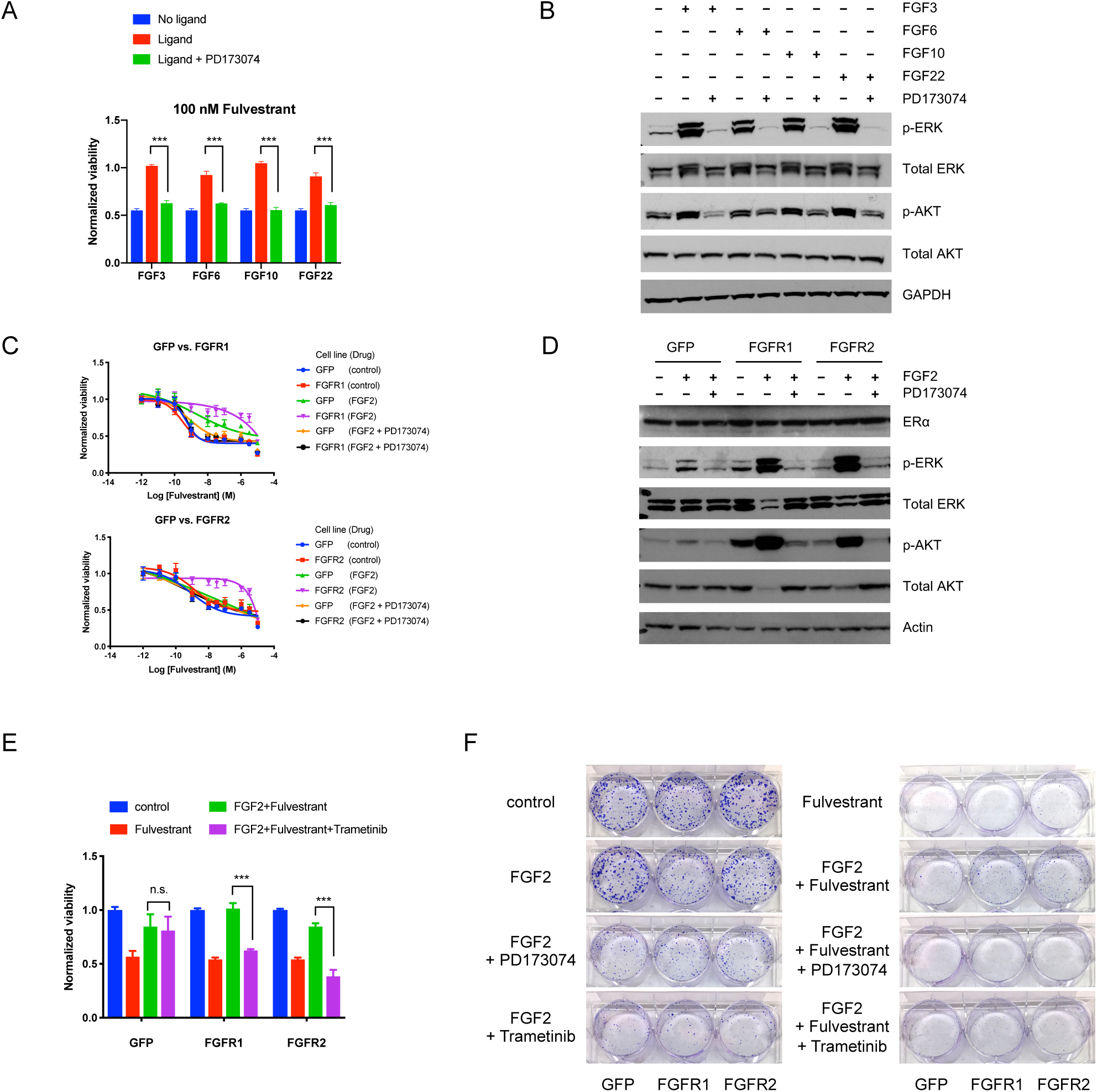
Active FGFR signaling leads to resistance to SERDs through activation of MAPK pathway. A, FGF ligands lead to resistance to fulvestrant, which was blocked by FGFR inhibitor PD173074. Recombinant FGF ligands were added into media every three days at the concentration of 100 ng/mL with or without 1 μM PD173014. T47D cells were treated with heparin (1 μg/mL) that facilitates the binding between FGF ligand and receptor, and sensitivity to 100 nM fulvestrant over six days was normalized to DMSO control. *** p-value < 0.001. Student t-test was performed for pair-wise comparisons. Results shown are ± SD and representative of three independent experiments. B, FGF ligands increased ERK and AKT phosphorylation, which was blocked by PD173074. T47D cells were treated as indicated for one hour before protein harvest for western blot. Results shown are representative of two independent experiments. C, FGFR1 or FGFR2 overexpression leads to resistance to fulvestrant, which was blocked by PD173074. GFP, FGFR1 or FGFR2 was overexpressed in T47D cells to establish stable T47D_GFP, T47D_FGFR1 and T47D_FGFR2 cells. The fulvestrant sensitivity of various cell lines were determined in the presence or absence of 10 ng/mL FGF2 and 1 μM PD173074 over six days of drug treatment. Results shown are ± SD and representative of three independent experiments. D, FGFR1 and FGFR2 induced phosphorylation of ERK and AKT in the presence of FGF2, which was blocked by PD173074. Results shown are representative of two independent experiments. Cells were treated with indicated conditions for one hour before protein harvest. E and F, Trametinib abrogated the resistance to fulvestrant conferred by FGFR1 or FGFR2. Cells were treated with different conditions as indicated: 10 ng/mL FGF2; 100 nM fulvestrant; 500 nM trametinib. CellTiter-Glo assay was performed to measure cell viability after six days for all dose response curves (E), data are ± SD. 2,000 cells were plated on Day 1 and treated on Day 2, and colony formation assay was performed after three weeks of drug treatment (F). Results shown are representative of three and two independent experiments, respectively. * p-value < 0.05, ** p-value < 0.01, *** p-value < 0.001, n.s. not significant. Student t-test was performed for pair-wise comparisons.

We next overexpressed FGFR1, FGFR2, or GFP in T47D cells through lentiviral transduction and examined the impact on susceptibility to SERDs. Overexpression of FGFR1 or FGFR2 alone did not affect sensitivity to fulvestrant or GDC-0810. However, with the addition of FGF2 ligand, both FGFR1 and FGFR2 rendered cells highly resistant to the two SERDs (Fig. 4C and Supplemental Data Fig. S11). In comparison, FGF2 ligand alone reduced sensitivity to SERDs in control cells expressing GFP to a much lesser extent than in the FGFR1 or FGFR2 expressing cells, suggesting the potent resistance phenotype requires both FGF ligand and receptor. This requirement for the presence of both FGF ligand and receptor for maximal resistance phenotype may also explain why only FGFs but not FGFR1 or FGFR2 scored in the resistance screen (Fig.1 A-B). The resistance phenotype resulting from FGFR1 and FGFR2 overexpression was completely reversed by the addition of PD173074 (Fig. 4C). Similar results were obtained in MCF7 cells (Supplemental Data Fig.S12 A-B).

FGFR1 and FGFR2 overexpression (in the presence of FGF2 ligand) induced more potent phosphorylation of AKT and ERK than the GFP control (Fig.4D). These results are consistent with previous findings that FGFR1 activation led to MAPK activation and fulvestrant resistance [28]. Activation of downstream effectors p-ERK and p-AKT by FGFR1/2 overexpression was reversed to baseline levels with PD173074 (Fig. 4D and Supplemental Data Fig. S12C). Examination of a larger number of kinases using kinase antibody arrays demonstrated that AKT, ERK and RSK (downstream effector of ERK) were the only kinases of those tested to exhibit increased phosphorylation following FGFR1/2 overexpression and FGF2 stimulation (Supplemental Data Fig.S13).

Collectively, these findings suggest that FGFR1 and FGFR2 cause SERD resistance through the activation of MAPK and/or PI3K/AKT pathways.

We examined the sensitivity of cells overexpressing FGFR1 or FGFR2 to several inhibitors of downstream effectors: the MEK inhibitor trametinib, the AKT inhibitor AZD5363, and the mTOR inhibitor everolimus. FGFR1 or FGFR2 overexpression in the presence of FGF2 led to hypersensitivity to trametinib (Supplemental Data Fig.S14A), but reduced sensitivity to AKT and mTOR inhibitors (Supplemental Data Fig.S14A).

We attempted to reverse FGFR-induced resistance to fulvestrant by inhibiting the MAPK pathway. Treatment of FGFR1 overexpressing cells with trametinib partially resensitized cells to fulvestrant, and treatment of FGFR2 overexpressing cells with trametinib fully resensitized the cells to fulvestrant (Fig.4E-F). Treatment with the mTOR inhibitor everolimus also partially reversed resistance conferred by FGFR1 or FGFR2 overexpression (Supplemental Data Fig.S14B). Colony formation assays produced similar results (Supplemental Data Fig.S14C).

Together, these results suggest that the MAPK pathway is the primary downstream effector of FGFR activation resulting in endocrine resistance. This is consistent with the pathway analysis of resistance genes in our initial overexpression screen, which demonstrated enrichment in MAPK pathway genes (Figure 1D), as well as our prior findings of MAPK activation through acquired HER2 mutations in endocrine resistance—adding further support to the idea that the MAPK pathway may be a common node of endocrine resistance [11, 14].

### FGFR activation confers cross-resistance to CDK4/6 inhibitors

Since the combination of endocrine therapy and CDK4/6 inhibitors is now a standard of care treatment for patients with ER+ metastatic breast cancer, we also examined the effect of FGFR signaling on sensitivity to the combination of fulvestrant and the CDK4/6 inhibitor palbociclib. In T47D cells, FGFR1 and FGFR2 overexpression in the presence of FGF2 also conferred resistance to combination treatment of fulvestrant and palbociclib (Fig. 5A). The resistance phenotype was again abrogated by PD173074 (Fig. 5A). Resistance to fulvestrant and palbociclib was also partially reversed by trametinib (Fig. 5B and Supplemental Data Fig. S14D), further providing the support for the role of MAPK pathway activation in FGFR-mediated drug resistance. The reversal of resistance by trametinib was accompanied by reduced ERK phosphorylation (Fig. 5C). Similar results were achieved in MCF7 cells, although everolimus was more effective than trametinib in reversing the resistance phenotype by FGFR1 or FGFR2 overexpression in this cell line (Supplemental Data Fig.S15 and Fig.S16).

**Figure 5.**
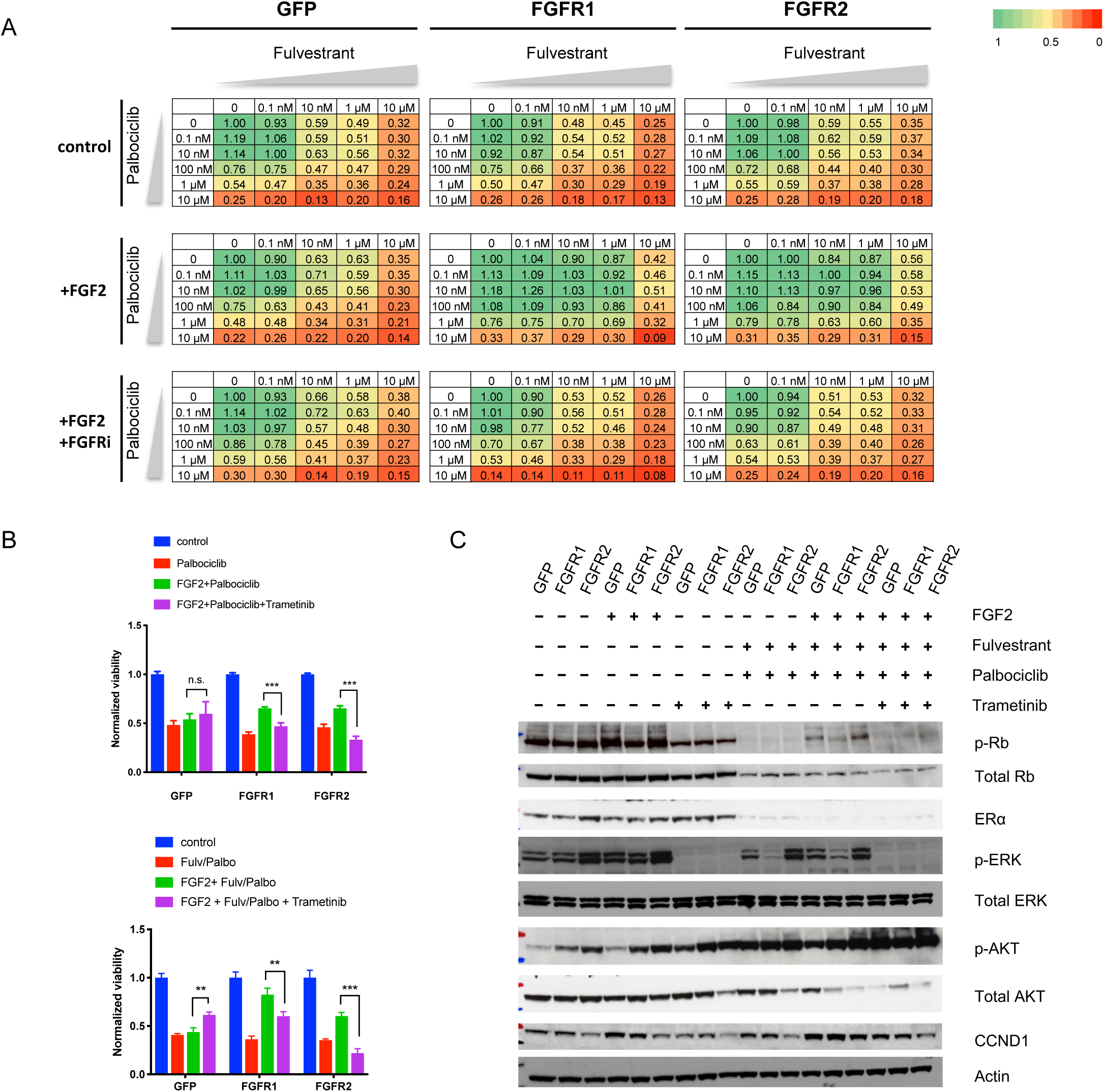
Active FGFR signaling leads to resistance to the combination of SERDs and CDK4/6 inhibitors. A, T47D_GFP, T47D_FGFR1 and T47D_FGFR2 cells were treated with the combination of fulvestrant and palbociclib at various doses for six days. The percentage of cell survival shown was normalized to DMSO control for each condition. The combination treatment was carried out under three conditions: 1, control; 2, with 10 ng/mL FGF2; 3, with 10 ng/mL FGF2 and 1 μM PD173074 (FGFRi), with media refreshed every three days. Results shown are representative of three independent experiments. B, MEK inhibitor trametinib abrogated the resistance to palbociclib alone (Top panel) or the combination of fulvestrant and palbociclib (Bottom panel) conferred by FGFR1 or FGFR2. Cells were treated with different conditions as indicated for six days before CellTiter-Glo assay. Concentration of drugs used: 10 ng/mL FGF2; 100 nM fulvestrant (Fulv); 1 μM palbociclib (Palbo); 500 nM trametinib. * p-value < 0.05, ** p-value < 0.01, *** p-value < 0.001. n.s. not significant. Student t-test was performed for pair-wise comparisons. Results shown are ± SD and representative of three independent experiments. C, Trametinib blocked ERK phosphorylation and reduced CCND1 and p-Rb levels. Cells were treated as indicated daily for two days before protein harvest and western blot. Results shown are representative of two independent experiments.

In the presence of fulvestrant and palbociclib, FGFR1 or FGFR2 overexpression was accompanied by increased p-Rb and CCND1 levels, both of which were partially reversed by trametinib (Fig.5C). CCND1 knockdown in cells overexpressing GFP, FGFR1 or FGFR2 impaired cell proliferation similarly across all three cell lines without affecting the IC50 of fulvestrant (Supplemental Data Fig.S17). This suggests that the proliferation advantage provided by active FGFR signaling is partially dependent on CCND1. This is consistent with prior results suggesting that CCND1 was involved in FGF2-mediated drug resistance [29].

Clinical evidence also supports the finding that FGFR alterations can cause resistance to CDK4/6 inhibitors. Following the acquisition of *FGFR2* N550K (along with *FGFR2* amplification), Pt 0300350 did not respond to the combination of letrozole and palbociclib (Fig.3), suggesting that *FGFR2* alterations may lead to intrinsic resistance to the combination of endocrine therapy and CDK4/6 inhibitors. Another patient with an *FGFR2* N550K mutation (FM Patient 2) also did not respond to the combination of fulvestrant and palbociclib (Supplemental Data Fig.S8). Collectively, this suggests targeting the FGFR pathway may also be a viable strategy to overcome FGFR/FGF-mediated resistance to SERDs and CDK4/6 inhibitors.

### FGFR2 mutations found in patients are activating with differential sensitivity to FGFR inhibitors

We identified 3 acquired mutations in the kinase domain of FGFR2 in patients who developed resistance to endocrine therapy. Two of these, FGFR2 N550K and K660N, are known activating FGFR2 mutations that have been previously identified in breast cancer [25, 27]. FGFR2 N550K is part of the molecular brake at the kinase hinge region, which allows the receptor to adopt an active conformation more easily (Fig.6A). N550K is a recurring hotspot mutation reported to confer resistance to several FGFR inhibitors including PD173074 and dovitinib [30]. FGFR2 K660N is located in a conserved region in the tyrosine kinase domain and has been confirmed to increase kinase activity [27, 30]. The third mutation, FGFR2 M538I, has not been previously reported in breast cancer. Based on its location, M538I appears to stabilize the active kinase confirmation by strengthening the hydrophobic spine of the FGFR2 kinase [30] (Fig.6A).

**Figure 6.**
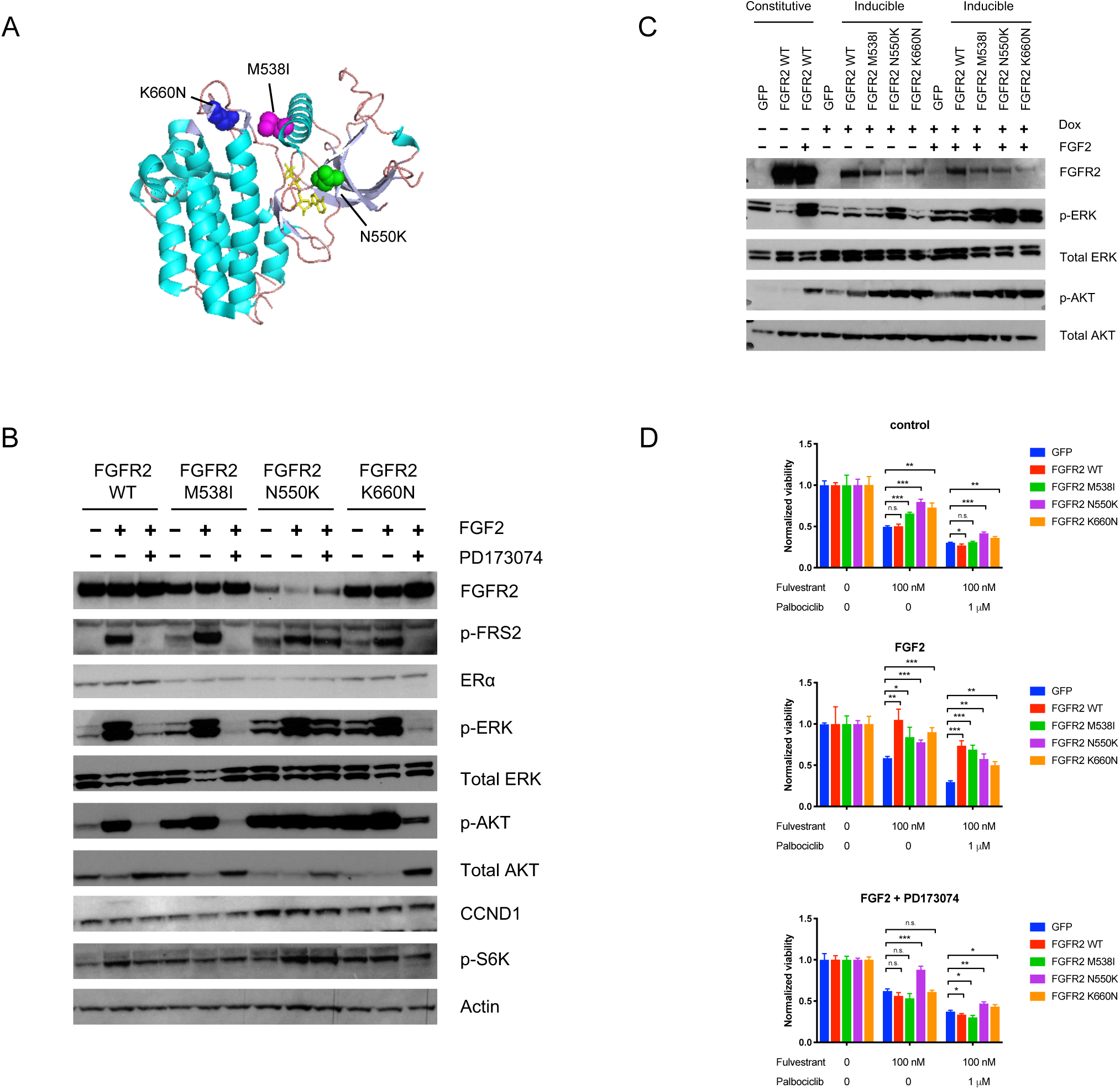
FGFR2 M538I, N5550K and K660N were activating mutations and conferred resistance to fulvestrant and/or palbociclib. A, Crystal structure of activated FGFR2 protein with mutations shown. FGFR2 is in complex with ATP analog (in yellow) and substrate peptide (PDB ID: 2PVF). B, Stable cell lines overexpressing FGFR2 wildtype (WT), M538I, N550K, and K660N were treated with 10 ng/mL FGF2 and/or 1 μM PD173014 for one hour before protein harvest. Results shown are representative of two independent experiments. C, Tetracycline-inducible cell lines that express FGFR2 WT, M538I, N550K and K660N were established and treated with 100 ng/mL doxycycline (Dox) to induce gene expression. Before protein harvest, cells were treated with doxycycline 24 hours before FGF2 stimulation for 3 hours followed by another 3 hours of FGF2 stimulation. Cells were treated with heparin (1 μg/mL) that facilitates the binding between FGF2 and FGFR2. Protein lysates from T47D cells expressing GFP and T47D cells with constitutive overexpression of FGFR2 were used as control. Results shown are representative of three independent experiments. D, Stable cell lines constitutively overexpressing GFP or FGFR2 constructs (as previously described) were examined for sensitivity to fulvestrant or combination of fulvestrant and palbociclib with or without the treatment of FGF2 and/or PD173074. * p-value < 0. 05, ** p-value < 0.01, *** p-value < 0.001, calculated as compared to GFP group in all conditions. Student t-test was performed for pair-wise comparisons. Results shown are ± SD and representative of three independent experiments.

We expressed all three FGFR2 kinase domain mutants in T47D cells through lentiviral transduction, as well as wildtype (WT) FGFR2 and GFP as negative controls. All three mutants elicited higher kinase activity than WT FGFR2 constitutively, demonstrated by levels of p-FRS2, a direct substrate for FGFR2 (Fig.6B). MAPK and AKT signaling were also increased, as indicated by increased p-ERK and p-AKT levels, respectively (Fig.6B). The addition of FGF2 ligand further enhanced downstream signaling for all FGFR2 mutants, and the enhanced signaling was blocked by PD173074 for FGFR2 M538I and K660N, but not for N550K (Fig.6B).

FGFR2 mutants were also expressed under a tetracycline responsive promoter in T47D cells grown in low doses of doxycycline to determine the functionality at lower expression levels. At lower levels of expression, FGFR2 N550K still led to increased levels of p-ERK and p-AKT, independent of FGF2 ligand stimulation. FGFR2 M538I and K660N also resulted in higher p-ERK and p-AKT levels in the presence of FGF2 as compared to FGFR2 WT (Fig.6C). Taken together, these results indicate that all three FGFR2 mutations acquired in breast cancer patients are functionally active – FGFR2 N550K is constitutively active while FGFR2 M538I and K660N may be more ligand-dependent at low levels of expression.

All 3 FGFR2 mutants led to modest resistance to fulvestrant (Fig.6D), which was enhanced in the presence of FGF2 ligand. PD173074 resensitized cells overexpressing FGFR2 M538I and FGFR2 K660N as well as WT FGFR2 to fulvestrant, but not cells overexpressing FGFR2 N550K (Fig.6D). Consistent results were observed when cells were treated with the combination of fulvestrant and palbociclib (Fig.6D). Similar results were obtained in MCF7 cells (Supplemental Data Fig.S18).

### Transcriptional changes induced by FGFR/FGF include ER reprograming and MAPK activation

To examine transcriptional changes associated with FGFR pathway activation, we performed RNA-Seq on ER+ T47D cells overexpressing FGFR1, FGFR2 (wildtype), FGFR2 activating mutants (M538I, N550K, K660N), or FGF3, as well as on GFP and parental lines as controls (Supplemental Table.9). We combined these profiles with transcriptional profiles we previously generated [11] from of ER+ T47D cells overexpressing wildtype and kinase-dead HER2, HER2 activating mutants (S653C, L755S, V777L and L869R), and ESR1 Y537S, for a total of 15 genetic perturbations examined.

Linear discriminant analysis of these profiles indicated that samples from cells overexpressing FGFR1, FGFR2, FGFR2 activating mutants, FGF3, wildtype HER2, or HER2 activating mutants are separated from controls (parental cells, GFP expressing, or kinase-dead HER2 D845A) along the first LD component (LD1), as well as from a separate group of cells overexpressing mutant ESR1 along the second LD component (LD2), indicating a common RTK/growth factor-driven cell state that is distinct from the mutant ER state (Fig. 7A). Along LD1, the cells overexpressing FGF3 have distinct scores from those overexpressing FGFR (p.value = 2.12×10^-10^, Welch’s t-test), suggesting that in our model system, FGF ligand overexpression is different from FGFR overexpression (Fig. 7A). The overall signature strength of RTKs and growth-factor signaling genes (defined in Supplemental Methods and shown in Supplemental Table.10) were highly correlated with scores on LD1 (Spearman’s rho= 0.926), suggesting that LD1 broadly represents RTK and growth-factor pathway activation.

**Figure 7.**
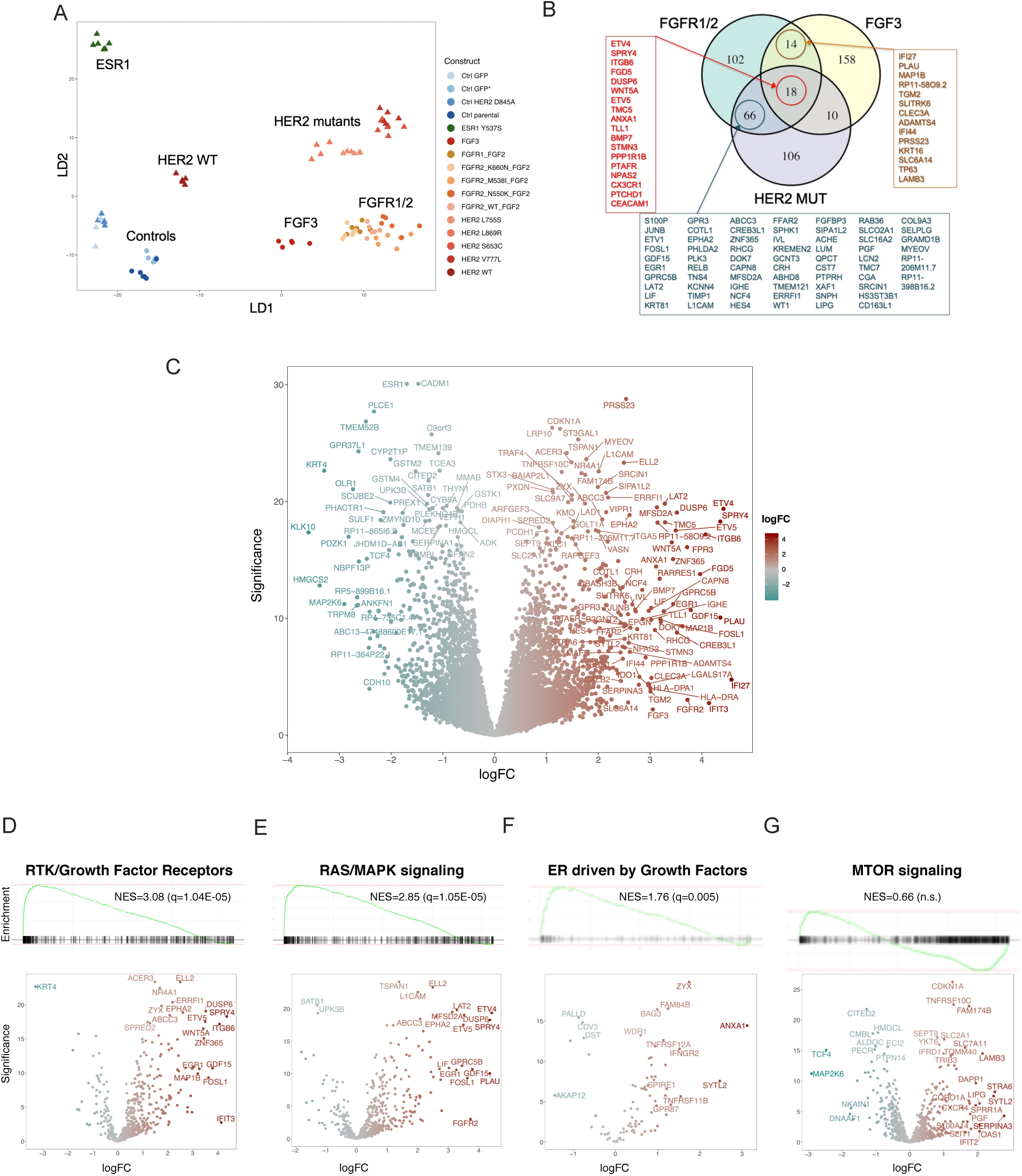
Transcriptional cell-state analysis of FGF/R activating perturbations reveals a distinct cell-state associated with MAPK activation and ER-reprograming. A, Linear Discriminant Analysis (LDA) projection of FGFR/FGF-activated cells and controls. Two-dimensional visualization of in the transcriptional footprints driven by FGF3, FGFR1, FGFR2, FGFR2 activating mutants, and GFP, as well as HER2 activating mutants, wild-type HER2, kinase-dead HER2 D845A, and ESR1 Y537S (previously published, with data points from HER2 study indicated by triangles [11]), all treated with DMSO. B, Venn diagram showing commonalities and differences among upregulated genes for cells expressing FGFR (FGFR1, FGFR2, and FGFR2 activating mutants), FGF3, and HER2 activating mutants. Upregulated genes are based on log fold change (logFC) comparing the activating constructs and controls under DMSO. C, Gene Set Enrichment Analysis (GSEA) plots and Volcano plot representation of differentially expressed genes (DEGs) comparing the transcriptional footprints from cells expressing FGFR1, FGFR2, FGFR2 activating mutants, and FGF3, collectively vs. controls, all under DMSO. The FGF/R-ACT transcriptional state was derived from significant DEGs. D-G, Selective volcano plots and GSEA plots indicate several pathways which are activated in the FGF/R-ACT state: D, RTK/Growth Factor Receptors signaling genes (624 genes), E, RAS/MAPK genes (701 genes). F, ER signaling driven by Growth Factors genes (95 genes), G, MTOR pathway genes (953 genes). See Supplemental Table.11 for gene sets, Supplemental Table.12 for a comprehensive list of gene sets and pathway associations. NES, normalized enrichment score; n.s. not significant.

Using differential expression analysis to compare activating constructs with controls, we defined transcriptional signatures FGFR1/2, FGF3, and HER2-MUT. The FGFR1/2 signature was generated by comparing FGFR1, FGFR2, FGFR2 M538I, FGFR2 N550K, and FGFR2 K660N overexpression (all with FGF2 ligand) to parental and GFP (with and without FGF2 ligand). The FGF3 signature was generated by comparing FGF3 overexpression (without FGF2 ligand) to parental and GFP (with and without FGF2 ligand). The HER2-MUT signature was generated by comparing the HER2 activating mutants (HER2 S653C, L755S, V777L and L869R) with GFP controls.

Comparison of the top 200 highly expressed genes in each of the FGFR1/2, FGF3, and HER2-MUT signatures (Supplemental Table.11) highlights the high degree of similarity between these groups, with 32 genes common to the FGFR1/2 and FGF3 signatures (Odds-ratio=39.63 p.value=4.7×10^-37^, two-sided Fisher’s exact test), 84 genes common to the FGFR1/2 and HER2-MUT signatures (Odds-ratio=219.4, p.value=1.025×10^-140^, two-sided Fisher’s exact test), and 18 upregulated genes common to all 3 groups (Fig 7B).

We next defined an FGF/R-ACT transcriptional state based on 377 genes that are upregulated in FGF3 and FGFR1/2 cell lines compared with controls (Fig. 7C and Supplemental Table.10). Gene Set Enrichment Analysis [31] of the FGF/R-ACT state vs. 5,150 previously characterized gene sets (Supplemental Table.12) demonstrated that the shared FGF/R-ACT state is enriched for upregulation of RTK/growth factor receptors signaling, RAS/MAPK signaling, and growth factor–induced ER target genes (Fig. 7D-F), similar to what we have previously shown for HER2 activating mutants [11]. This is consistent with a model in which FGFR/FGF activation leads to ER reprogramming via MAPK pathway activation, potentially shifting the transcriptional spectrum from genes activated by transcription factor AF2 to those activated by AF1. Of note, the genes associated with mTOR signaling were variably upregulated and downregulated in FGF/R-ACT state (Fig. 7G), suggesting only moderate mTOR activation as compared to the robust MAPK activation signal observed. Similar transcriptional signatures were seen when the FGF/R-ACT signature was characterized under fulvestrant treatment, demonstrating that the FGF/R-ACT state is present even when ER signaling is inhibited, consistent with the phenotype of fulvestrant resistance (Supplemental Table.11-12).

### Activating FGFR2 mutations can be targeted with irreversible FGFR inhibitors

FGFR2 M538I and N550K have been shown to confer resistance to multi-kinase inhibitor dovitinib in BaF3 cells, and N550K is also resistant to PD173074 (Fig.6 B and D)[30]. Because of the differential responses of these FGFR2 mutants to PD173074, we tested the ability of additional FGFR inhibitors to resensitize cells expressing these mutants to fulvestrant. FIIN-2 and FIIN-3 are two irreversible covalent pan-FGFR inhibitors that target a cysteine conserved in FGFR1-4 and have exquisite selectivity for some FGFR2 mutations including M538I and K659N [32]. In addition, AZD4547 is a selective FGFR inhibitor that was previously shown to inhibit FGFR2 N550K [33].

Both FIIN-2 and FIIN-3 were more effective in inhibiting the downstream signaling (p-FRS2, p-ERK and p-AKT) induced by FGFR2 N550K as compared to PD173074 and AZD4547 (Fig.8A). Furthermore, FIIN compounds reduced the level of downstream effectors back to baseline levels, with FIIN-3 being more potent than FIIN-2 (Supplemental Data Fig.S19). T47D cells stably overexpressing FGFR2 mutant were exquisitely sensitive to FIIN-2 and FIIN-3 as compared to cells expressing GFP or FGFR2 WT (Supplemental Fig.S20). The resistance to fulvestrant induced by both WT FGFR2 and all mutant FGFR2 was completely blocked by FIIN-2 (1 μM) and FIIN-3 (100 nM), but not by PD173074 (1 μM) and AZD4547 (1 μM) (Fig.8B). While resistance to WT FGFR1/2 and FGFR2 M538I and FGFR2 K660N can be reversed by multiple FGFR inhibitors, for some mutants like FGFR2 N550K, only the irreversible pan-FGFR inhibitors successfully resensitized cells to fulvestrant, highlighting the fact that specific resistance mutations might require different strategies to overcome or preempt endocrine resistance.

**Figure 8.**
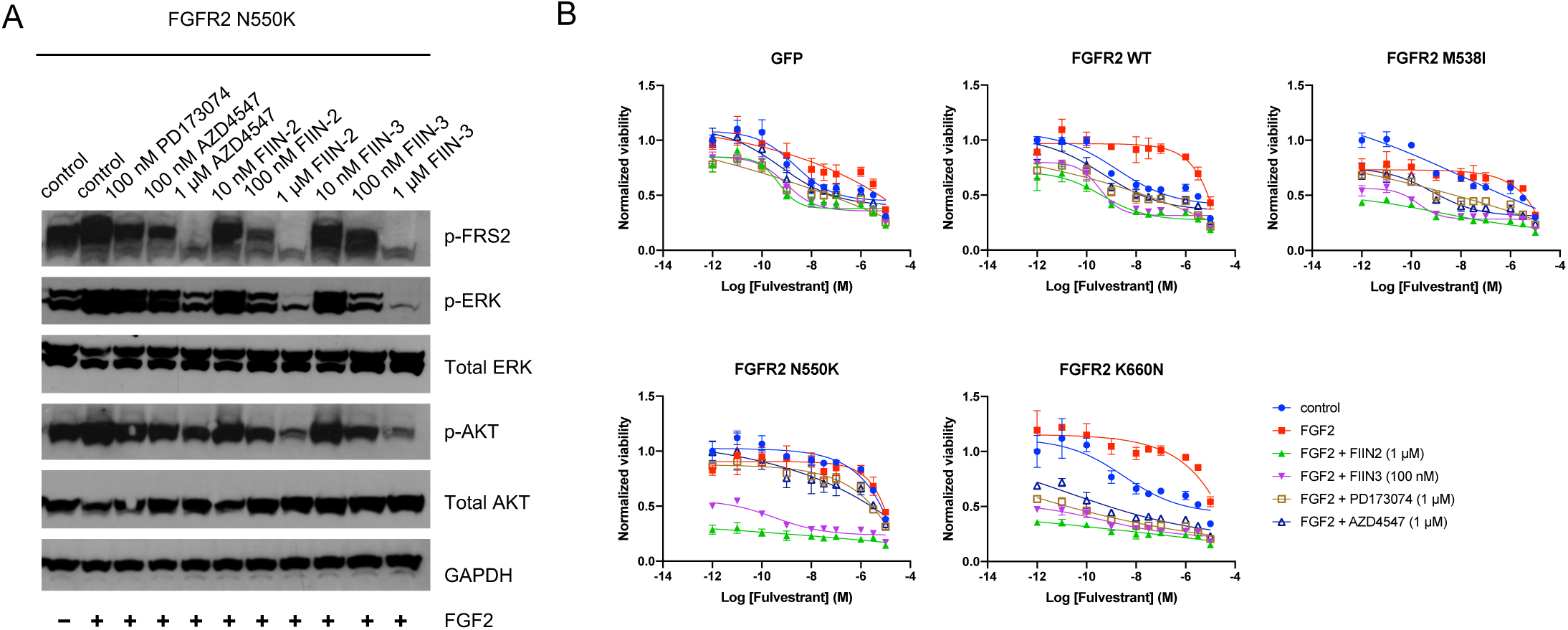
Activating FGFR2 mutations were targeted by irreversible kinase inhibitors FIIN-2 and FIIN-3. A, T47D cells overexpressing FGFR2 N550K cells were treated as indicated for three days and retreated for three hours before protein harvest and western blot. Results shown are representative of two independent experiments. B, All stable cells lines expressing GFP or FGFR2 constructs were treated with fulvestrant under the following conditions: control, 10 ng/mL FGF2, 10 ng/mL FGF2 with 1 μM AZD4547, 10 ng/mL FGF2 with 1 μM FIIN-2, or 10 ng/mL FGF2 with 100 nM FIIN-3. Drug response curves were determined by CellTiter-Glo. Results shown are ± SD and representative of three independent experiments.

Next, we evaluated the effect of ten different drug combinations on the transcriptional signatures in FGFR/FGF expressing cell lines and control cells profiled by RNA-seq. The activated RAS/MAPK signature, ER signaling driven by growth factors signature, and FGF/R-ACT signature present in cells overexpressing FGFR/FGF constructs persisted under treatment with fulvestrant, palbociclib, or their combination (Fig. 9). Treatment with FIIN-3, alone or in combination with fulvestrant and/or palbociclib largely reversed the activation of all three signatures (Supplemental Table.13). For example, all FGFR/FGF constructs showed suppression of FGF/R-ACT signature by FIIN-3 treatment (FIIN-3 vs. DMSO, median effect size = −5.42, Cohen’s D test with Hedges correction; median p.value = 7.858×10^-6^, Welch’s t-test), albeit with less suppression observed for FGF3 (effect size = −1.86, p.value = 0.00028) and FGFR2 N550K (effect size = −3.74, p.value = 6.722×10^-5^) constructs (Fig. 9; Supplemental Table.13). Effective suppression of all three signatures was also achieved by trametinib plus fulvestrant, with or without palbociclib (Fig. 9; Supplemental Table 13),

**Figure 9.**
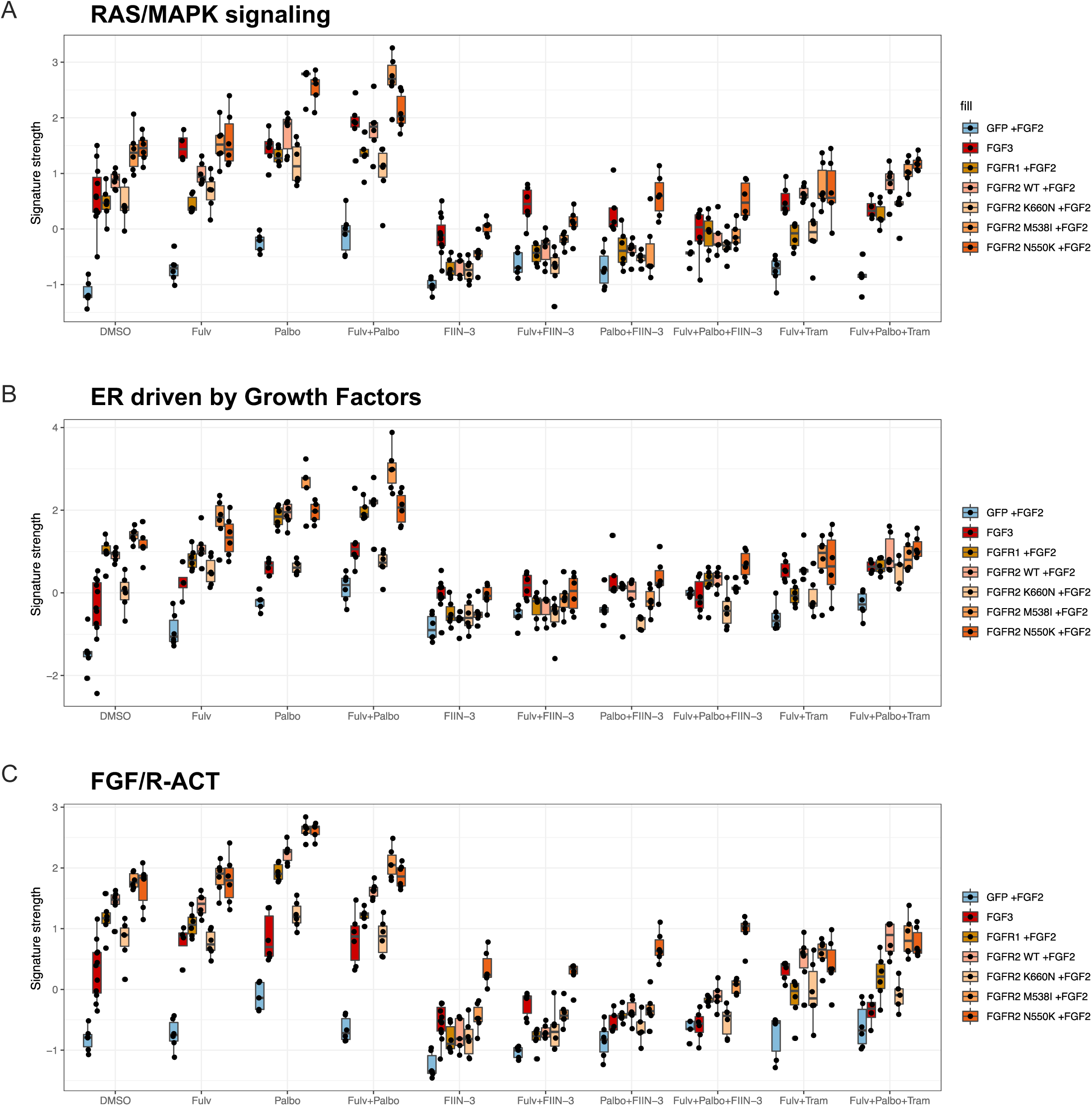
RNA-seq measurement of oncogenic signatures reflects differential sensitivity of FGF/R activating perturbations under various drugs. Three oncogenic signatures were compared across various drug conditions for resistant FGFR/FGF cell lines. A, RAS/MAPK signature. B, ER signaling driven by growth factors signature. C, FGF/R-ACT signature. The signature strength for each sample is depicted in the Y-axis with a Z-score scaled across all 647 samples. Each box plot represents the distribution of the signature strength among replicates of each experimental condition as indicated. Drug conditions include: “DMSO” (No Drug), “Fulv” (Fulvestrant), “Palbo” (Palbociclib), “Fulv+Palbo” (Fulvestrant and Palbociclib), “FIIN-3”, “Fulv+FIIN-3” (Fulvestrant and FIIN-3), “Palbo+FIIN-3” (Palbociclib and FIIN-3), “Fulv+Palbo+FIIN-3” (Fulvestrant, Palbociclib and FIIN-3), “Fulv+Tram” (Fulvestrant and Trametinib), “Fulv+Palbo+Tram” (Fulvestrant, Palbociclib and Trametinib. Construct perturbations include overexpression of GFP (Control), FGF3, FGFR1, and FGFR2 (WT, K660N, M538I and N550K). For cell lines overexpressing FGFR1/2, FGF2 was supplemented in the media. See Supplementary methods for details and Supplemental Table.11 for gene sets definitions.

### Targeting MAPK pathway via SHP2 inhibition can overcome FGFR-induced endocrine resistance

Data from the gain-of-function ORF screen, the transcriptomic analysis, and our individual *in vitro* experiments together suggest that MAPK pathway may represents a common node for drug resistance in ER+ metastatic breast cancer. FGFR2 mutants, in particular, rendered cells more sensitive to trametinib than did GFP or WT FGFR2 (Fig.10A), further supporting the finding that FGFR signaling requires the MAPK pathway in this context. Src homology phosphotyrosyl phosphatase 2 (SHP2) is protein downstream of RTKs that is required for RAS activation. Several recent studies have demonstrated that co-targeting SHP2 prevented adaptive resistance to MEK inhibition in multiple RAS-driven cancer models[34–36]. Consistent with this, cell lines expressing FGFR2 mutants were hypersensitive to the SHP2 inhibitor SHP099 (Fig.10B). We next examined the ability of combinations of SHP099, trametinib, fulvestrant, and/or palbociclib to overcome resistance in FGFR-expressing cell lines. Similar to trametinib, SHP099 as a single agent partially rescued resistance to fulvestrant and/or palbociclib conferred by FGFR1, and completely rescued resistance conferred by FGFR2. Cells expressing the FGFR2 constructs (wildtype or activating mutations) were particularly sensitive to the treatment regimen of trametinib and SHP099 in addition to fulvestrant and palbociclib (Fig.10C-D), in comparison with minimal inhibitory effect of MEK and SHP2 inhibition in GFP control cells. The cell viability phenotype was accompanied with consistent changes in p-ERK levels; SHP099 alone partially reduced ERK phosphorylation and the combination of SHP099 and trametinib completely abrogated p-ERK levels (Supplemental Fig.S21). These results suggest that targeting MEK and SHP2 may serve as an even more effective strategy to overcome multiple forms of FGFR pathway mediated resistance to endocrine therapy and CDK4/6 inhibitors in MBC, as well as, potentially, other RTK-induced mechanisms of resistance.

**Figure 10.**
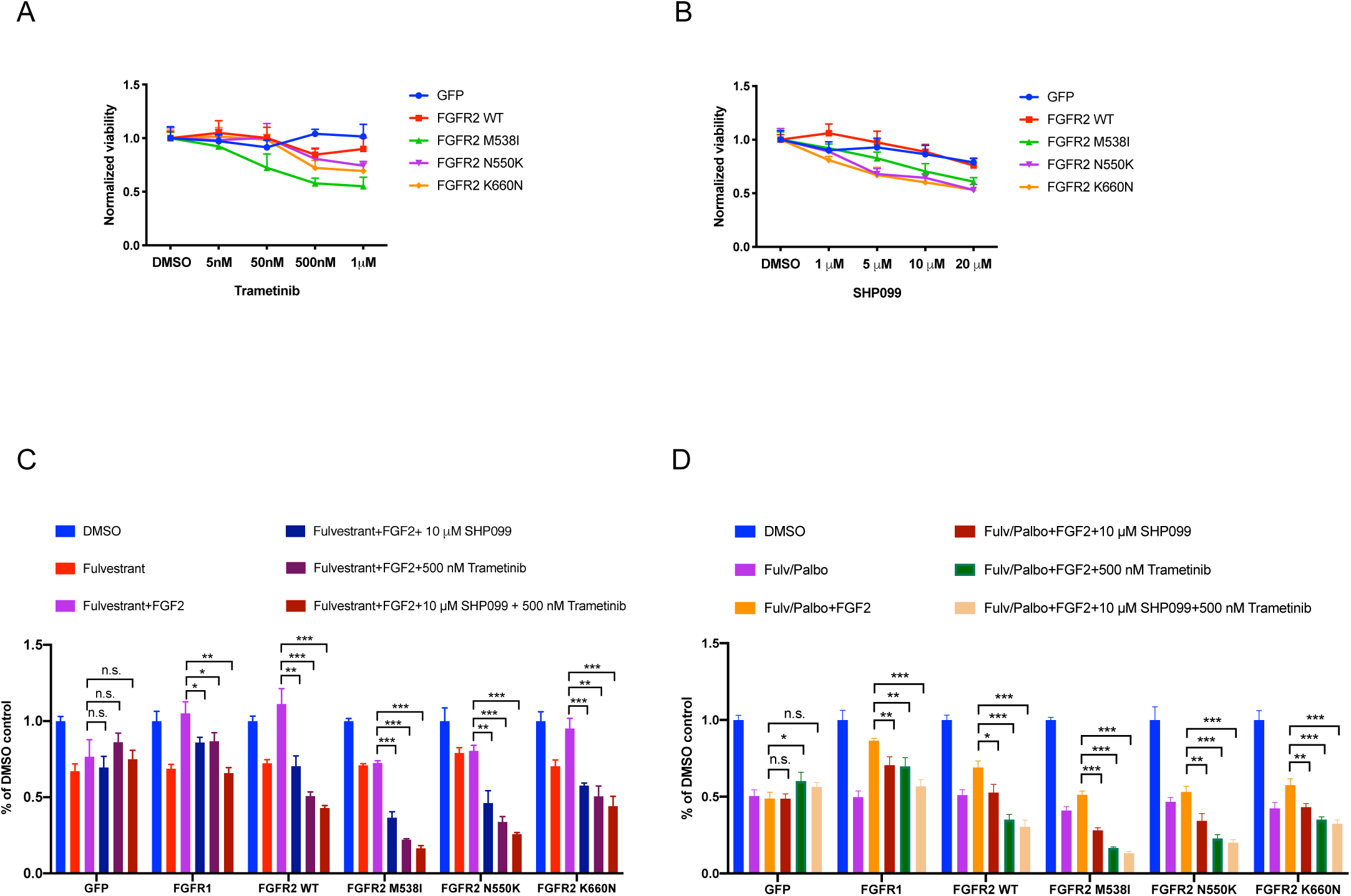
FGFR-induced resistance phenotype was reversed by MEK and SHP2 inhibition. A and B, Trametinib (A) and SHP099 (B) sensitivity of T47D cell lines expressing GFP or different FGFR2 constructs was examined respectively. Results shown are ± SD and representative of two independent experiments. C and D, SHP099 as a single agent and in combination with trametinib rescued resistance to fulvestrant (C) and/or palbociclib (D) conferred by FGFR1 or FGFR2 wildtype and mutant constructs. Concentration of drugs used: 10 ng/mL FGF2; 100 nM fulvestrant; 1 μM palbociclib; 500 nM trametinib; 10 μM SHP099. * p-value < 0.05, ** p-value < 0.01, *** p-value < 0.001. n.s. not significant. Student t-test was performed for pair-wise comparisons. Results shown are ± SD and representative of three independent experiments.

## Discussion

In this study, we used a genome-scale gain-of-function screen to identify potential mechanisms of resistance to selective estrogen receptor degraders. We nominated several different candidate resistance genes and pathways, particularly genes in the ERBB pathway, FGFR pathway, insulin receptor signaling and the MAPK pathway. Consistent with this finding, genomic profiling of paired pre-treatment and post-treatment tumor samples from 60 patients with ER+ metastatic breast cancer who developed resistance to endocrine therapy identified acquired alterations in *FGFR1, FGFR2*, and *FGF3* in 20% of patients. Experimental studies confirmed that these alterations confer resistance to endocrine therapy as well as CDK4/6 inhibitors, through activation of MAPK pathway, and demonstrated that this resistance can be reversed by FGFR inhibitors, MEK and SHP2 inhibitors. Taken together, our results suggest that activating FGFR pathway alterations are a distinct mechanism of acquired resistance to multiple forms of ER-directed therapy in MBC that can be overcome by FGFR and/or MAPK pathway inhibitors.

Our findings are consistent with two recently published results. Formisano, Arteaga, and colleagues identified *FGFR1* amplification as a mechanism of resistance to CDK4/6 inhibitors [37]. In another study, Drago et al. demonstrated that FGFR1 amplification confers resistance to ER, PI3K and CDK4/6 inhibitors while retaining TORC sensitivity [38]. Our results also extend our understanding of endocrine resistance with multiple novel findings. Experimentally, this includes functional screen data for SERD resistance that includes over 10,000 genes that implicate multiple novel pathways and genes in endocrine resistance. It also includes evidence showing that FGFR1 overexpression, FGFR2 overexpression, FGF3 overexpression, and the specific three FGFR2 mutants found in patients *all* result in resistance to endocrine therapy and/or CDK4/6 inhibitors that can be overcome by specific targeted therapies. Clinically, the WES analysis of 60 pairs of pre-treatment and post-resistance tumor biopsies identified multiple alterations in the FGFR pathway in post endocrine therapy tumor biopsies, including FGF3 amplifications as well as FGFR1 and FGFR2 alterations. Most significantly, we present here a rigorous demonstration that 50% of cases these FGFR/FGF pathway alterations are *acquired* after resistance to therapy.

Our genome-scale screen provided a comprehensive view into the resistance mechanisms to SERDs. Similar resistance genes were nominated for fulvestrant and GDC-0810, thereby confirming the two drugs have similar mechanism of action. Of note, two ESR1 ORFs conferred resistance specifically to GDC-0810 but not fulvestrant, possibly due to GDC-0810 having a less potent effect on ER degradation than fulvestrant [21, 39]. Among the resistance mechanisms shared by fulvestrant and GDC-0810, many are frequently altered in ER+ MBC, such as *CCNDs/CDK6, KRAS/MAPK, EGFR/ERBB2* and *PIK3CA/AKTs/PIMs*, and agents targeting those alterations are under clinical development to be combined with endocrine therapy [14, 40]. We also identified potential resistance mechanisms that are not characterized to the same extent, such as G protein-coupled receptors, Wnt pathway (*FZD10, RSPO1, RSPO3*) and Src family kinases (*YES1, FYN, FGR*), providing clues as to the potential crosstalk between these pathways and ER signaling [41–43] and suggesting that breast cancer patients harboring functional alterations in these pathways may develop resistance to SERDs. Notably, some of the resistance genes were also nominated in gain-of-function screens designed to identify resistance mechanisms for MAPK pathway inhibitors in melanoma (*AXL, CRK, CRKL, FGR, GPCR* genes) [15] and PI3K inhibitors in ER+ breast cancer (*AKT1, AKT2, CRKL, FGF3, FGF10, PIM* genes) [17]. This may reveal multi-drug resistance mechanisms and thus guide clinical drug combinations to overcome resistance. We have provided the full genome-scale screen data as a resource to the community of researchers interested in resistance to ER-directed therapies as well as the biology of estrogen receptor dependencies in ER+ breast cancer.

Our ultimate goal is to identify resistance mechanisms that are clinically relevant and can be therapeutically targeted. By comparing paired pre-treatment and post-treatment tumors, our evolutionary analyses identified acquired *FGFR1* and *FGFR2*, and *FGF3* alterations in 12 out of 60 post-treatment samples, further highlighting a potential role for the FGFR pathway in driving drug resistance and disease progression.

Most notably, all four alterations in FGFR2 in our cohort were found to be acquired after the development of resistance to endocrine therapy. Our overall findings are consistent with other recent studies which noted some patients with acquired *FGFR1* and *FGFR2* alterations following treatment of endocrine therapy [37, 38, 44, 45], and provides a mechanistic explanation for these acquisitions.

This analysis was enabled by a novel method we developed to compare the magnitude of amplification in matched pre- and post-treatment samples while considering key confounders to allow for more reliable assignment of copy gain or loss. Since matched tumor samples of the same patient are highly variable in the cancer cell fraction (purity) and often variable in ploidy (with genome duplication taking place in the metastatic setting), we computed the purity-corrected copy number above ploidy and set a relatively stringent threshold of changes in CNAP to define acquired amplification (see Methods), as cancer clones bearing amplifications with high focality and magnitude in FGFR/FGF genes are more likely to induce dependency on FGFR pathway and result in endocrine resistance.

Our genomic analysis has some limitations and caveats. The observed alterations may not exclusively result from endocrine therapy as some patients received other therapies between the two collected biopsies. Moreover, tumors with FGFR/FGF alterations also harbor alterations in other cancer genes, which may contribute to drug resistance as well (Supplemental Data Fig.S6 and Supplemental Table.5). Despite these caveats, with the evidence from unbiased screens, genomic evidence in relevant patient samples, and confirmatory experimental models, the FGFR pathway clearly emerges as a clinically important resistance mechanism for SERDs and CDK4/6 inhibitors.

Strategies to target the FGFR pathway in breast cancer patients with FGFR alterations are currently being assessed in clinical trials. FGFR inhibitors currently under clinical development in breast cancer include non-selective tyrosine kinase inhibitors (dovitinib and lucitanib), FGFR1-3 selective inhibitors (AZD4547 and BGJ398), and others, although clinical trials have achieved mixed results to date [46–51]. The combination of FGFR inhibitors and endocrine therapy is also being clinically investigated. For example, the combination of dovitinib and fulvestrant showed promising clinical activity [46]. As FGFR pathway activation also results in resistance to CDK4/6 inhibitors, a triple combination with the addition of CDK4/6 inhibitors may also be considered. One challenge for the use of FGFR inhibitors is to identify reliable biomarkers. Our results suggest focal and high level amplifications, clonal activating mutations or high expression levels of FGFR and FGF genes, particularly in the metastatic setting, may be used to guide the clinical use of FGFR inhibitors. Activating alterations in *FGFR2*, which are rare in primary treatment naïve breast cancer but appear to be clonally acquired in a subset of patients with resistant ER+ MBC, may be a particularly good biomarker for the development of FGFR inhibitors.

Our work also highlights that the effective clinical use of FGFR inhibitors needs to consider the variable drug sensitivity of different FGFR2 mutations, which were acquired in some patients following endocrine therapy. The two irreversible pan-FGFR kinase inhibitors, FIIN-2 and FIIN-3, had superior efficacy in targeting all FGFR2 mutants including N550K when compared to other FGFR inhibitors, although *in vivo* efficacy, off-target effects and toxicity of FIIN compounds still warrant further investigation.

Alterations in FGFR1 and FGFR2 activated the MAPK pathway, and MEK inhibition was able to overcome the resistance conferred by FGFR pathway to some degree. We previously demonstrated acquired ERBB2 mutations resulted in a reprogrammed ER signature and an elevated MAPK transcriptional signature [11]. In this study, the transcriptional analysis results suggest that MAPK activation resulting from FGFR/FGF overexpression is more pronounced than mTOR activation, which had a mixed signature. This is consistent with our experimental findings, which demonstrated that MAPK inhibition is particularly effective in cells expressing FGFR2 activating alterations. Other MAPK genes were also found to be enriched in the gain-of-function screen. Furthermore, increased frequency of alterations in MAPK pathway genes have been found in tumors post hormonal therapy, including *EGFR, ERBB2* and *NF1* [14]. The fact that multiple mechanisms of resistance to ER-directed therapies and/or CDK4/6 inhibitors activate the MAPK pathway suggests that this may be an important node of resistance in ER+ MBC. Thus, combining endocrine therapy and CDK4/6 inhibitors with agents that target MAPK pathway, such as MEK inhibitors and/or SHP2 inhibitors [52, 53], may be a unifying strategy to overcome or prevent resistance resulting from multiple genetic aberrations that lead to resistance in ER+ MBC.

In summary, the integration of a functional genomic screen and genomic analysis of pre- and post-treatment biopsies revealed the FGFR pathway as an important resistance mechanism for endocrine therapy and CDK4/6 inhibitors in ER+ breast cancer. With the increasing use of SERDs and CDK4/6 inhibitors in the clinic, we anticipate that the prevalence of FGFR/FGF alterations might increase in the future. Targeting the FGFR pathway with FGFR inhibitors or agents that target downstream MAPK signaling may improve clinical outcomes in patients with aberrations in FGFR/FGF genes. Furthermore, our study highlights the need to sequence metastatic biopsy or blood biopsies at the time of resistance to identify patients with these alterations who may benefit from targeting the FGFR pathway.

## Methods

### Cell culture

293T, T47D and MCF7 cells were purchased from American Type Culture Collection (ATCC) and were cultured as described in the Supplemental Methods.

### Genome-Scale Gain-of-Function Screen

The pooled lentiviral ORF library hORFeome [22] consists of 17,255 barcoded human open reading frames (ORFs), corresponding to 10,135 distinct human genes with at least 99% nucleotide and protein match. These ORFs were cloned into pLX317 vector and pooled together for transfection into 293T cells to make pooled lentivirus (with 2nd generation packaging plasmids). In 6-well plates, pooled lentivirus was infected in cells to achieve ∼50% infection rate and ensure ∼1000 infected cells per ORF for 17255. Media was supplemented with 4 μg/mL polybrene (Thermo Fisher Scientific # TR1003G) to boost transfection efficiency. After infection, cells were pooled and selected with 1.5 μg/mL puromycin for 5 days. Upon completion of selection, cells were plated for three different drug conditions: DMSO, 100 nM fulvestrant, 1 µM GDC-0810. There were three replicates for each condition screened. A subset of cells was saved for sequencing as early time point (ETP) samples to confirm ORF representation. The dose for each drug was chosen for the two drugs to achieve potent anti-proliferation effect that could be rescued with ESR1 mutant Y537N and Y537S. Infected cells were passaged upon confluency and maintained in DMSO or drugs for 21 days to allow sufficient time for cells carrying resistance to be enriched from the population. At the end of the time course, cells were harvested for isolating genomic DNA as late time point samples (LTP).

All genomic DNA samples were amplified with PCR primers flanking the ORF region and sequenced. The ORF representation at the final harvesting (LTP) is compared to the representation of ORFs in cells collected before drug addition (ETP). Cells carrying ORFs that are driving resistance will grow and gradually enrich the population and therefore, will be over-represented in the sequencing data for the final passage compared to the early time point. An ORF with significant enrichment (a Z score >3) is defined as a resistance candidate gene. A secondary validation screen was performed as described in the Supplemental Methods.

### Patients and Tumor Samples

Prior to any study procedures, all patients provided written informed consent for research biopsies and whole exome sequencing of tumor and normal DNA, as approved by the Dana-Farber/Harvard Cancer Center Institutional Review Board (DF/HCC Protocol 05-246). Metastatic core biopsies were obtained from patients and samples were immediately snap frozen in OCT and stored in −80°C. Archival FFPE blocks of primary tumor samples were also obtained. A blood sample was obtained during the course of treatment, and whole blood was stored at −80°C until DNA extraction from peripheral blood mononuclear cells (for germline DNA) was performed. In a few instances, cell free DNA was obtained from plasma for circulating tumor DNA analysis, as previously described[54].

### Whole Exome (WES) analysis

DNA was extracted from primary tumors, metastatic tumors, plasma, and peripheral blood mononuclear cells (for germline DNA) from all patients and whole exome sequencing was performed, as detailed in the Supplemental Methods. Sequencing data were analyzed using tools to identify somatic point mutations and small insertions/deletions (indels), and copy number changes using established algorithms (see Supplemental Methods).

To better measure segment-specific copy-number, we subtracted the genome ploidy for each sample to compute copy number above ploidy (CNAP). CNAP of at least 3 are considered as amplifications (AMP), CNAP below 3 are considered low amplification and ignored in our analysis). CNAP of at least 6 are considered high amplifications (HighAMP), and CNAP of at least 9 and fewer than 100 genes [55] is considered very high focal amplification (FocalAMP).

The evolutionary classification of amplifications accounts for the magnitude of the observed copy-number difference between the pre-treatment and the post-treatment samples. If the difference between the CNAP of the post-treatment and the CNAP of the pre-treatment is smaller than 50%, the amplification is defined as “Shared”. If the CNAP of the post-treatment is larger than the CNAP by more than 50% and the lower pre-treatment CNAP is not at “FocalAMP” level, the evolutionary classification is “Acquired”. If CNAP of the post-treatment is smaller by at least 50%, comparing to the pre-treatment sample and the lower post-treatment CNAP is not at “FocalAMP” level, the evolutionary classification is “Loss”. Otherwise, the evolutionary classification of amplifications is defined as “Indeterminate”.

### RNA-seq characterization of genomically perturbed cells under various drug conditions

We performed RNA-Seq on T47D cells perturbed to overexpress FGFR pathway activation including FGFR1, FGFR2 (WT, K660N, M538I and N550K), and FGF3, as well as GFP and parental as a control, as described above. Cells were plated in 96-well plates, and then treated with DMSO, fulvestrant (100 nM), palbociclib (1 μM), FIIN-3 (100 nM), and trametinib (500 nM) as single agent and in combinations for 24 hours. FGFR1/2 cell lines were treated with or without FGF2 (10 ng/mL) in various conditions. RNA was extracted from cells and sequencing libraries prepared as described in Supplementary Methods. For each specific construct and treatment combination we performed at least 6 replicates, for a total of 672 RNA-Seq profiles.

### Generation of plasmids and engineered cells

T47D or MCF7 cells were infected with lentivirus to derive stable cell lines overexpressing wildtype (WT) or mutant ORFs. All WT ORFs were obtained from the Broad Institute. Mutant ORFs (FGFR2 M538I, N550K and K660N) were made using QuickChange II site-directed mutagenesis kit (Agilent Technologies #200523). Most stable cells lines express ORFs in pLX317 vector and were selected with puromycin (Life Technologies #A1113803). Stable cell lines expressing CCND1 and PIM1 in pLX304 vector were selected with blasticidin (Life Technologies #A1113903).

For inducible cell lines, WT or Mutant FGFR2 IIIb ORFs were cloned into pInducer20 Tetracycline-inducible lentiviral vector (Addgene #44012) using gateway cloning technology (Invitrogen #11791019). Lentivirus was then generated and infected to establish Tet-inducible cells lines cultured in 10% Tet-system approved FBS (Clontech #631106) and 500 μg/mL G418 (Life Technologies #10131035). Doxycycline (Clontech #631311) was used to induce ORF expression.

### Kill curves and CellTiter-Glo viability assay

Cells were plated in 96-well tissue culture ViewPlate (Perkin Elmer # 6005181) on Day 1 and treated with drug on Day 2. Media with or without drugs was refreshed on Day 5. On Day 8, cells were equilibrated to room temperature, media was removed, and cells were lysed in a mixture of 50 µL media and 50 µL CellTiter-Glo 2.0 reagent (Promega # G9243) per well. Plates were then incubated on an orbital shaker for 2 mins. Following another 10 mins of incubation at room temperature to stabilize signal, luminescence was recorded to measure cell viability on Infinite M200 Pro microplate reader (Tecan).

### Colony formation assay

2,000-30,000 cells were plated in 6-well plates on Day 1 and treated on Day 2. Media was refreshed every 3-4 days until crystal violet staining. On the day of staining, cells were fixed with ice-cold 100% methanol for 10 minutes and then incubated with 0.5% crystal violet solution (Sigma Aldrich #C6158) in 25% methanol at room temperature for 10 minutes.

### Western blotting

Western blotting was performed as described in supplemental methods.

### Statistical analysis

Statistical analyses related to drug response curve were performed with two-tailed student t-test in Graphpad Prism. Fisher’s exact test was used to calculate odds ratio and q-value for volcano plots in the RNA sequencing analysis. Cohen’s D test with Hedges correction and Welch’s t-test were used to estimate the effect size and significance for signature strength of gene sets.

## Data availability

Tumor and germline whole exome sequencing data generated and analyzed for this study will be deposited in the access-controlled public repository dbGAP (https://www.ncbi.nlm.nih.gov/gap). Additional data generated in this study including tumor exome analysis and RNA-seq data are available within the paper and in the supplementary information files.

## Acknowledgements

We thank Qaren Quartey, Christian Kapstad and Gabriela Johnson for technical assistance; Karla Helvie, Laura Dellostritto, Lori Marini, Nelly Oliver, Shreevidya Periyasamy, Colin Mackichan, Max Lloyd, and Mahmoud Charif for assistance with patient sample collection and annotation; and Flora Luo, Tinghu Zhang and Nathanael Gray for providing reagents. We thank Jorge Gómez Tejeda Zañudo for helpful discussions and comments on the manuscript. We are grateful to all the patients who volunteered for our tumor biopsy protocol and generously provided the tissue analyzed in this study.

## Grant support

This work was supported by the Department of Defense W81XWH-13-1-0032 (N.W.), AACR Landon Foundation 13-60-27-WAGL (N.W.), NCI Breast Cancer SPORE at DF/HCC #P50CA168504 (N.W., N.U.L and E.P.W), Susan G. Komen CCR15333343 (N.W.), The V Foundation (N.W.), The Breast Cancer Alliance (N.W.), The Cancer Couch Foundation (N.W.), Twisted Pink (N.W.), Hope Scarves (N.W.), Breast Cancer Research Foundation (N.U.L. and E.P.W.), ACT NOW (to Dana-Farber Cancer Institute Breast Oncology Program), Fashion Footwear Association of New York (to Dana-Farber Cancer Institute Breast Oncology Program), Friends of Dana-Farber Cancer Institute (to N.U.L.), Stand Up to Cancer (N.W.), National Science Foundation (N.W.), SU2C-TVF Convergence Scholarship (P.M.), and The American Association for Cancer Research Basic Science Fellowship (P.M.).

## Author contributions

P.M., O.C. and N.W. conceived and designed the study; P.M. K.K. and J.K. performed experiments; O.C. and J.B. performed the computational analyses with input from S.F.; P.E., S.A.W., A.G.W., and N.W. performed clinical data abstraction and annotation; P.M. and M.S.C performed the RNA sequencing experiments; O.R. helped with the execution of the RNA-Seq; A.R. helped with the RNA-Seq data analysis; U.N. performed the prior HER2 mutant RNA sequencing experiments; J.C., V.M., and M.C. provided the clinical and genomic data for the 3 patients in the Foundation Medicine cohort; F.P and D.R. helped with the design and execution of the ORF screen; E.P.W., N.U.L., and N.W. oversaw patient enrollment and sample collection on the metastatic biopsy protocol; P.M., O.C. and N.W. wrote the manuscript with input from all authors; N.W. supervised the study.

## Notes

#### Summary of Updates

We added Figure.7 to demonstrate FGFR/FGF-induced transcriptional changes through RNA-seq analysis and revealed several pathways including MAPK pathway were activated by FGFR signaling. Figure.9 was added to show that three oncogenic signatures induced by FGFR/FGF can be reversed by irreversible FGFR inhibitor and partially reversed by MEK inhibitor. Figure.10 was added to show that targeting MAPK pathway via SHP2 inhibition can overcome FGFR-induced endocrine resistance. New Supplemental Figure.S21 is related to Figure.10 and showed effect of SHP2 and MEK inhibitors on downstream signaling. Description of results from these figures are included in the "Results" section. We revised the "Method" and "Supplemental Methods" section to include RNA-seq related protocol and methods. In addition, Supplemental tables.9-13 are related to RNA-seq analysis and thus were included in the supplementary information. New authors were added to reflect the additional data we have added to the manuscript. We modified the statistical testing in figures where we have used student-t test to be more detailed. Lastly, we moved original Fig.7B to Supplemental Figure.S20.

