## Supplemental Data for "Acquired FGFR and FGF alterations confer resistance to estrogen receptor (ER) targeted therapy in ER+ metastatic breast cancer"

### SUPPLEMENTAL METHODS

#### *Cell culture*

293T, T47D and MCF7 cells were purchased from American Type Culture Collection (ATCC). 293T cells were cultured in DMEM (GIBCO #11995-065) with 10% fetal bovine serum (Gemini #100-119). T47D cells were cultured in Phenol Red-free RPMI-1640 (GIBCO #11835-030) with 10% FBS. MCF7 cells were cultured in Phenol Red-free MEM-α (GIBCO #41061-029) with 10% FBS.

#### *Chemical Reagents*

Fulvestrant was purchased from Sigma Aldrich (# I4409); SHP099 (#406794) and GDC-0810 (#206041) were purchased from MedKoo BioSciences. Recombinant human FGF2 (#233FB025), FGF3 (#1206F3025), FGF6 (#238F6025), FGF10 (#345FG025) and FGF22 (#3867FG025) were purchased from R&D Systems. Heparin (#H3149) was purchased from Sigma Aldrich. PD173074 was purchased from StemCell Technologies Inc. (#72162) and Selleck Chemicals (#S1264). Palbociclib (#S1116), dovitinib (#S1018), ponatinib (#S1490) and everolimus (# S1120) were purchased from Selleck Chemicals. Trametinib (#NC0370092), AZD5363 (#NC0488926), AZD4547 (#NC0660421) were purchased from Thermo Fisher Scientific. FIIN-2 and FIIN-3 compounds were obtained from the laboratory of Dr. Nathanael Gray.

#### *Secondary validation screen*

A smaller pooled lentiviral library consisting of 570 ORFs was generated to confirm candidate resistance genes identified in the primary ORF screen. This library consists of 121 candidate resistance genes (ORFs that were enriched after selection by fulvestrant or GDC-0810 with a Z-score > 3), 344 negative control ORFs, and 95 candidate sensitizing genes (ORFs that were depleted after drug selection) to mimic the distribution of ORF composition in the genome-scale ORF library. In addition, we included several ORFs that were not in the primary ORF screen.

The secondary validation screen was conducted in both T47D and MCF7 cell lines using a similar protocol as the primary screen. Cells were infected with pooled virus and treated with 100 nM fulvestrant or GDC-0810 (1 μM for T47D cells and 100 nM for MCF7 cells) for three weeks. The dose of GDC-0810 used in MCF7 cells was reduced due to higher drug sensitivity. Cells were harvested for sequencing to determine the representation of ORFs every week during the 3-week drug treatment to monitor the dynamics of ORF representation.

***Whole Exome Sequencing***

DNA was extracted from primary tumors, metastatic tumors, and peripheral blood mononuclear cells (for germline DNA) from all patients and whole exome sequencing was performed, as detailed below. In several instances, cell free DNA was obtained from plasma for circulating tumor DNA analysis, as previously described[1].

*DNA extraction:* DNA extraction was performed as previously described[2]. For whole blood, DNA is extracted using magnetic bead-based chemistry in conjunction with the Chemagic MSM I instrument manufactured by Perkin Elmer. Following red blood cell lysis, magnetic beads bind to the DNA and are removed from solution using electromagnetized rods. Several wash steps follow to eliminate cell debris and protein residue from DNA bound to the magnetic beads. DNA is then eluted in TE buffer. For frozen tumor tissue, DNA and RNA are extracted simultaneously from a single frozen tissue or cell pellet sample using the AllPrep DNA/RNA kit (Qiagen). For FFPE tumor tissues, DNA and RNA are extracted simultaneously using Qiagen's AllPrep DNA/RNA FFPE kit. All DNA is quantified using Picogreen

*Library Construction:* DNA libraries for massively parallel sequencing were generated as previously described[2] with the following modifications: the initial genomic DNA input into the shearing step was reduced from 3µg to 10-100ng in 50µL of solution. For adapter ligation, Illumina paired-end adapters were replaced with palindromic forked adapters (purchased from Integrated DNA Technologies) with unique dual indexed 8 base index molecular barcode sequences included in the adapter sequence to facilitate downstream pooling. With the exception of the palindromic forked adapters, all reagents used for end repair, A-base addition, adapter ligation, and library enrichment PCR were purchased from KAPA Biosciences in 96-reaction kits. In addition, during the post-enrichment solid phase reversible immobilization (SPRI) bead cleanup, elution volume was reduced to 30µL to maximize library concentration, and a vortexing step was added to maximize the amount of template eluted.

*Solution-phase hybrid selection*: After library construction, hybridization and capture were performed using the relevant components of Illumina's Rapid Capture Exome Kit and following the manufacturer's suggested protocol, with the following exceptions: first, all libraries within a library construction plate were pooled prior to hybridization. Second, the Midi plate from Illumina's Rapid Capture Exome kit was replaced with a skirted PCR plate to facilitate automation. All hybridization and capture steps were automated on the Agilent Bravo liquid handling system.

*Preparation of libraries for cluster amplification and sequencing:* After post-capture enrichment, library pools were then quantified using quantitative PCR (KAPA Biosystems) with probes specific to the ends of the adapters; this assay was automated using Agilent's Bravo liquid handling platform. Based on qPCR quantification, libraries were normalized and denatured using 0.1 N NaOH on the Hamilton Starlet.

*Cluster amplification and sequencing:* Cluster amplification of denatured templates was performed according to the manufacturer's protocol (Illumina) using HiSeq 2500 Rapid Run v1/v2, HiSeq 2500 High Output v4 or HiSeq 4000 v1 cluster chemistry and HiSeq 2500 (Rapid or High Output) or HiSeq 4000 flowcells. Flowcells were sequenced on HiSeq 2500 using v1 (Rapid Run flowcells) or v4 (High Output flowcells) Sequencing-by-Synthesis chemistry or v1 Sequencing-by-Synthesis chemistry for HiSeq 4000 flowcells. The flowcells were then analyzed using RTA v.1.18.64 or later. Each pool of whole exome libraries was run on paired 76np runs, with a two 8 base index sequencing reads to identify molecular indices, across the number of lanes needed to meet coverage for all libraries in the pool.

*Sequence data processing:* Exome sequence data processing was performed using established analytical pipelines at the Broad Institute.  A BAM file was produced with the Picard pipeline (see URLs) which aligns the tumor and normal sequences to the hg19 human genome build using Illumina sequencing reads. The BAM was uploaded into the Firehose pipeline (see URLs), which manages input and output files to be executed by GenePattern[3].

*Sequencing quality control:* Quality control modules within Firehose were applied to all sequencing data for comparison of the origin for tumor and normal genotypes and to assess fingerprinting concordance. Cross-contamination of samples was estimated using ContEst[4].

***Somatic Alteration Assessment***

MuTect[5] was applied to identify somatic single-nucleotide variants. Indelocator (see URLs), Strelka[6], and MuTect2 (see URLs) were applied to identify small insertions or deletions. A voting scheme with inferred indels requiring at least 2 out of 3 algorithms.

Artifacts introduced by DNA oxidation (so called OxoG) during sequencing were computationally removed using a filter-based method[7]. In the analysis of primary tumors that are formalin-fixed, paraffin-embedded samples [FFPE] we further applied a filter to remove FFPE-related artifacts[8].

Reads around mutated sites were realigned with Novoalign (see URLs) to filter out false positive that are due to regions of low reliability in the reads alignment. At the last step, we filtered mutations that are present in a comprehensive WES panel of 8,334 normal samples (using the Agilent technology for WES capture) aiming to filter either germline sites or recurrent artifactual sites. We further used a smaller WES panel of normal 355 normal samples that are based on Illumina technology for WES capture, and another panel of 140 normals sequenced within our cohort[9] to further capture possible batch-specific artifacts. Annotation of identified variants was done using Oncotator[10].

#### *Copy Number and Copy Ratio Analysis*

To infer somatic copy number from WES, we used ReCapSeg (see URLs), calculating proportional coverage for each target region (i.e., reads in the target/total reads) followed by segment normalization using the median coverage in a panel of normal samples. The resulting copy ratios were segmented using the circular binary segmentation algorithm[11].

To infer allele-specific copy ratios, we mapped all germline heterozygous sites in the germline normal sample using GATK Haplotype Caller[12] and then evaluated the read counts at the germline heterozygous sites in order to assess the copy profile of each homologous chromosome. The allele-specific copy profiles were segmented to produce allele specific copy ratios.

#### *Cancer Cell Fraction and Evolutionary Analysis*

*Analysis using ABSOLUTE*: To properly compare SNVs and indels in paired metastatic and primary samples, we considered the union of all mutations called in either of the two samples. We evaluated the reference and alternate reads in each patient's primary and metastatic tumors, including mutations that were not initially called in one of the samples. These mutations in matched samples were used as input for ABSOLUTE[13]. The ABSOLUTE algorithm uses mutation-specific variant allele fractions (VAF) together with the computed purity, ploidy, and segment-specific allelic copy-ratio to compute cancer cell fractions (CCFs).

*Analysis of clonal dynamics using PHYLOGIC*: To evaluate the mutation clonality in the patient-matched primary and metastatic samples, we used PHYLOGIC clustering of the mutation-specific cancer cell fractions (CCFs), as previously described[14,15]. Further details regarding CCF and Evolutionary analysis can be found online (see URLs).

#### *Evolutionary analysis of copy-number variation*

*Corrected quantification of copy number:* gene amplifications are based on the purity corrected measure for the segment containing that gene, based on ABSOLUTE[13] (rescaled_total_cn).

For the FGF3, FGF4, FGF19, CCND1 amplicon, due to lower reliability in the inference of genomic segmentation (inferred breakpoints and amplicon inclusion), we manually reviewed the segments and their exome probes. This resulted in two cases whose FGF3 amplicon was re-defined as "Indeterminate" from "Acquired" due to lower support of the amplicon segmentation (ID 0300307, ID 0300338). For the same reason, we further re-defined CCND1 as "Indeterminate" from "Lost" for these three cases ID 0300177, ID 0300191, and ID 0300069, and FGF19 was re-defined as "Indeterminate" for ID0300069.

#### *ctDNA sequencing from Foundation Medicine cohort*

Hybrid capture-based genomic profiling was carried out on ctDNA patients with estrogen receptor-positive breast cancer. Peripheral blood samples were submitted by clinicians in the course of routine clinical care between May 2016 and March 2017. Sequencing of 62 genes was carried out to a median unique coverage depth of 7503×. Genomic alterations in ctDNA were evaluated and compared with matched tissue samples and genomic datasets of tissue from breast cancer.

#### ***Tissue sequencing from Foundation Medicine cohort***

Tumor samples from patients with ER-positive breast cancer were sequenced using a validated hybrid capture-based comprehensive genomic profiling (CGP) assay (FoundationOne) in a CLIA-certified, CAP-accredited, NY State-approved laboratory (Foundation Medicine, Cambridge MA) [16]. Samples underwent pathologist review to ensure sufficient tumor material (minimum of 20% tumor nuclei) and to confirm the pathologic diagnosis. At least 50 ng DNA was extracted from 40 μm of formalin-fixed, paraffin-embedded sections. CGP was performed on hybridization-captured, adaptor ligation-based libraries to a median coverage depth of 673× for up to 315 cancer-related genes plus select introns from up to 28 genes frequently rearranged in cancer. Results were analyzed for base substitutions, short insertions/deletions (indels), rearrangements, and copy number alterations (amplification and homozygous deletion). Custom filtering was applied to remove benign germline events as described [17]. Reportable genomic alterations (GAs) were called as known if the specific variant was present in the COSMIC database, if the variant has been characterized as pathogenic, or if the variant had likely functional status (disruptive alterations in tumor suppressor genes); all other variants were classified as variants of unknown significance (VUS). In this set 129/1,365 were HER2 positive (as defined by ERBB2 amplification on the FoundationOne assay)

#### ***Generation of Plasmids and Engineered Cell Lines***

T47D or MCF7 cells were infected with lentivirus to derive stable cell lines overexpressing wildtype (WT) or mutant ORFs. All WT ORFs were obtained from the Broad Institute. Mutant ORFs (FGFR2 M538I, N550K and K660N) were made in FGFR2 open reading frame in a pDORN221 vector backbone using QuickChange II site-directed mutagenesis kit (Agilent Technologies #200523). The mutant ORFs were then cloned into the pLX307 vector backbone using the gateway LR clonase kit (Invitrogen #11791019). The pLX317-GFP, pLX317-FGFR1, and pLX317-FGFR2 (WT) and pDONR221-FGFR2 (WT) plasmids were obtained from the Broad Institute. The VSV-G envelope and Δ8.91 packaging plasmids, and pLX307 vector were kind gifts from Levi Garraway.

For inducible cell lines, WT or Mutant FGFR2 IIIb ORFs were cloned into pInducer20 Tetracycline-inducible lentiviral vector (Addgene #44012) using gateway cloning technology (Invitrogen #11791019). Lentivirus was then generated using psPAX2 (kind gift from Dr. Adam Bass's laboratory) and VSV-G packaging plasmid and infected to establish Tet-inducible cells lines cultured in 10% Tet-system approved FBS (Clontech #631106) and 500 μg/mL G418 (Life

Technologies #10131035). Doxycycline (Clontech #631311) was used to induce ORF expression.

The primers used to make FGFR2 point mutations are as follows:

| Gene | Primer | Sequence |
|---|---|---|
| FGFR2 M538I | Forward primer (5'-3') | CCAATCATCTTCATTATCTCCATCTCTGACACCAGATCAGA |
| | Reverse primer (5'-3') | TCTGATCTGGTGTCAGAGATGGAGATAATGAAGATGATTGG |
| FGFR2 N550K | Forward primer (5'-3') | GCAGGCTCCAAGAAGTTTTATGATATTCTTGTGTTTCCCAATC |
| | Reverse primer (5'-3') | GATTGGGAAACACAAGAATATCATAAAACTTCTTGGAGCCTGC |
| FGFR2 K660N | Forward primer (5'-3') | CCGCCCATTGGTGGTGTTTTTGTAATAGTCTATATTGTTGATATC |
| | Reverse primer (5'-3') | GATATCAACAATATAGACTATTACAAAAACACCACCAATGGGCGG |

***Western blotting***

Cell lysates were prepared in RIPA buffer (Sigma Aldrich # R0278) supplemented with dithiothreitol (Life Technologies # 15508013), phenylmethane sulfonyl fluoride (Sigma Aldrich # P7626), protease inhibitor cocktail (Sigma Aldrich # P8340) and phosphatase inhibitor tablets (Roche # 4906845001). Mini-PROTEAN TGX 4-15% precast gel (Bio-rad # 4561084) or NuPAGE 4-12% Bis-Tris Midi protein gels (Invitrogen # WG1402A) were used for western blot. After electrophoresis, gel was transferred using Trans-Blot Turbo PVDF or nitrocellulose transfer pack (Bio-Rad # 1704158 and # 1704156). Membranes were probed with specific primary antibodies at 4 °C overnight and then incubated with corresponding HRP-conjugated secondary antibodies. We use Pierce ECL Plus Western Blotting Substrate (Thermo Fisher Scientific # 32132X3) as developing reagent.

Primary antibodies used in this study are: FGFR1 (Cell Signaling Technology CST #9740S), FGFR2 (Santa Cruz #6930), ERα (SC-543), p-ERK (CST #9106S), ERK (CST #4695S), p-AKT (CST #12694S), AKT (CST #9272S), p-S6K (CST #9205S), Actin (SC-1616-R), p-FRS2 (CST #3861S), GAPDH (SC-20357), CCND1 (CST #2978S), p-Rb (CST #9308S), Rb (CST #9309S). Secondary antibodies are from Invitrogen (anti-mouse A16090, anti-rabbit 32260, anti-goat 81-1620).

***Reverse transcription and quantitative PCR***

mRNA was extracted from cells with RNeasy Mini kit (Qiagen # 74104) and cDNA was made using High-Capacity cDNA Reverse Transcription Kit (Life Technologies # 4368814).

Quantitative-PCR was performed with SYBR Green master mix (Life Technologies # A25742) on 7500 Real-Time PCR system.

Primers:

| Gene | Primer | Sequence |
|---|---|---|
| CCND1 | Forward primer (5'-3') | CATCTACACCGACAACTCCATC |
| | Reverse primer (5'-3') | TCTGGCATTTTGGAGAGGAAG |
| GAPDH | Forward primer (5'-3') | ACATCGCTCAGACACCATG |
| | Reverse primer (5'-3') | TGTAGTTGAGGTCAATGAAGGG |

***Kinase array***

Proteome Profiler Human Phospho-Kinase Array Kit (R&D systems ARY003B) detects phosphorylation of 43 human kinases with antibodies spotted on nitrocellulose membrane that bind to target protein in samples. Proteome Profiler Human Phospho-MAPK Array Kit (R&D systems ARY002B) detects phosphorylation of 26 kinases in the MAPK pathway with capture antibodies spotted on membrane that bind to target proteins The signal of phosphorylation was detected with chemiluminescent detection reagents. Experiments were conducted according to protocol from manufacturer.

Kinases included in Human Phospho-Kinase Array Kit

| | | |
|---|---|---|
| Akt 1/2/3 (S473) | Hck (Y411) | PLC gamma-1 (Y783) |
| Akt 1/2/3 (T308) | HSP27 (S78/S82) | PRAS40 (T246) |
| AMPK alpha1 (T183) | HSP60 | Pyk2 (Y402) |
| AMPK alpha2 (T172) | JNK 1/2/3 (T183/Y185, T221/Y223) | RSK1/2/3 (S380) |
| beta-Catenin | Lck (Y394) | Src (Y419) |
| Chk-2 (T68) | Lyn (Y397) | STAT2 (Y689) |
| c-Jun (S63) | MSK1/2 (S376/S360) | STAT3 (S727) |
| CREB (S133) | p27 (T198) | STAT3 (Y705) |
| EGF R (Y1086) | p38 alpha (T180/Y182) | STAT5a (Y699) |
| eNOS (S1177) | p53 (S15) | STAT5a/b (Y699) |
| ERK1/2 (T202/Y204, T185/Y187) | p53 (S392) | STAT5b (Y699) |
| FAK (Y397) | p53 (S46) | STAT6 (Y641) |
| Fgr (Y412) | P70 S6 Kinase (T389) | TOR (S2448) |
| Fyn (Y420) | p70 S6 Kinase (T421/S424) | WNK-1 (T60) |
| GSK-3 alpha/beta (S21/S9) | PDGF R beta (Y751) | Yes (Y426) |

Kinases included in MAPK Array Kit

| | | |
|---|---|---|
| Akt1 | HSP27 | p38 beta |
| Akt2 | JNK1 | p38 delta |
| Akt3 | JNK2 | p38 gamma |
| Akt pan | JNK3 | p53 |
| CREB | JNK pan | p70 S6K |
| ERK1 | MKK3 | RSK1 |
| ERK2 | MKK6 | RSK2 |
| GSK-3 alpha/beta | MSK2 | TOR |
| GSK-3 beta | p38 alpha | |

***CCND1 RNAi***

CCND1 siRNAs were purchased from Origene (SR300410A, SR300410B, and SR300410C) was transfected using SITRAN 1.0 (Thermo Fisher Scientific). SR30004 was used as scrambled negative control siRNA.

| siRNA | Sequence |
|---|---|
| SR300410A | rCrCrArArUrArGrGrUrGrUrArGrGrArArArUrArGrCrGrCTG |
| SR300410B | rGrCrUrArUrGrGrArArGrUrUrGrCrArUrArArUrUrArUrUAT |
| SR300410C | rCrGrArUrUrUrCrArUrUrGrArArCrArCrUrUrCrCrUrCrUCC |

***Cell growth and treatment for RNA-seq experiments***

T47D cells were infected with lentivirus encoding FGFR pathway activation including FGFR1, FGFR2 (WT, K660N, M538I and N550K), and FGF3, as well as GFP in the pLX307 plasmid as described above. Following puromycin selection, cells were plated in 96-well plates, and then treated with DMSO, fulvestrant (100 nM), palbociclib (1 μM), FIIN-3 (100 nM), trametinib (500 nM), and the same concentrations in combinations - fulvestrant + FIIN-3, fulvestrant + trametinib, fulvestrant + palbociclib, palbociclib + FIIN-3, fulvestrant + palbociclib + FIIN-3, fulvestrant + palbociclib + trametinib. All drug treatments were given for 24 hours. The cells were washed 3X with ice-cold PBS and lysed with TCL buffer (*Qiagen 157013305*) containing 1% β-mercaptoethanol (*Sigma M3148*), transferred to PCR plates (*Qiagen 951020401*), sealed and immediately frozen at −80°C until RNA extraction. For each specific construct and drug condition we performed 6 biological replicates, for a total of 672 transcriptomes.

***RNA-seq***

Plates with cell lysates were thawed and purified with 2.2x RNAClean SPRI beads (Beckman Coulter Genomics). RNA captured beads were air-dried and processed immediately for RNA secondary structure denaturation (72˚C for three minutes) and cDNA synthesis. We used the SMART-Seq2 protocol[18] with minor modifications in the reverse transcription step. We made a 15μl reaction mix for each PCR and performed 10 cycles for cDNA amplification. We used 0.2ng cDNA of each population and one-eighth of the standard Illumina NexteraXT (Illumina FC-131-1096) reaction volume in both the tagmentation and PCR amplification steps. Uniquely indexed libraries were pooled and sequenced with NextSeq 500 high output V2 75 cycle kits (Illumina FC-404-2005) and 38 × 38 paired-end reads on an Illumina NextSeq 500 instrument, aggregating three NextSeq runs on a NextSeq 500 instrument.

***RNA-Seq data pre-processing and QC***

Reads were mapped to the human genome (hg19) with STAR aligner[19] with default parameters. Transcriptome quality and expression quantification was conducted using RNA-SeQC[20] to calculate Transcripts per Million (TPM) estimates. Samples with less than 7,500 unique genes were removed from subsequent analysis, excluding 25 samples and retaining 647 of 672 profiles, to a minimum of 4 replicates for each experimental condition. Samples passing QC had a mean of 13,004 detected genes by at least three reads (median = 13,139) and a mean of 5,411,989 uniquely mapped pairs of reads (median = 5,211,120).

***Data normalization and batch correction***

We merged and harmonized the RNA-seq data from our study with published RNA-seq data, using a similar approach to Nayar et. al[21]. Transcripts per million (TPM) were followed by log2 transformation. To correct for batch effects between studies, we used ComBat with default parameters[22].

***Linear Discriminant Analysis (LDA)***

Linear Discriminant Analysis (LDA) was performed with the R implementations of the function lda() as part of the "MASS" package, using the first ten Principal Components (PCs), using the prcomp() function in "stats" R package.

***Differential expression analysis***

Differential expression analysis was performed over the raw read counts using *limma* package [23] with *voom* assessment [23] of counts normalization and while accounting for the among-plates batch effect.

***Transcriptional gene-sets and signatures analysis***

We used Fast Gene Set Enrichment Analysis[24] to calculate enrichment for a given set of DEGs. Significant genes (q.value < 0.01) were ranked by logFC, with 100,000 permutations and gene set size limited to min of 3 and max of 1,000 genes, respectively. We additionally used Fisher's exact test (two-sided) to measure association between the each gene set in our compendium and the top 1,000 up-regulated genes in the experimental condition, based on differential expression analysis, ranked by Log(folds change). A total set of 5,150 gene sets was analyzed including the c2, c6, and hallmark collections from MSigDB [25] augmented with a breast cancer gene set collection[26-28].

For selected signaling pathways we had produced "Meta" gene sets that include a union of canonical (and partially overlapping) gene sets relevant to these pathways: (1) RTK/Growth Factor Receptors – union of EGFR_UP.V1_UP[29], PEDERSEN_METASTASIS_BY_ERBB2_ISOFORM_1, PEDERSEN_METASTASIS_BY_ERBB2_ISOFORM_2[30], ACEVEDO_FGFR1_TARGETS_IN_PROSTATE_CANCER_MODEL_UP[31], NAGASHIMA_EGF_SIGNALING_UP, NAGASHIMA_NRG1_SIGNALING_UP[32], resulting in 624 genes; (2) RAS/MAPK – union of RAS.ONCOGENE[33], RAS.ACT[34], RASERK.ACT[35] BILD_HRAS_ONCOGENIC_SIGNATURE[33], MEK_UP.V1_UP[29], HALLMARK_KRAS_SIGNALING_UP, resulting in 701 genes; (3) MOTR– union of MTOR_UP.V1_UP [36], HALLMARK_MTORC1_SIGNALING UP, KEGG_MTOR_SIGNALING_PATHWAY UP, PARENT_MTOR_SIGNALING_UP[37], BIOCARTA_MTOR_PATHWAY UP, resulting in 953 genes; (4) ER - union of HALLMARK_ESTROGEN_RESPONSE_LATE[25], HALLMARK_ESTROGEN_RESPONSE_EARLY[25],

YANG_BREAST_CANCER_ESR1_UP[38], VANTVEER_BREAST_CANCER_ESR1_UP[39], ESR1.DESMEDT[40], GOZGIT_ESR1_TARGETS_UP[41], resulting in 755 genes.

To evaluate signature strength (gene set score) for single samples we used AUCell [42] with 20% of top-ranking genes, using the log2 normalized, batch-corrected expression for each sample. For each gene set, the values were Z-score scaled across all 647 samples. For quantitative evaluation of signature strength differences between experimental conditions we used Cohen's D test with Hedges correction to estimate the effect size, and Welch's t-test to estimate significance.

#### *Statistical analysis*

Statistical analyses related to drug response curve were performed with student t-test in Graphpad Prism. Fisher's exact test was used to calculate odds ratio and q-value for volcano plots in the RNA-Seq analysis. Cohen's D test with Hedges correction and Welch's t-test were used to estimate the effect size and significance for signature strength for gene sets.

#### *URLs*

Picard (http://picard.sourceforge.net/); Firehose (http://www.broadinstitute.org/cancer/cga/Firehose); Indelocator (http://www.broadinstitute.org/cancer/cga/indelocator); MuTect2 (https://software.broadinstitute.org/gatk/documentation/tooldocs/current/org_broadinstitute_gatk_tools_walkers_cancer_m2_MuTect2); Novoalign (www.novocraft.com/products/novoalign/); ReCapSeg (http://gatkforums.broadinstitute.org/categories/recapseg-documentation); Oncotator (http://www.broadinstitute.org/cancer/cga/oncotator); CCF and Evolutionary analysis (http://www.broadinstitute.org/cancer/cga/acsbeta); RNA-SeQC (https://github.com/broadinstitute/rnaseqc).

### SUPPLEMENTAL TABLES

**Supplemental Table.1 (Excel File, 2 tabs)** A complete list of ORFs with a Z score > 3 in fulvestrant or GDC-0810 arm in the overexpression screen. The LFC values for all ORFs in the screen are also shown.

**Supplemental Table.2 (Excel File, 2 tabs)** Pathways nominated by GSEA analysis for resistance genes in fulvestrant or GDC-0810 arm.

| Breast Cancer Sample Type | No. of Pts | FGFR1 Amplification | | FGFR2 Amplification | | FGF3 Amplification | | Recurrent / Activating FGFR1 Mutation | | Recurrent / Activating FGFR2 Mutation | | Recurrent / Activating FGF3 Mutation | |
|---|---|---|---|---|---|---|---|---|---|---|---|---|---|
| | | No. | % | No. | % | No. | % | No. | % | No. | % | No. | % |
| This Study (Endocrine Resistant, ER+ MBC) | 60 | 9 | 15.0% | 3 | 5.0% | 17 | 28.3% | 0 | 0% | 2 | 3.33% | 0 | 0% |
| TCGA (Treatment-naive, ER+ primary breast cancer) | 739 | 90 | 12.2% | 7 | 0.9% | 122 | 16.5% | 6 | 0.81% | 3 | 0.4% | 1 | 0.1% |
| Enrichment Odds Ratio | | 1.3 | | **5.5** | | **2.0** | | 0 | | **8.318** | | 0 | |
| Enrichment p-value | | 0.32 | | **0.0331** | | **0.0199** | | 1 | | **0.049** | | 1 | |

**Supplemental Table.3** Comparison of the frequency of FGFR1/FGFR2/FGF3 alterations in the current study (endocrine-resistant ER+ metastatic breast cancer) versus in treatment-naive ER+ primary breast cancers in The Cancer Genome Atlas (TCGA PanCan Atlas; data version January 28 2016). Counts of amplifications were based on genes with CNAP of at least 3, using the same purity corrected inference in both cohorts. Counts of mutations were based on coding SNVs (silent mutations and mutations in introns, UTRs, and IGR are not included). In TCGA there were 11 coding SNVs in FGFR2, but only 3 of these were recurrent or “hotspots” and/or known activating mutations. The enrichment p.value was calculated used a one-sided fisher's exact test. No., number; Pts, patients.

**Supplemental Table.4 (Excel File, 1 tab)** Biopsy information including purity and ploidy for tumor samples across 24 patients with FGF/FGFR alterations.

**Supplemental Table.5 (Excel File, 3 tabs)** Complete exome and mutational information across all samples (SNVs and CNVs). For convenience, gene-level copy number for *FGFR1, FGFR2, FGF3, FGF4, FGF19*, and *CCND1* are shown in Tab 3.

**Supplemental Table.6 (Excel File, 2 tabs)** Evolutionary status and clonal fraction of SNVs and evolutionary status of CNVs across all samples.

**Supplemental Table.7 (Excel File, 3 tabs)** Complete clinicopathologic information on the 12 patients with acquired FGF or FGFR alterations, including patient information (tab 1), biopsy information (tab 2), and treatment and response information (tab 3).

**Supplemental Table.8 (Excel File, 1 tab)** FGFR alterations acquired in ER+ breast cancer patients from the Foundation Medicine (FM) cohort.

**Supplemental Table.9 (Excel File, 1 tab)** RNA-seq experiment conditions and QC stats.

**Supplemental Table.10 (Excel File, 5 tabs)** Gene sets defined and used in this study and signature-strength for these sets for each sample

**Supplemental Table.11 (Excel File, 8 tabs)** Differentially expressed genes.

**Supplemental Table.12 (Excel File, 4 tabs)** Gene sets associations and enrichment

**Supplemental Table.13 (Excel File, 1 tab)** Transcriptional signatures under drug treatment
