## Supplemental Figures for "Acquired FGFR and FGF alterations confer resistance to estrogen receptor (ER) targeted therapy in ER+ metastatic breast cancer"

Fig.S1

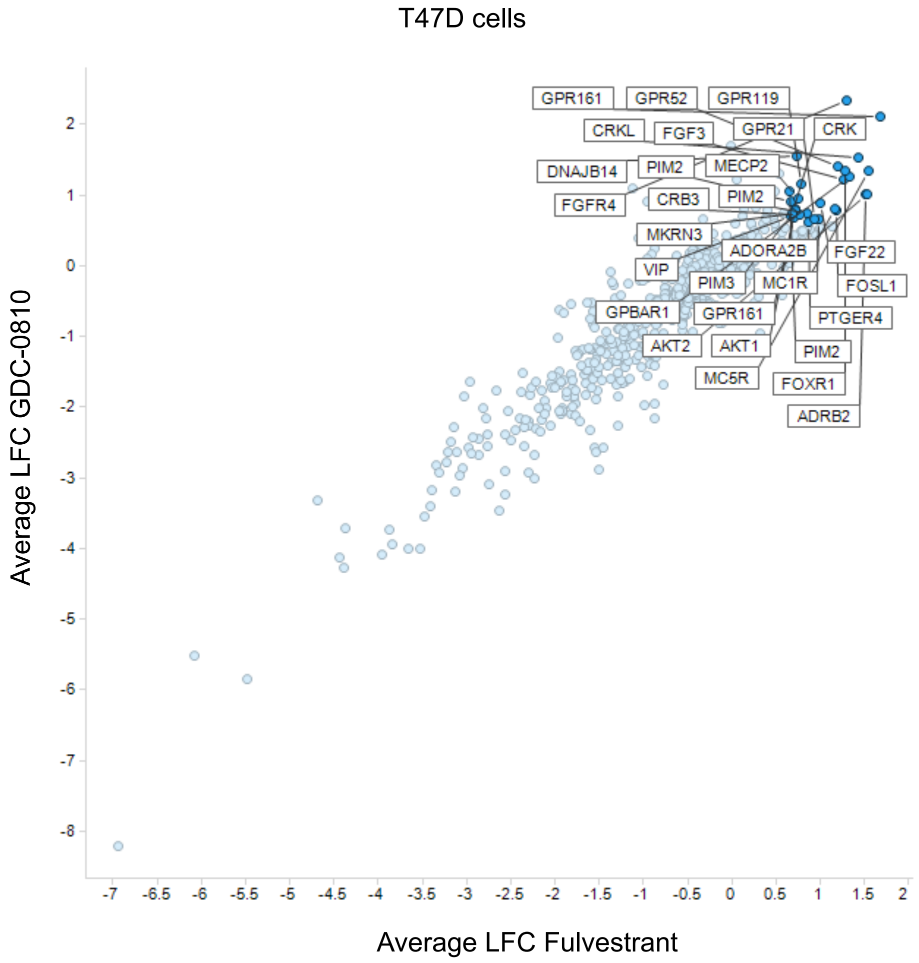


| Gene symbol | Pathway |
|---|---|
| GPR161 | G-protein coupled receptors |
| MC5R | G-protein coupled receptors |
| AKT2 | Serine/Threonine kinases |
| ADRB2 | G-protein coupled receptors |
| CRKL | Adaptor proteins |
| GPBAR1 | G-protein coupled receptors |
| FGFR4 | FGF/FGFR pathway |
| FOXR1 | Transcription factor |
| FGF3 | FGF/FGFR pathway |
| GPR52 | G-protein coupled receptors |
| FOSL1 | Transcription factor |
| ADORA2B | G-protein coupled receptors |
| FGF22 | FGF/FGFR pathway |
| MC1R | G-protein coupled receptors |
| GPR21 | G-protein coupled receptors |
| PTGER4 | G-protein coupled receptors |
| PIM2 | Serine/Threonine kinases |
| GPR119 | G-protein coupled receptors |
| CRB3 | Others |
| MECP2 | Others |
| MKRN3 | Others |
| DNAJB14 | Others |
| GPR161 | G-protein coupled receptors |
| PIM3 | Serine/Threonine kinases |
| AKT1 | Serine/Threonine kinases |
| CRK | Adaptor proteins |
| VIP | Others |
| PIM2 | Serine/Threonine kinases |
| PIM2 | Serine/Threonine kinases |

**Figure.S1 Resistance genes for fulvestrant and GDC-0810 identified in the secondary validation screen in T47D cells.** A, The secondary screen was conducted as described in the supplemental methods. The average LFC of each ORF was plotted for both the fulvestrant (X-axis) and GDC-0810 (Y-axis) arms. ORFs with an average LFC > 0.6 in both drug arms are labeled. The average LFC was calculated from three replicates in fulvestrant arm and two replicates for GDC-0810 arm (the third replicate for GDC-0810 failed sequencing). B, Labeled ORFs are ranked by LFC in the fulvestrant arm and categorized into functional pathways.

Fig.S2

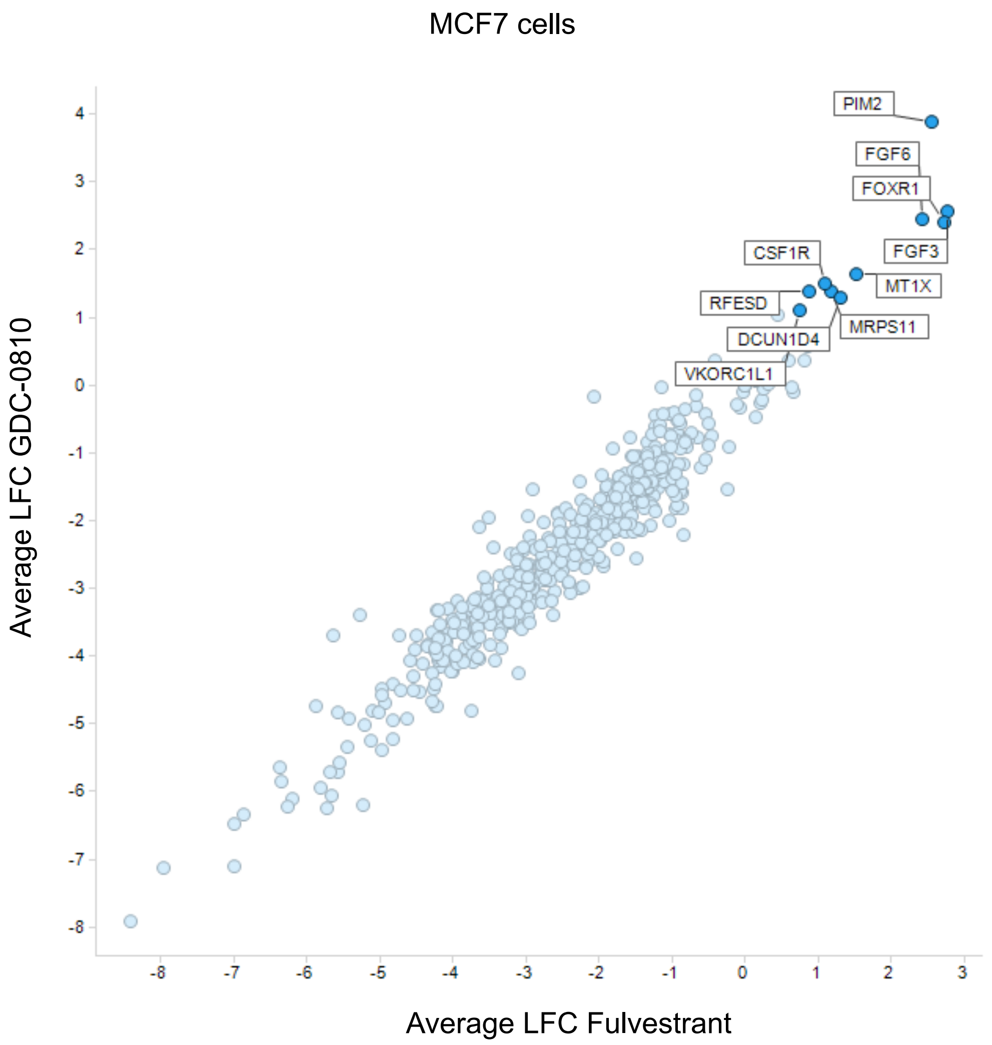


| Gene symbol | Pathway |
|---|---|
| FGF3 | FGF/FGFR pathway |
| FOXR1 | Transcription factor |
| PIM2 | Serine/Threonine Kinases |
| FGF6 | FGF/FGFR pathway |
| MT1X | Others |
| DCUN1D4 | Others |
| MRPS11 | Ribosomal protein |
| CSF1R | Receptor tyrosine kinase |
| RFESD | Others |
| VKORC1L1 | Others |

**Figure.S2 Resistance genes for fulvestrant and GDC-0810 identified in the secondary validation screen in MCF7 cells.** A, The secondary screen was conducted as described in the supplemental methods. The average LFC of each ORF was plotted for both the fulvestrant (X-axis) and GDC-0810 (Y-axis) arms. ORFs with an average LFC > 0.6 in both drug arms are labeled. The average LFC was calculated for three replicates in both drug arms. B, Labeled ORFs are ranked by LFC in the fulvestrant arm and categorized into functional pathways.

Fig.S3

T47D cells

A

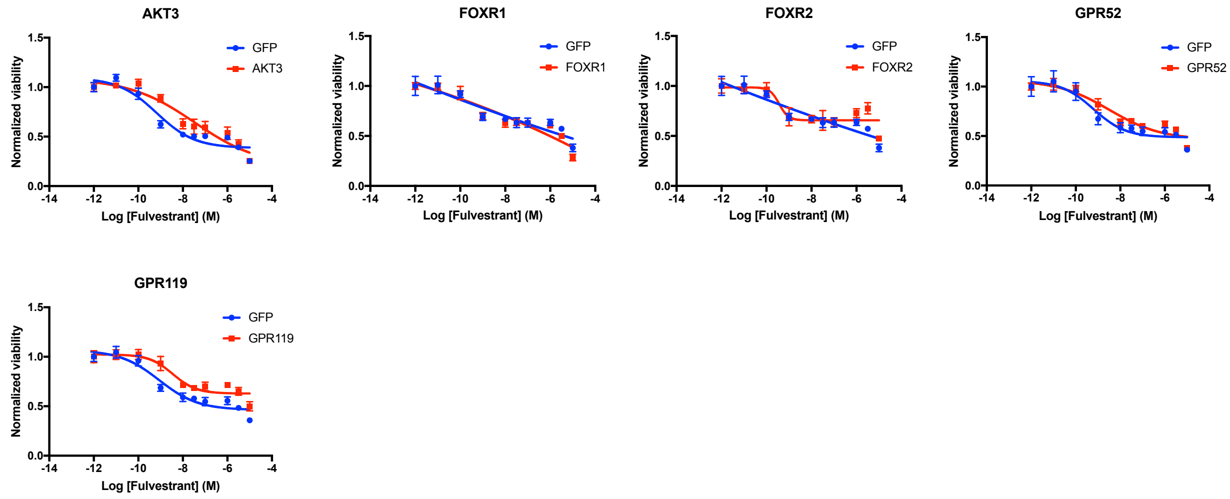


B

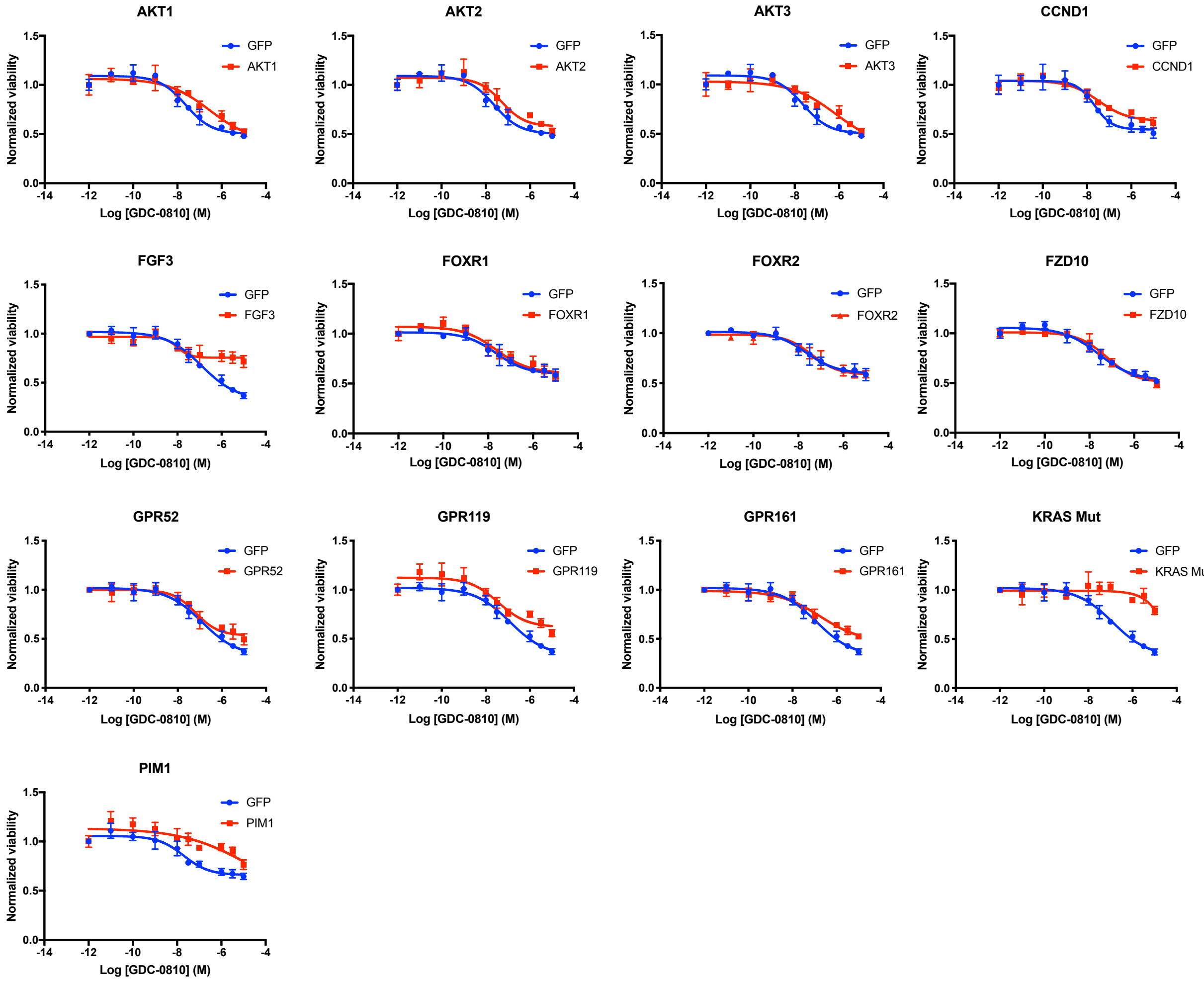


**Figure. S3 Additional genes were validated to confer resistance to fulvestrant and GDC-0810.** Individual genes were overexpressed in T47D cells and drug response to fulvestrant (A) and GDC-0810 (B) were examined in comparison to T47D cells expressing GFP. CellTiter-Glo assay was performed to measure cell viability and all the data points were normalized to DMSO conditions.

Fig.S4

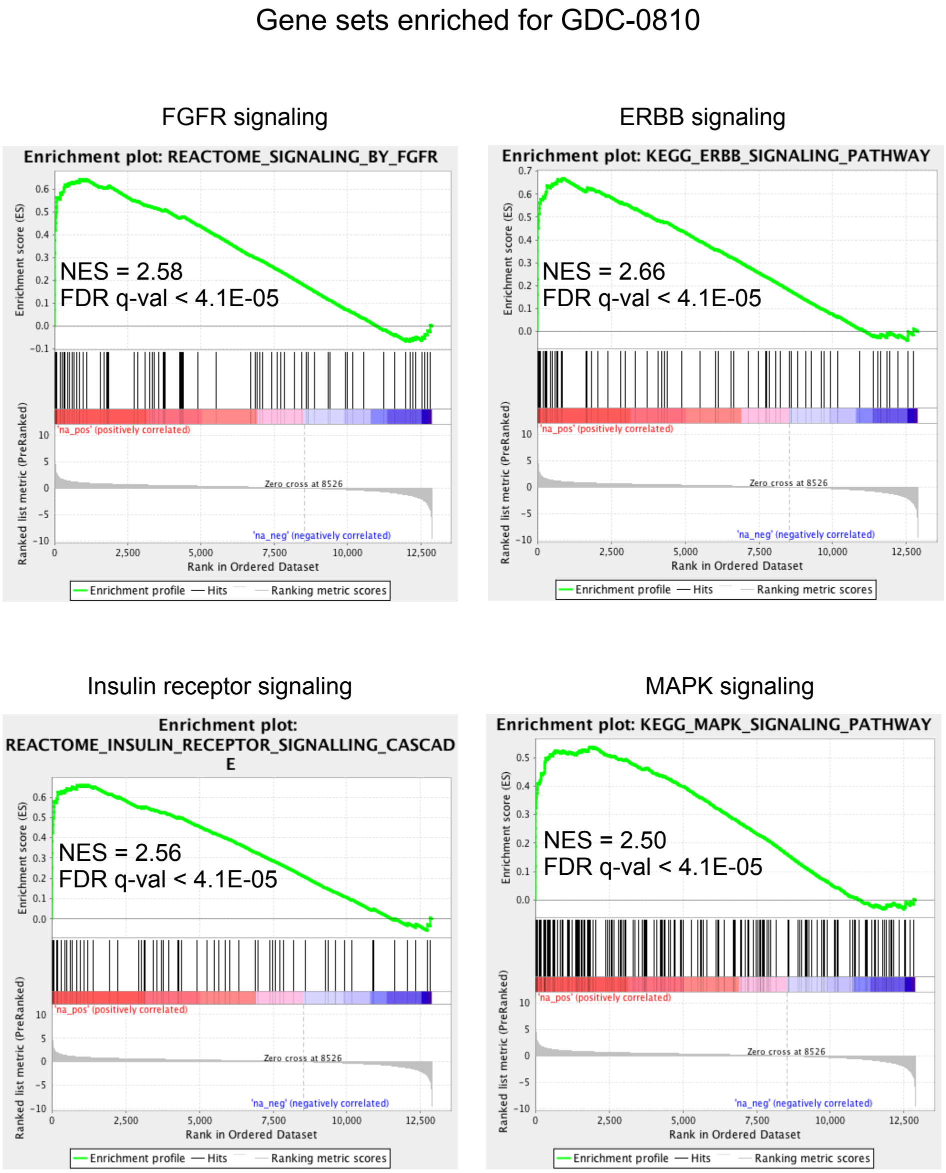


**Figure. S4 Resistance genes for GDC-0810 were enriched in similar oncogenic pathways as identified for fulvestrant.** GSEA was performed for the gene list ranked by LFC in the GDC-0810 arm. For genes with multiple ORFs in the library, the ORF with highest LFC was selected for ranking. NES, normalized enrichment score. Gene sets used are from c2.cp.v5.2.symbols.gmt and 1000 permutations were performed. Results for FGFR pathway, ERBB signaling, insulin receptor signaling and MAPK pathway are shown. A full list of nominated pathways is shown in Supplemental table.2.

Fig.S5

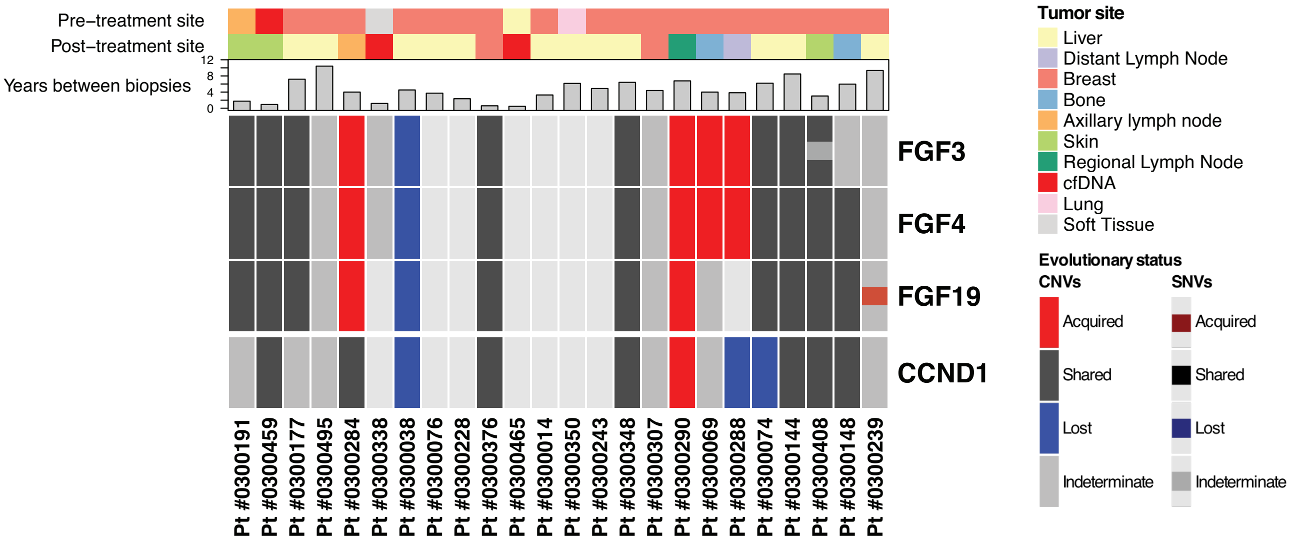


**Figure.S5 Evolutionary classification of the four proximal genes FGF3, FGF4, FGF19, and CCND1.** FGF3, FGF4, FGF19, and CCND1 are in immediate genomic proximity on Chr 11 (11q13.3) and are often co-amplified. In a subset of our patients, these genes do not have the same evolutionary classification, as they do not share the same amplicon in either the pre-treatment or the post-treatment sample. Further details about these amplicons are described in Supplemental Methods.

Fig.S6

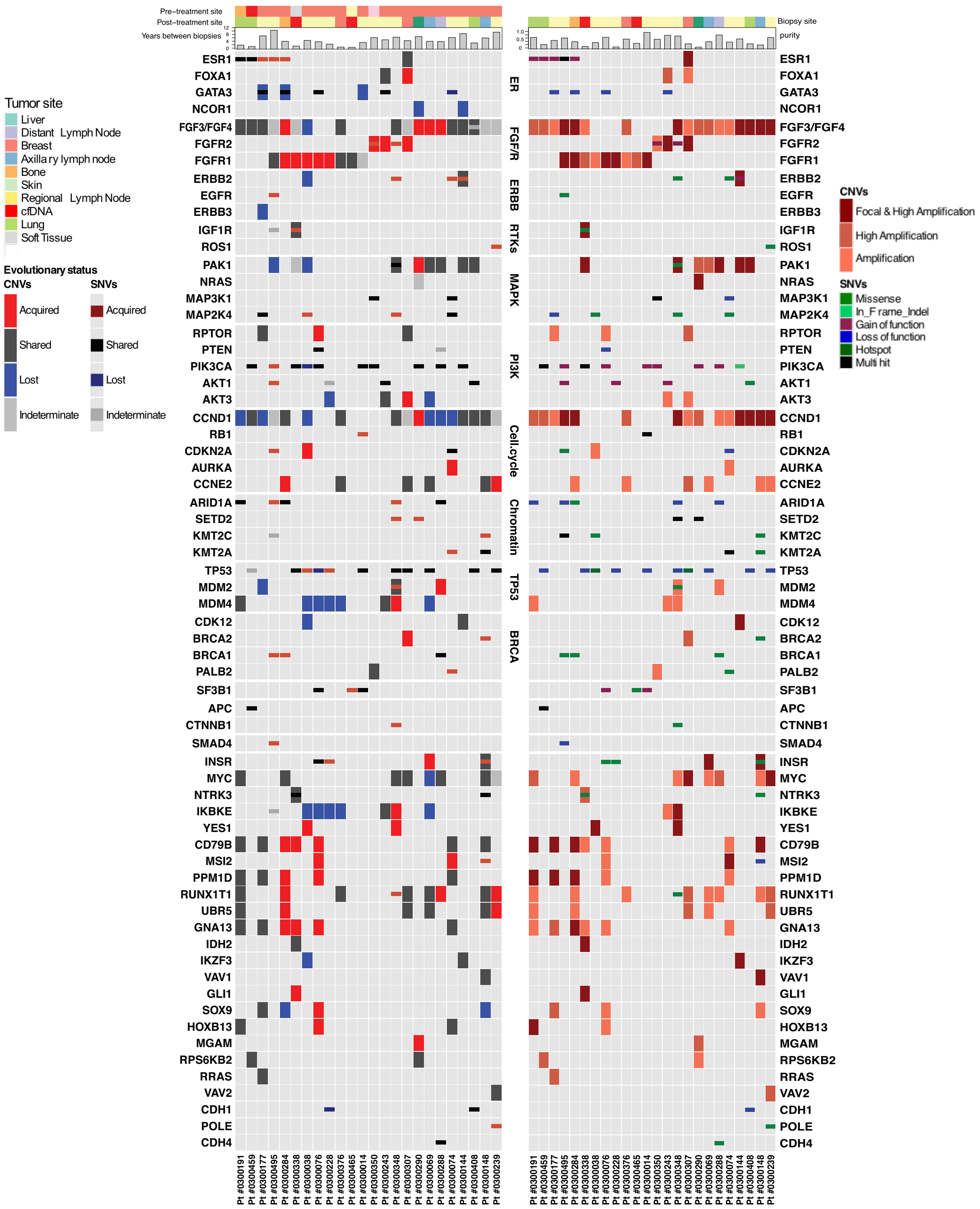

**Figure.S6 Evolutionary classification and mutational classification of key cancer genes in the post-treatment samples.** Left panel, evolutionary classification of selected cancer genes are shown. Alterations are presented as color-coded heatmap comparing the pre-treatment and post-treatment mutational status (red = acquired, blue = lost, black = shared, grey = indeterminate); Right panel, alteration types for the post-treatment samples. Notably, alterations that are denoted as lost or indeterminate in the left panel may not be observed in the post-treatment sample, thus not shown in the right panel. Denoted alterations include copy number and single nucleotide variation (SNV) of selected cancer genes in ER positive breast cancer. Alterations are presented with four levels of amplification – "Focal & High Amplification", "High Amplification", "Amplification" (in red spectrum colors), and various SNVs annotated as "Missense" and "In_Frame_Indel". Functional mutations are annotated as "Gain of function", "Loss of function", "Hotspot", and "Multi hit" if a gene is altered by multiple SNVs. Clinical and pathology tracks depict the site of biopsy for both matched samples.

Fig.S7

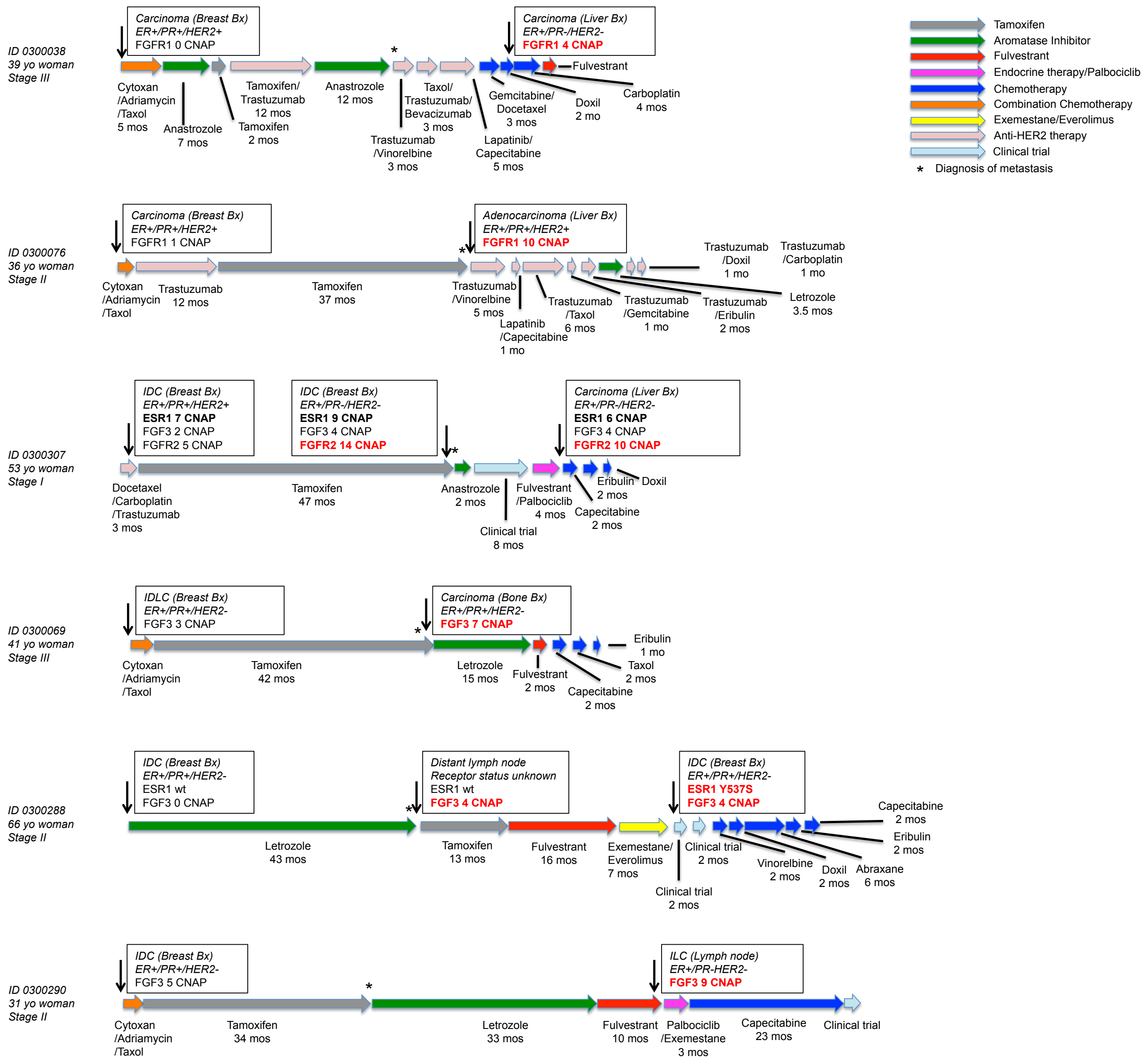


**Figure.S7 Additional clinical vignettes of patients who acquired FGFR or FGF alterations following endocrine therapy.** The clinical vignettes for selected patients with acquired alterations in FGFR1, FGFR2, and/or FGF3 illustrate detailed information on age and stage of disease at diagnosis, therapies patients received, duration of response to each therapy, and time of biopsies collected during the clinical course. For each biopsy, available information on biopsy type, tissue site, receptor status and selected genomic alterations detected by whole exome sequencing is shown. In each case, the asterisk indicates the time that metastatic disease was diagnosed. The complete clinicopathologic information for each patient is provided in Supplemental Table.7. IDC: invasive ductal carcinoma, IDLC: invasive ductal-lobular carcinoma; CNAP: copy number above ploidy; yo: years old; Bx: biopsy; PR: progesterone receptor; wt: wildtype.

Fig.S8

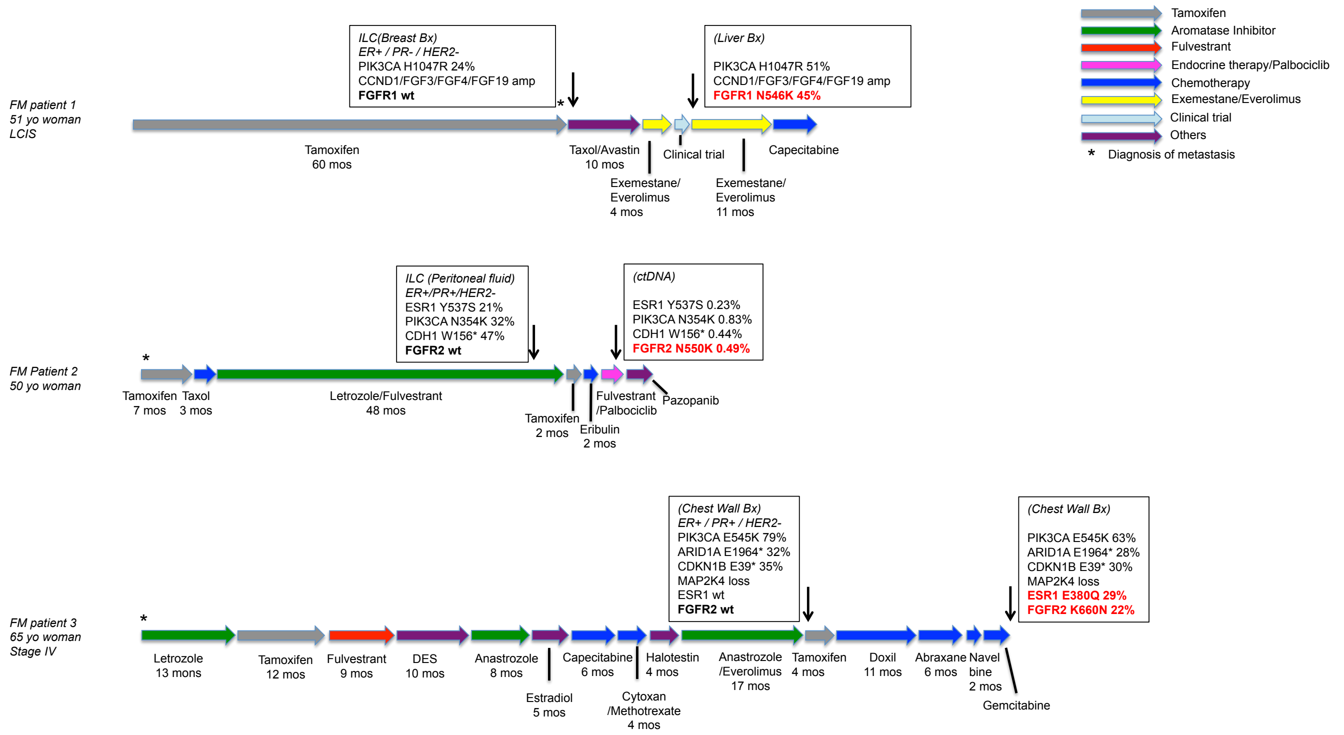


**Figure.S8 Additional clinical vignettes of patients who acquired FGFR alterations following endocrine therapy from the Foundation Medicine (FM) cohort.** The clinical vignettes for three patients with acquired alterations in FGFR1 or FGFR2 illustrate detailed information on age and stage of disease at diagnosis, therapies patients received, duration of response to each therapy, and time of biopsies collected during the clinical course. For each biopsy, available information on biopsy type, tissue site, receptor status and selected genomic alterations detected by panel sequencing is shown. In each case, the asterisk indicates the time that metastatic disease was diagnosed. ILC: invasive lobular carcinoma, LCIS: lobular carcinoma in situ, yo: years old; Bx: biopsy; PR: progesterone receptor; wt: wildtype. The allelic frequencies of selected alterations detected by panel sequencing are shown, the allelic frequencies of all alterations are revealed in Supplemental Table.8.

Fig.S9

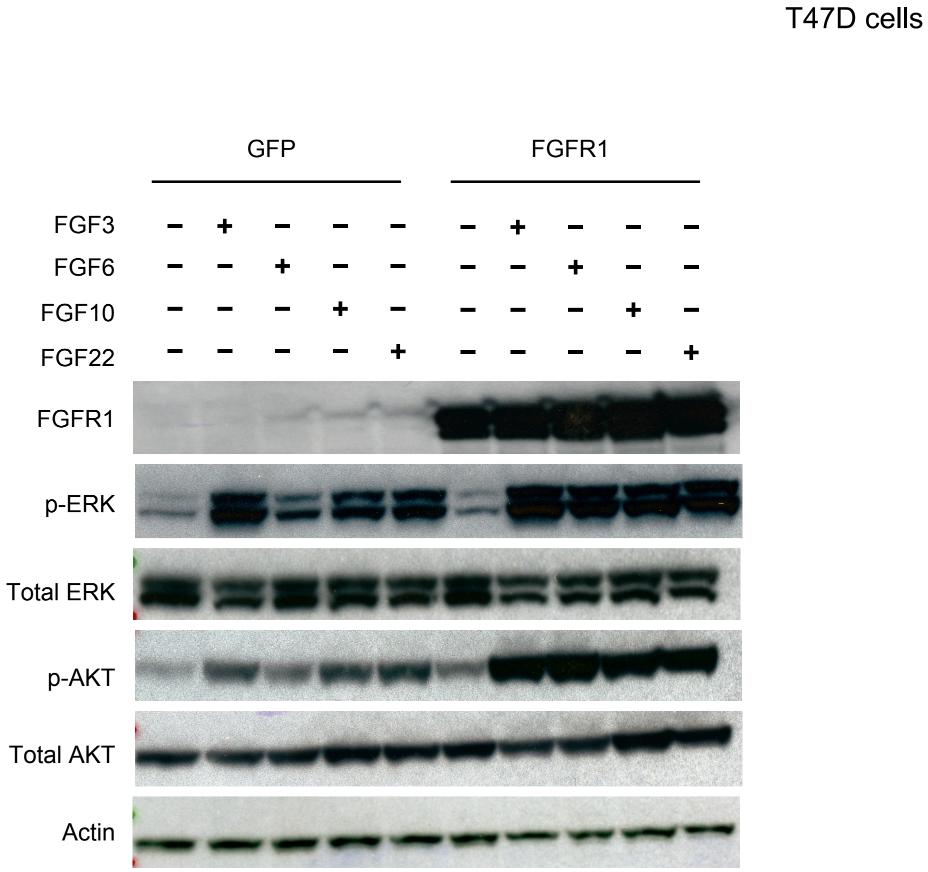


**Figure.S9 FGF ligands induced downstream AKT and ERK signaling, which was enhanced in the presence of FGFR1 expression.** T47D overexpressing GFP or FGFR1 cells were treated with various FGF ligands (100 ng/μL) for one hour with or without PD173074 (1 μM) before protein harvest for western blot.

Fig.S10

MCF7 cells

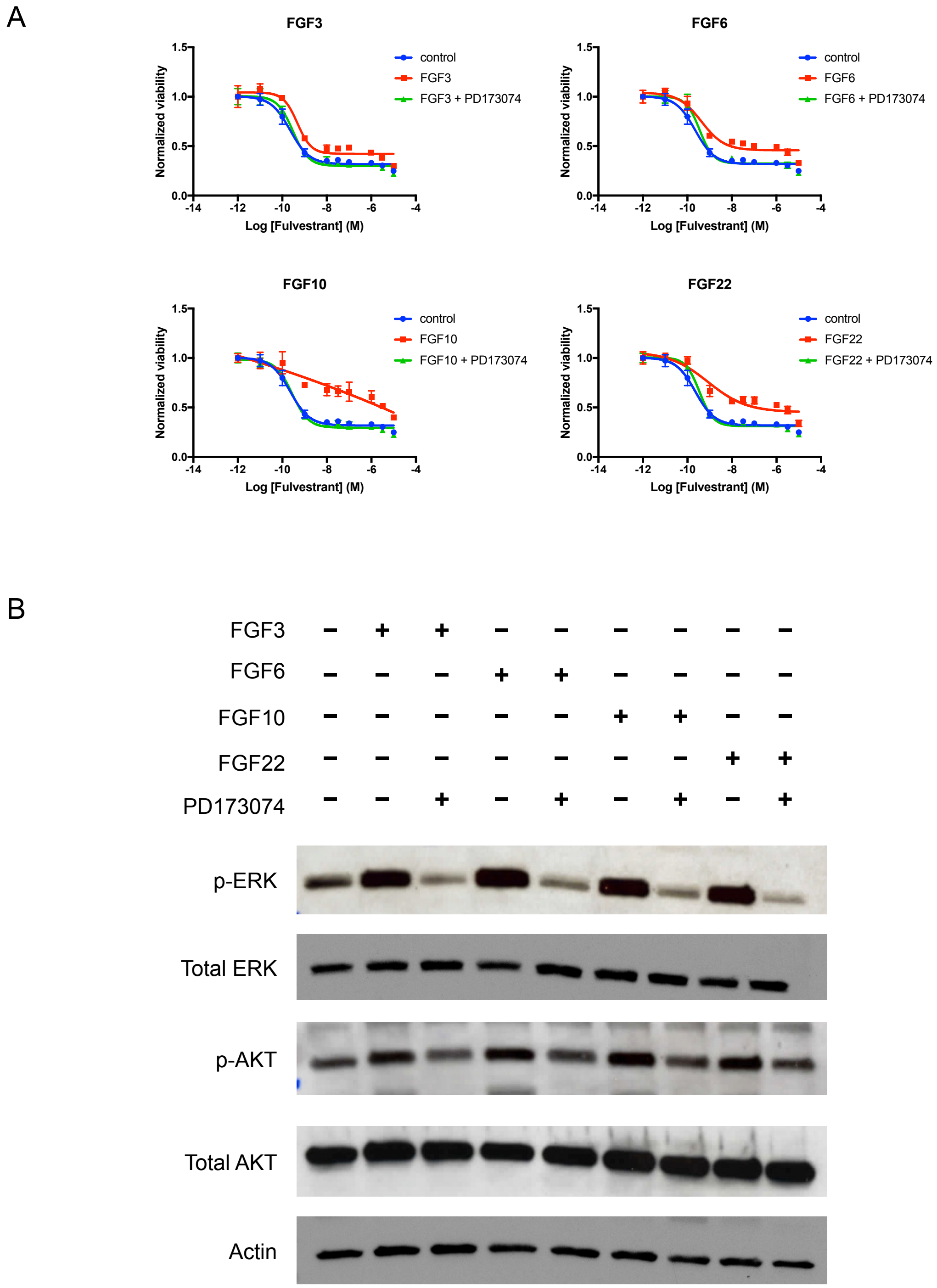


**Figure.S10 FGF ligands conferred resistance to fulvestrant in MCF7 cells, which was blocked by PD173074.** A, Recombinant FGF ligands were added into media every three days at the concentration of 100 ng/mL with or without 1 μM PD173014. Sensitivity to fulvestrant over six days was examined in MCF7 cells. B, FGF ligands increased ERK and AKT phosphorylation, which was blocked by PD173074. MCF7 cells were treated as indicated for one hour before protein harvest for western blot.

Fig.S11

T47D cells

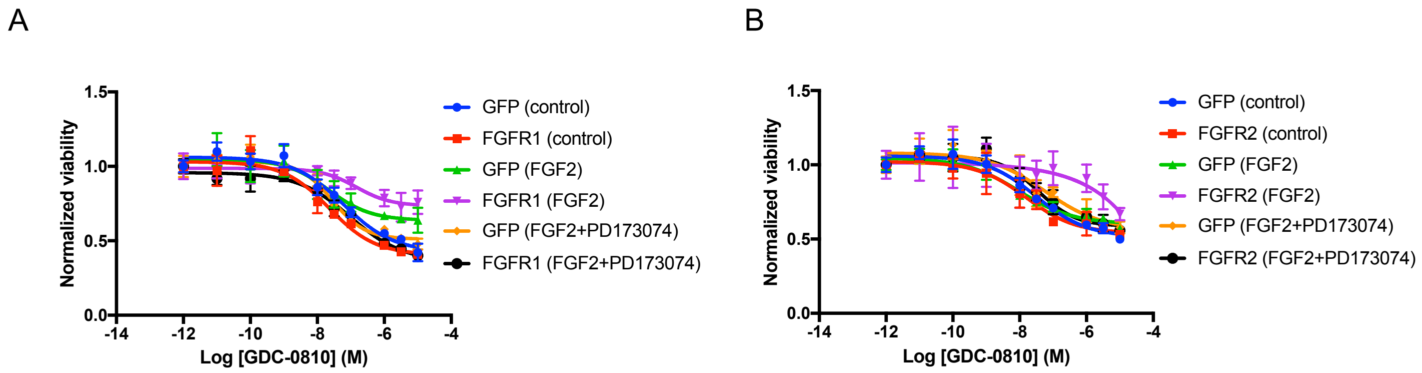


**Figure.S11 FGFR1/2 overexpression led to resistance to GDC-0810 in T47D cells, which was blocked by PD173074.** GFP, FGFR1 or FGFR2 was overexpressed in T47D cells to establish T47D_GFP, T47D_FGFR1 and T47D_FGFR2 cells. The GDC-0810 sensitivity of T47D_FGFR1 (A) and T47D_FGFR2 cells (B) was compared to T47D_GFP in the presence or absence of 10 ng/mL FGF2 and 1 μM PD173074 over six days of drug treatment.

Fig.S12

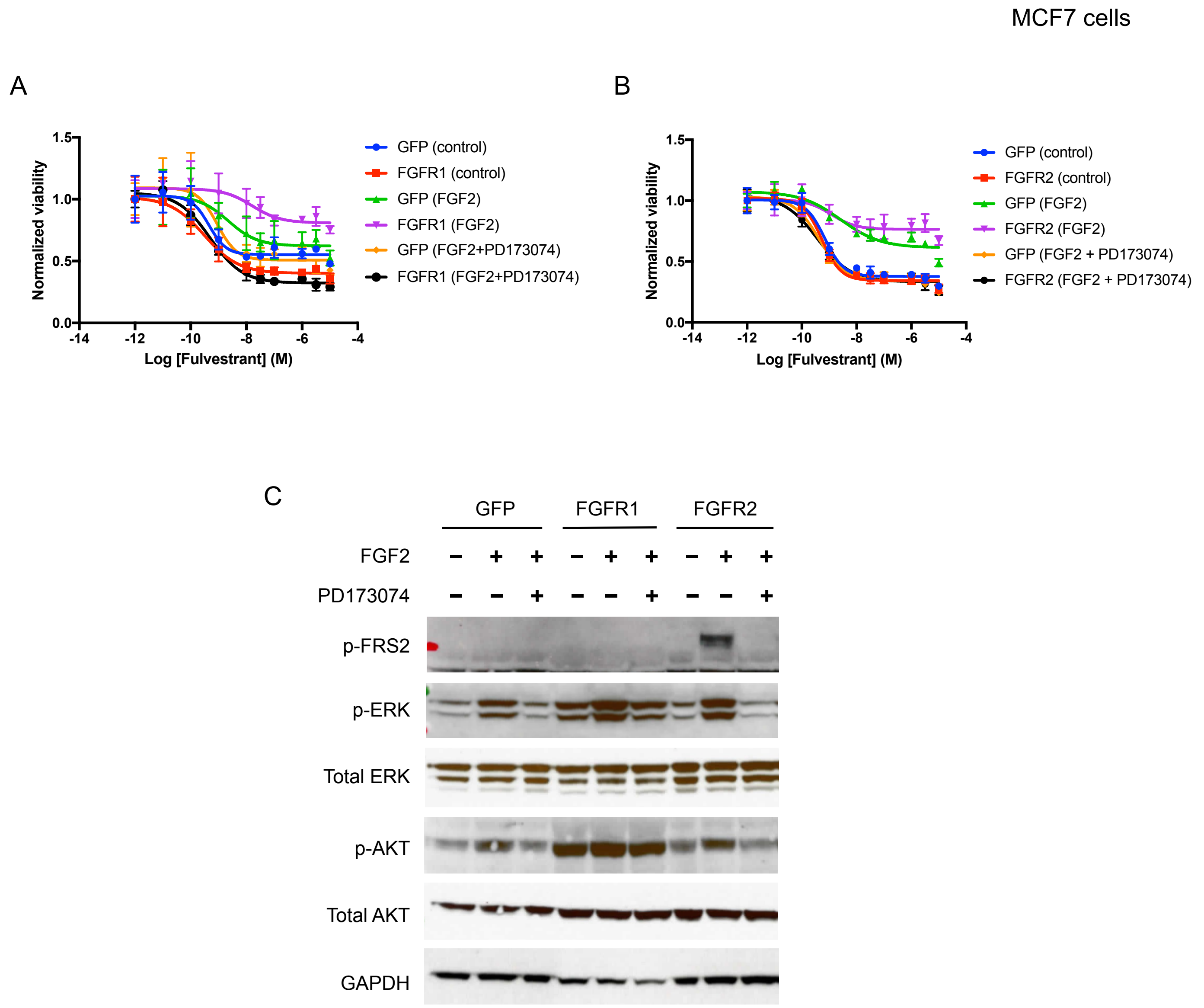


**Figure.S12 FGFR1/2 overexpression led to resistance to fulvestrant in MCF7 cells, which was blocked by PD173074.** GFP, FGFR1 or FGFR2 was overexpressed in MCF7 cells to establish MCF_GFP, MCF7_FGFR1 and MCF7_FGFR2 cells. The fulvestrant sensitivity of MCF7_FGFR1 (A) and MCF7_FGFR2 cells (B) was compared to MCF7_GFP in the presence or absence of 10 ng/mL FGF2 and 1 μM PD173074 over six days of drug treatment. (C) In MCF7 cells, FGFR1 and FGFR2 induced phosphorylation of ERK and AKT in the presence of FGF2, which was blocked by PD173074. Cells were treated with indicated conditions for one hour before protein harvest.

Fig.S13

A

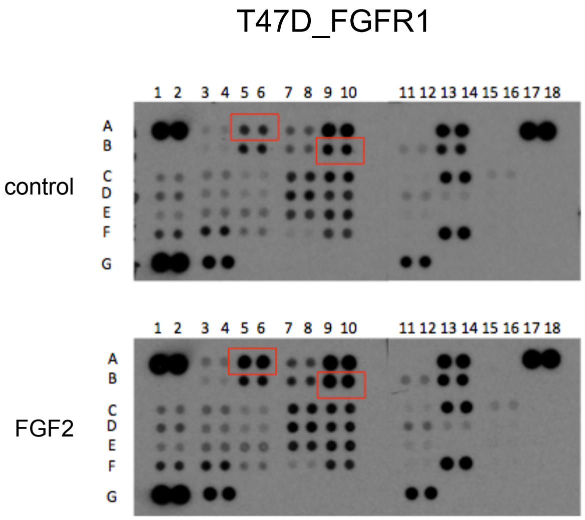


| | | |
|---|---|---|
| A5, A6 | ERK1/2 | T202/Y204, T185/Y187 |
| B9, B10 | Akt1/2/3 | S473 |

Kinase included in the array can be found: https://www.rndsystems.com/products/proteome-profiler-human-phospho-kinase-array-kit_ary003b

B

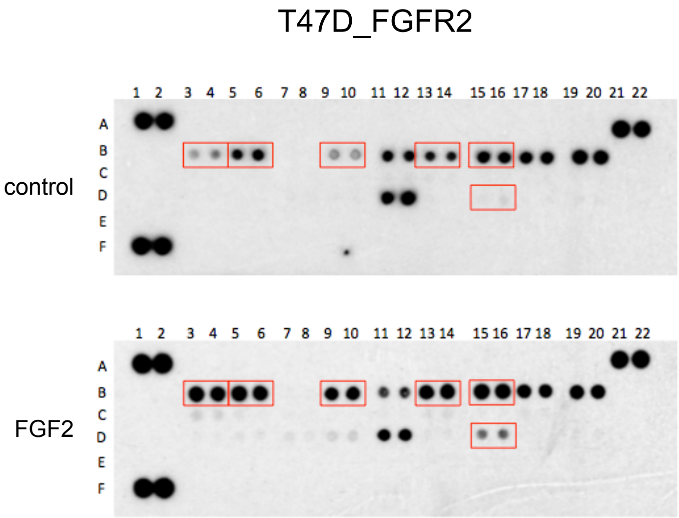


| | | |
|---|---|---|
| B3, B4 | Akt1 | S473 |
| B5, B6 | Akt2 | S474 |
| B9, B10 | Akt pan | S473, S474, S472 |
| B13, B14 | ERK1 | T202/Y204 |
| B15, B16 | ERK2 | T185/Y187 |
| D15, D16 | RSK1 | S380 |

Kinase included in the array can be found: https://www.rndsystems.com/products/proteome-profiler-human-phospho-mapk-array-kit_ary002b

**Figure.S13 Active FGFR signaling led to increased phosphorylation of AKT and ERK.** T47D_FGFR1 and T47D_FGFR2 cells were treated with or without FGF2 (10 ng/mL) daily for three days and harvested for protein extraction six hours after the last ligand treatment. For phosphorylation signal detection, 400 µg protein from T47D_FGFR1 cells was incubated with human kinase antibody array consisting of antibodies for 43 kinases, A1, A2, A17, A18, G1 and G2 are reference spots included to demonstrate the array has been incubated with Streptavidin-HRP during the assay procedure (A); 200 µg protein from T47D_FGFR2 cells was incubated with the MAPK pathway kinase array consisting of antibodies for 26 kinases, A1, A2, A21, A22, F1, and F2 are reference spots (B).

Fig.S14

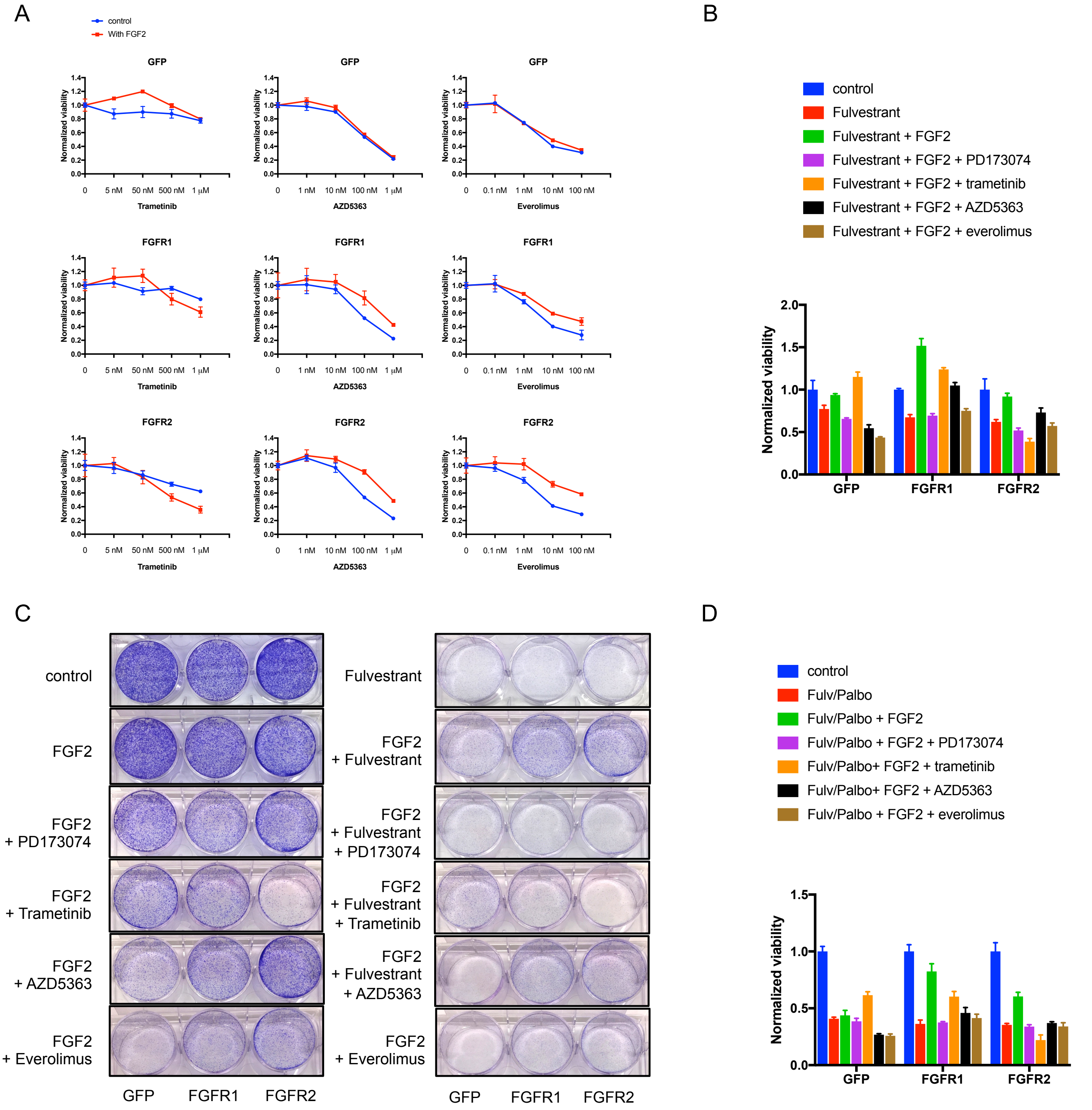


**Figure.S14 Active FGFR signaling led to hypersensitivity to MEK inhibitor trametinib and resistance to AKT inhibitor AZD5363 and mTOR inhibitor everolimus.** A, T47D_GFP, T47D_FGFR1 and T47D_FGFR2 cells were treated with various drugs to examine dose response in the presence or absence of FGF2. CellTiter-Glo assay was performed to measure cell viability following drug treatment. B, Everolimus abrogated resistance to fulvestrant conferred by FGFR1. Trametinib abrogated resistance to fulvestrant conferred by FGFR2. C, Trametinib is more effective to reduce colony formation induced by active FGFR signaling than AZD5363 and everolimus. 30,000 cells were plated on Day 1 and treated on Day 2, and colony formation assay was performed after 9 days of drug treatment D, Everolimus abrogated resistance to the combination of fulvestrant and palbociclib conferred by FGFR1. Trametinib abrogated resistance to the combination of fulvestrant and palbociclib conferred by FGFR2. Drug concentrations used in B-D: FGF2: 10 ng/mL, fulvestrant: 100 nM, palbociclib: 1 μM, PD173074: 1 μM, trametinib: 500 nM, AZD5363: 100 nM, everolimus: 10 nM.

Fig.S15

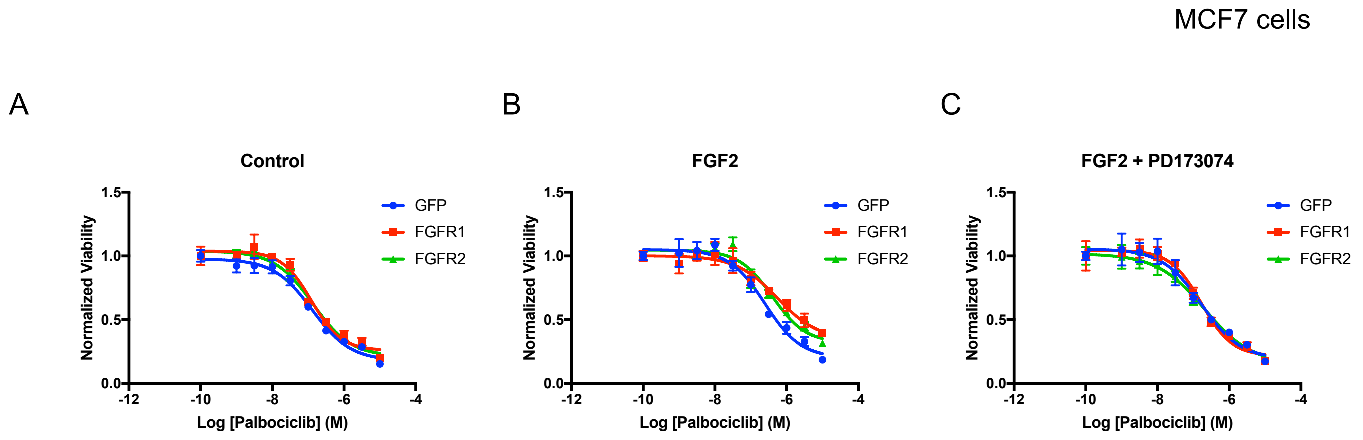


**Figure.S15 FGFR1/ 2 overexpression conferred moderate resistance to palbociclib in MCF7 cells.** GFP, FGFR1 or FGFR2 was overexpressed in MCF7 cells. The sensitivity of various cells lines to palbociclib was compared in the following conditions: control (A), 10 ng/mL FGF2 (B), 10 ng/mL FGF2 and 1 μM PD173074 (C), over six days of drug treatment.

Fig.S16

MCF7 cells

A

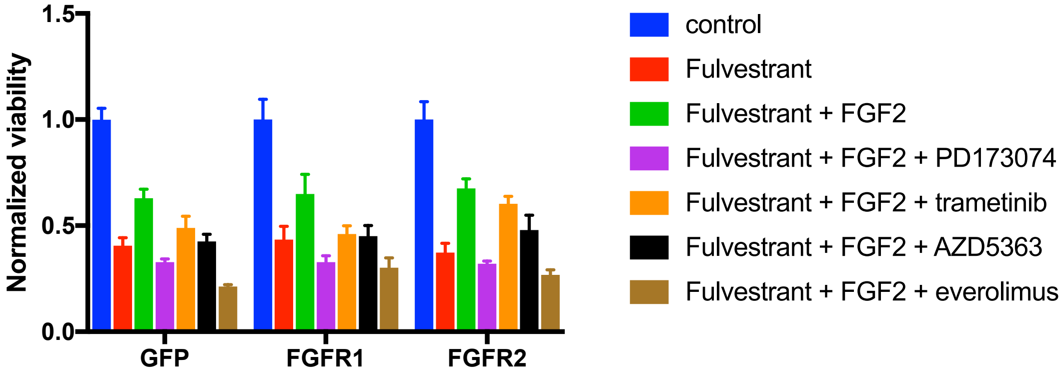


B

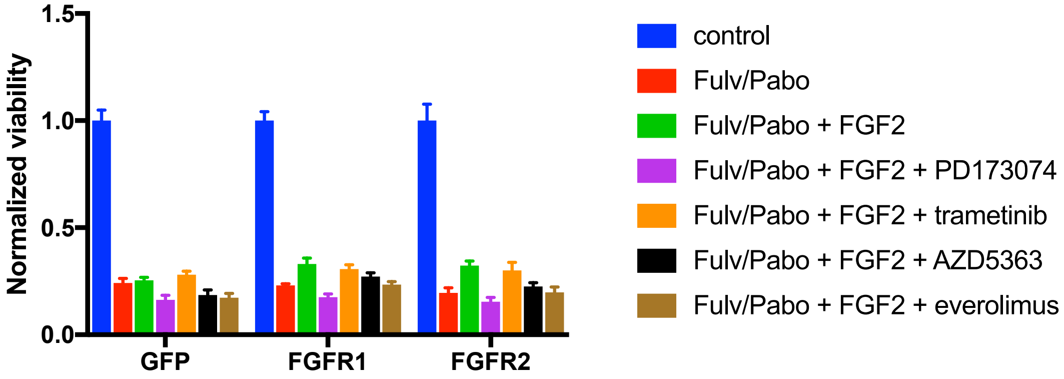


**Figure.S16 FGFR1 and FGFR2 conferred resistance to combination of fulvestrant and palbociclib in MCF7 cells.** Resistance to fulvestrant (A) and the combination of fulvestrant and palbociclib (B) conferred by FGFR1 or FGFR2 was blocked by PD173074 and everlolimus, and partially blocked by trametinib and AZD5363. Cells were treated with indicated conditions and cell viability was measured after six days. Concentrations of drugs used: FGF2: 10 ng/mL, fulvestrant: 100 nM, palbociclib: 1 µM, PD173074: 1 µM, trametinib: 50 nM, AZD5363: 100 nM, everolimus: 10 nM.

Fig.S17

T47D cells

A

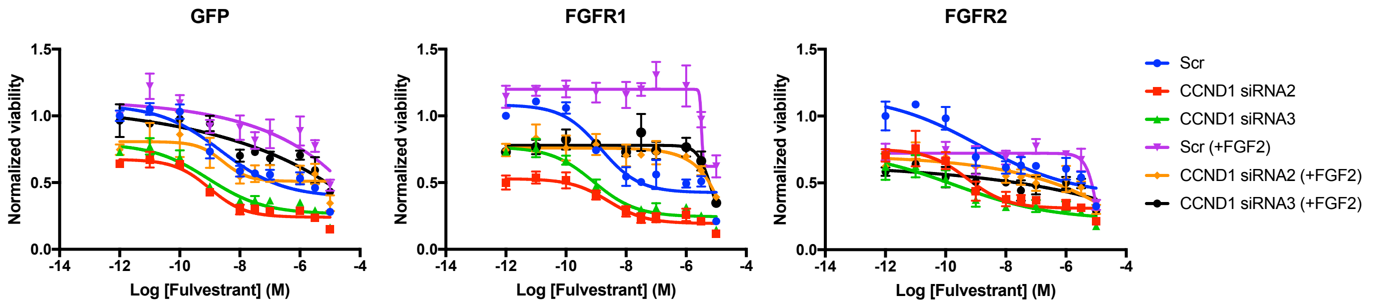


B

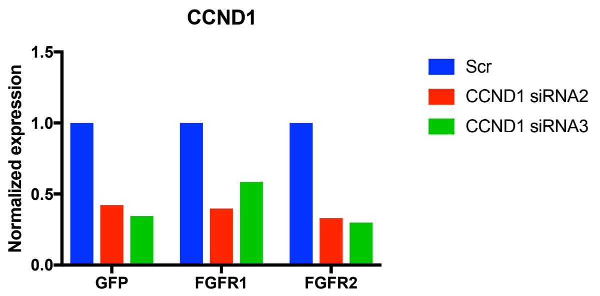


**Figure.S17 CCND1 knockdown reduced cell proliferation in T47D cells overexpressing FGFR1 or FGFR2.** A, CCND1 knockdown was achieved by two different siRNAs targeting CCND1 in T47D cells expressing GFP, FGFR1 and FGFR2. Cells were plated for drug treatment two days after siRNA transfection. Drug response was measured in the presence or absence of 10 ng/ µL FGF2 and 1 µM PD173074 after treatment for six days. B, CCND1 knockdown was confirmed by RT-PCR.

Fig.S18

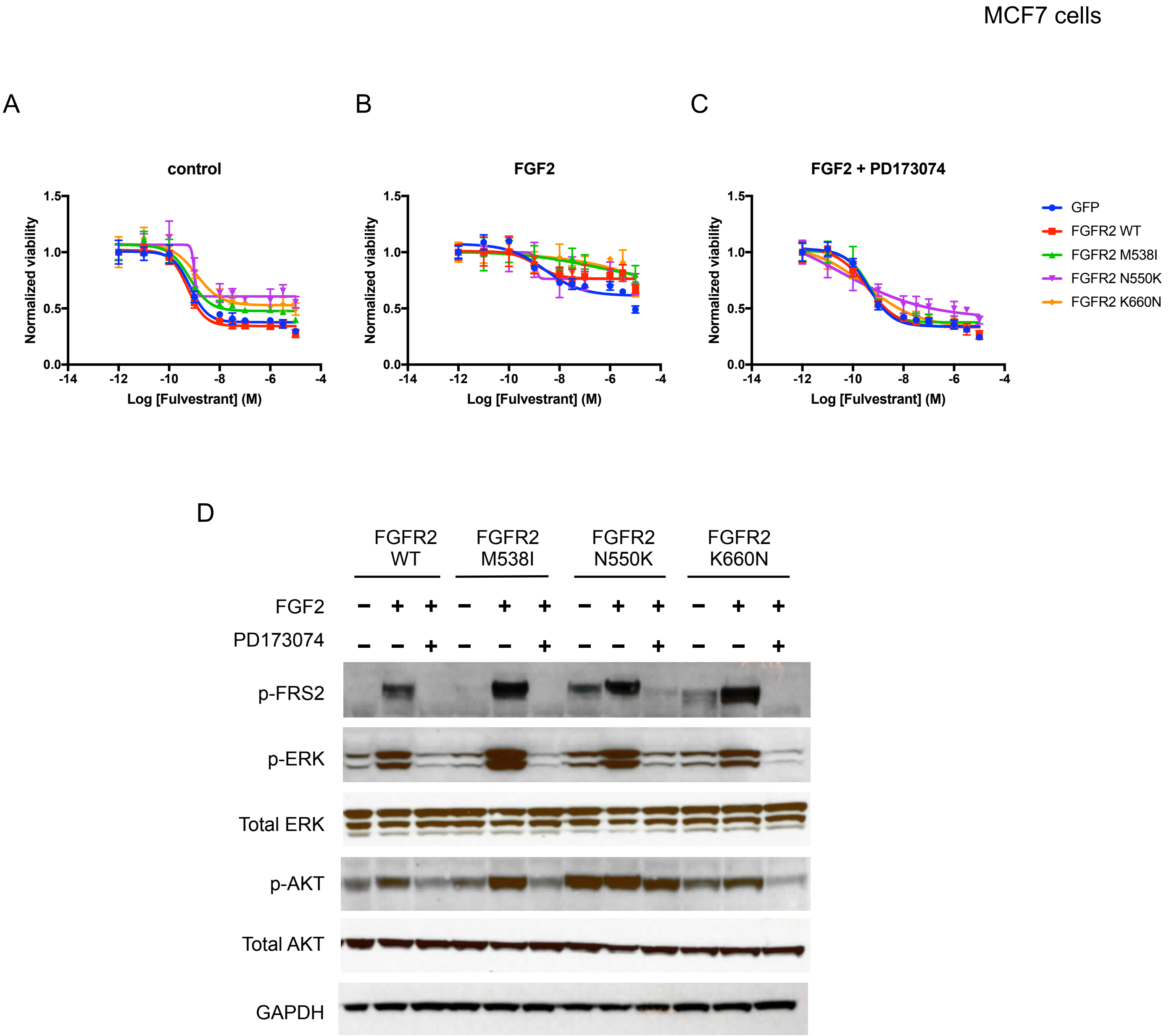


**Figure.S18 FGFR2 mutants conferred resistance to fulvestrant in MCF7 cells.** MCF7 cells overexpressing GFP, FGFR2 WT, FGFR2 M538I, FGFR2 N550K and FGFR2 K660N were established. Drug response to fulvestrant was examined in three conditions: control (A), 10 ng/mL FGF2 (B) and 10 ng/mL FGF2 with 1 μM PD173074 (C). D, FGFR2 mutants induced active downstream AKT and ERK signaling. MCF7 cell lines overexpressing different FGFR2 constructs were treated as indicated for one hour before protein harvest and western blot.

Fig.S19

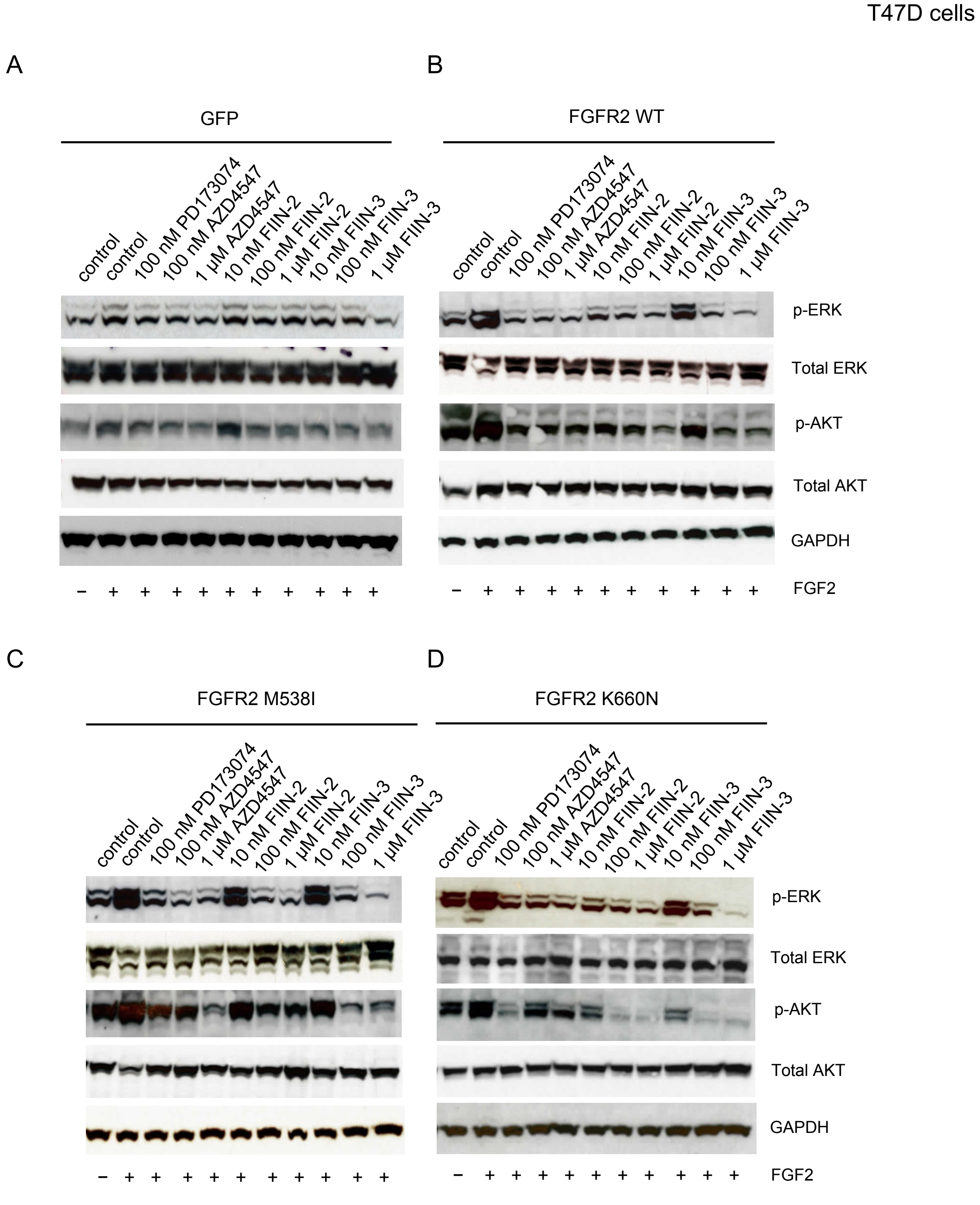


**Figure.S19 FIIN-2 and FIIN-3 are effective in targeting FGFR2 WT and FGFR2 mutants.** PD173074 and AZD4547 effectively blocked downstream AKT and ERK signaling of FGFR2 WT, FGFR2 M538I and FGFR2 K660N, although FIIN-2 and FIIN-3 exhibited higher potency. Cells overexpressing GFP (A), FGFR2 WT (B), FGFR2 M538I (C) and FGFR2 K660N (D) were treated as indicated for one hour before protein harvest and western blot.

Fig.S20

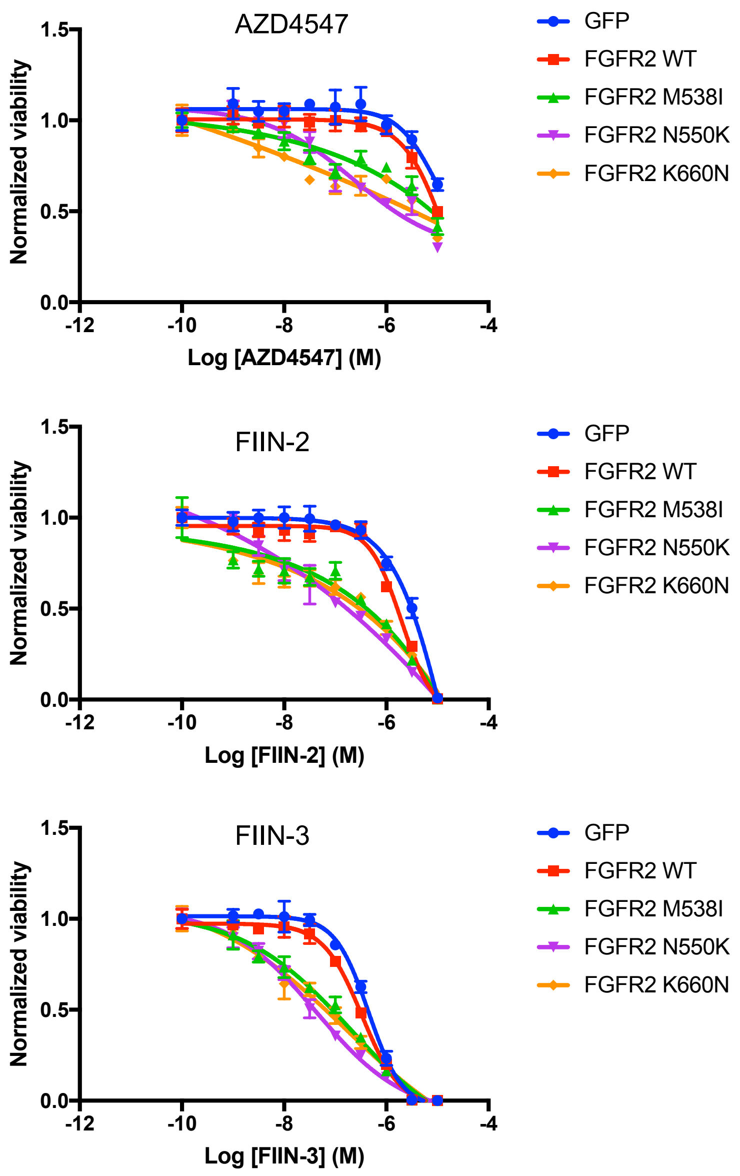


**Figure. S20 FGFR2 mutants induced hypersensitivity to FGFR inhibitors.** Stable cell lines overexpressing FGFR2 mutants (M538I, N550K and K660N) exhibited higher sensitivity to FGFR inhibitors AZD4547, FIIN-2 and FIIN-3, as compared to T47D cells overexpressing GFP or wildtype FGFR2.

Fig.S21

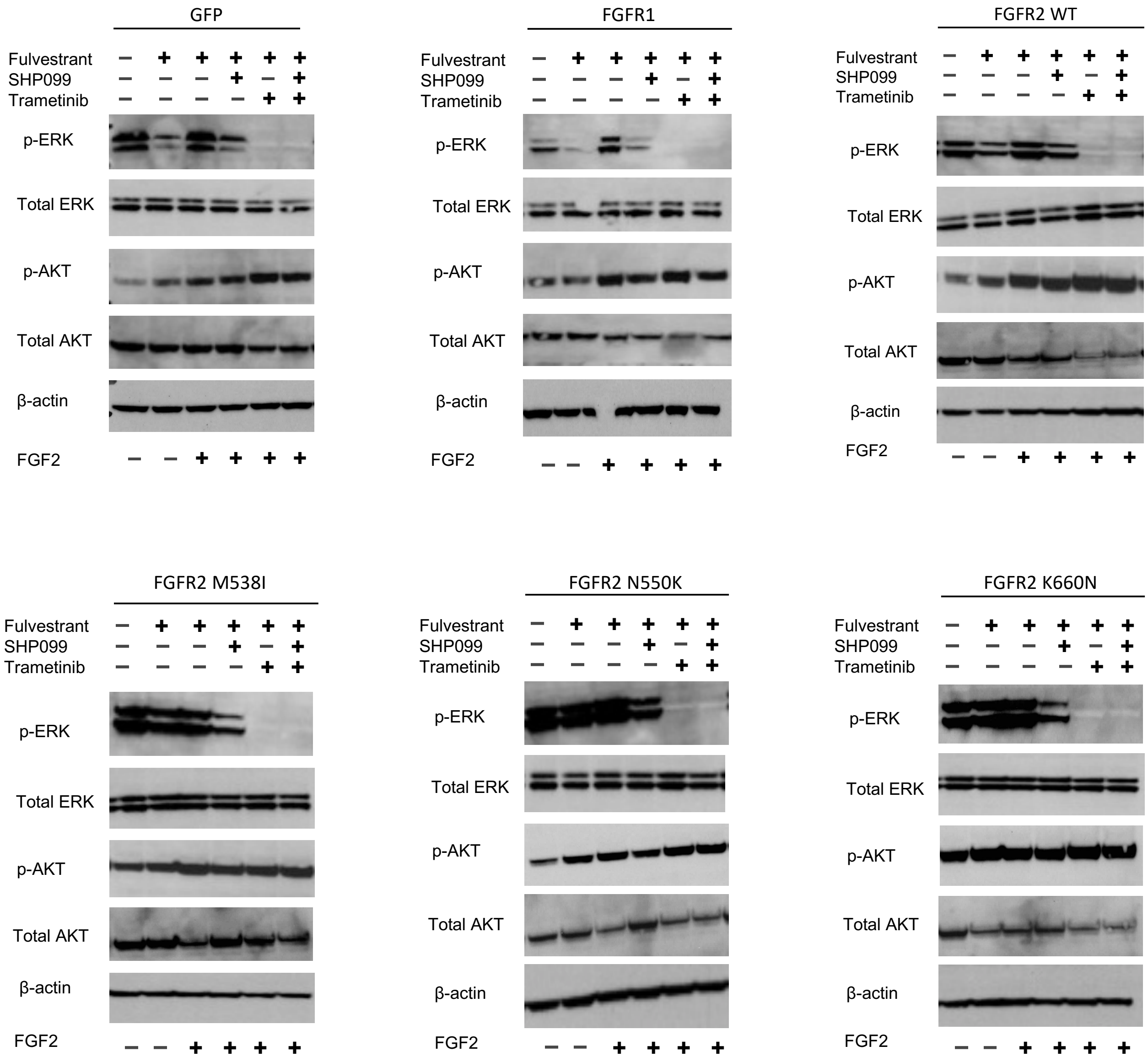


**Figure. S21 Downstream ERK signaling of FGFR1/2 was inhibited by SHP2 and/or MEK inhibitors.** SHP099 alone partially reduced ERK phosphorylation and the combination of SHP099 and trametinib completely abrogated p-ERK levels. Cells were treated as indicated for two days before protein harvest and western blot. Concentration of drugs used: 10 ng/mL FGF2; 1 μM fulvestrant; 500 nM trametinib; 10 μM SHP099; 500 nM FIIN-3.
